# Kin and group selection are both flawed but useful data analysis tools

**DOI:** 10.1101/742122

**Authors:** jeff smith, R. Fredrik Inglis

## Abstract

For understanding the evolution of social behavior in microbes, mathematical theory can aid empirical research but is often only used as a qualitative heuristic. How to properly formulate social evolution theory has also been contentious. Here we evaluate kin and multilevel selection theory as tools for analyzing microbial data. We reanalyze published datasets that share a common experimental design and evaluate these theories in terms of data visualization, statistical performance, biological interpretation, and quantitative comparison across systems. We find that the canonical formulations of both kin and multilevel selection are almost always poor analytical tools because they use statistical regressions that are poorly specified for the strong selection and nonadditive fitness effects common in microbial systems. Analyzing both individual and group fitness outcomes helps clarify the biology of selection. We also identify analytical practices in empirical research that suggest how theory might better handle the challenges of microbial data. A quantitative, data-driven approach thus shows how kin and multilevel selection theory both have substantial room for improvement as tools for understanding social evolution in all branches of life.

## Introduction

For biologists seeking to understand the evolution of cooperation and social behavior, mathematical theory has been a valuable guide for empirical research. Kin selection theory, for example, has been enormously influential in part because it shows that social evolution can be understood from just a few important quantities: how a behavior affects the fitness of an actor, how that behavior affects the fitness of other individuals, and the genetic relatedness between social partners [1, 2]. This theory was developed with a major focus on animal behavior, so its mathematical form was developed in terms of the quantitative-genetic phenotypes most easily measured in animal systems [3]. Similarly, evolutionarily stable strategy (ESS) models of kin selection were formulated to facilitate predictions and empirical tests using comparative data [4] with substantial success [5, 6, 7].

More recently, microbial systems have become popular for addressing social evolution [8]. Aside from these organisms’ intrinsic biological interest and importance as pathogens and mutualists, they allow for empirical methods that would be impractical or impossible in non-microbial systems. Fitness, for example, is notoriously difficult to measure in multicellular organisms [9]. In microbial systems, however, it is common practice to directly measure the fitness effect of a defined change in genotype or environment under controlled laboratory conditions. Evolution can be even be observed as it happens and replayed under different conditions [10, 11].

Microbial systems also have their challenges. It can be challenging to measure or even identify the phenotypes responsible for some fitness effect. And since meiotic sex is often infrequent or absent, studies of microbial genetics tend to focus on the effects of defined, discrete genotypes instead of quantitative genetic analyses of continuous-valued traits (but see [12]). It can be similarly difficult to study microbial behavior in natural environments, making comparative tests of ESS models rare. There is thus a disconnect between the terms in most mathematical formulations of social evolution theory and the types of data microbiologists most often collect. Perhaps as a result, researchers of microbial social evolution most often use *ad hoc* methods to analyze and interpret their data, employing theory as only a verbal heuristic or source of qualitative predictions.

Microbes are not intractable to mathematical theory, however. In the epidemiology of infectious disease, for example, there is extensive close interaction between models and data, with models formulated to make predictions in terms of the data empiricists collect [13, 14]. Models are tailored to the richness of available data and then used to make quantitative predictions. Deviation of data from these predictions is then used to refine models in an iterative cycle of improvement. One of the most important results to come out of this theory is a summary statistic called *R*_0_: the average number of secondary infections from a single infected individual in a wholly susceptible population [13, 14, 15]. *R*_0_ provides an estimate of outbreak population dynamics and pathogen fitness, allows researchers to compare very different infectious agents, and guides public health responses to diseases like SARS [16, 17] and Ebola [18].

What would a mathematical formulation of social evolution look like if it were able to closely engage with the quantitative details of typical microbial data? Is there a version of kin selection theory that would be better suited to microbes? Or would it be more promising to use another approach like multilevel selection, which examines how selection acts within and between groups of interacting individuals [19, 20]? The proper formulation of social evolution theory has been contentious [21, 22], but it has so far been difficult to evaluate the usefulness of different approaches as analytical tools because they have been debated mostly as abstract mathematics or as verbal interpretations of specific models and results.

Here we examine what happens where the proverbial rubber meets the road. We test how useful different theoretical approaches are for analyzing experimental data from social interactions among microbes. As a basis for comparison, we focus on a common experimental design we call a “mix experiment” which measures how microbial genotypes affect the performance of interacting individuals (Fig. 1A). We analyze datasets from published mix experiments using the canonical formulations of kin and multilevel selection theory and use quantitative measures of statistical performance to assess what these approaches do well, where they run into problems, and how often these problems occur. For guidance on how theory might better handle the challenges of microbial data, we also identify analytical practices in empirical research that are robust across different microbial systems, provide insight into the causes of selection, and allow quantitative comparison of social selection across systems. A quantitative, data-driven approach can thus be a productive way forward to identify how theory can best aid our understanding of social evolution in all branches of life.

**Figure 1:**
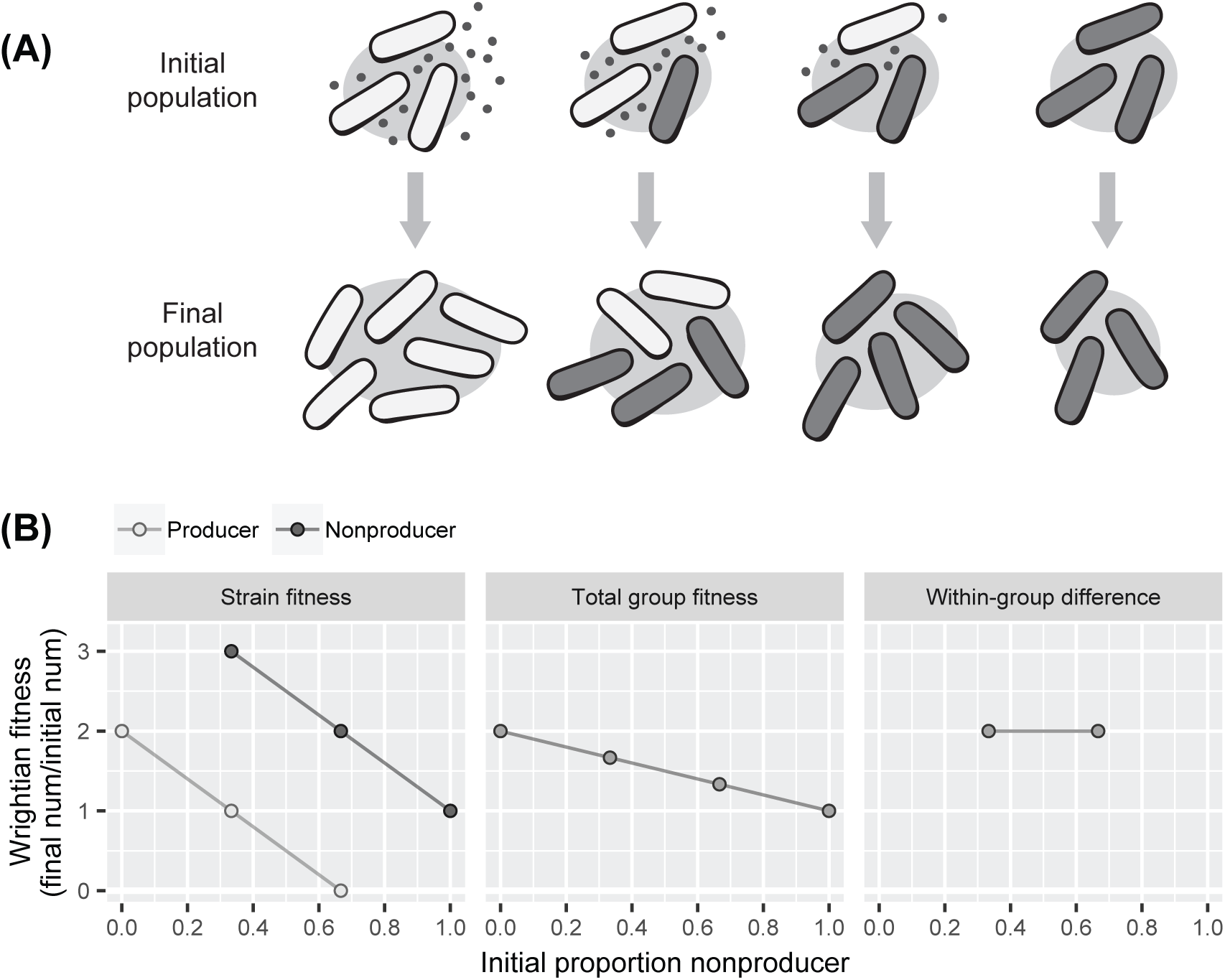
Kin and multilevel selection analysis of microbial mix experiments. **(A)** Diagram of typical mix experiment, measuring the performance of two strains as a function of their initial frequency in a population. Populations are created with a specified set of initial conditions and then some time later assayed to determine the abundance of each genotype. The time course of the experiment may include several microbial generations. Illustration shows classic “cheater” phenotype in which nonproducers of an external good outcompete producers within groups but decrease total group productivity. **(B)** Fitness models used by social evolution theory. Kin selection describes strain fitness as a multiple linear regression on its own genotype and that of its social neighbors. Multilevel selection describes total group fitness and the within-group difference between strains as linear regressions on mean group genotype. Both approaches use Wrightian fitness. Plots show analyses applied to hypothetical mix experiment shown in **(A)**. Lines show statistical models. Points show hypothetical data.

## Methods

### Choosing and processing datasets

We surveyed published literature for mix experiments with different genotypes of the same microbial species, focusing on studies of social evolution. We included experiments published in the last twenty years that measured the asexual survival and reproduction of strains as a function of their initial frequency, holding constant the total number of individuals. We aimed for breadth of species and social trait, prioritizing studies with several mixing frequencies (as opposed to a single 50:50 mix). We identified 39 studies that included experiments matching our criteria. We sought raw data from journal websites, public data archives, or directly from study authors. We reformatted datasets to express each experiment’s initial and final states as the number or density of individuals belonging to each strain, the proportion of individuals belonging to a strain, and/or the total number of individuals—whatever most closely matched the measured quantities. So that our results would not be overly dominated by a few studies with many mix experiments, we included a maximum 12 datasets per study, prioritizing those with most replication and covering the observed range of interactions. In all, we obtained 80 datasets from 20 different studies for further analysis (Table S1).

### Applying mathematical theory

Taking mix experiments as assays for the fitness effects in kin and multilevel selection theory, we let the initial and final populations represent one part of a life cycle expressed in discrete time. Table 1 summarizes our mathematical notation. Discrete-time social evolution theory uses Wrightian fitness (Eqn. S3). We let a strain’s absolute Wrightian fitness be *w* = *n*′*/n*, which measures the fold increase in absolute abundance of the strain over the course of an experiment. A genotype becomes more abundant if *w* > 1 and less abundant if *w* < 1. When convenient, we also analyzed Malthusian fitness *m* = ln *w*.

**Table 1:**
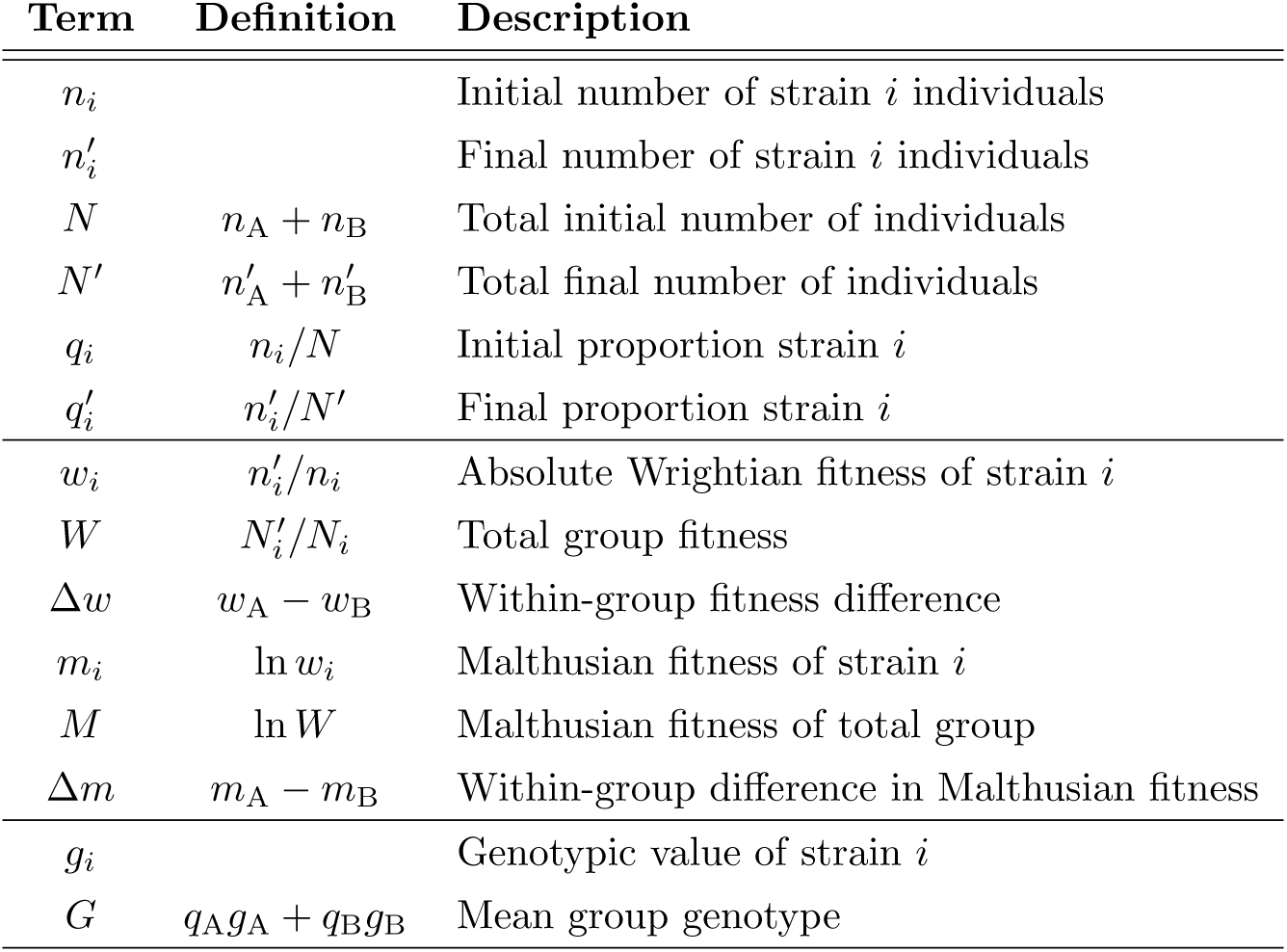
Mathematical notation and fitness measures

A common form of kin selection theory is neighbor-modulated fitness, which describes an individual’s fitness as a function of its own genotype and that of its social partners [23, 2]. If individual genotype is *g*, where *g* = 1 for the focal strain of interest and *g* = 0 for its partner, and mean group genotype is *G*, then kin selection describes fitness using a multiple least squares regression of the form *w* ∼ *g* + *G* (Fig. 1, Eqn. S6). The indirect fitness effect of an individual’s social partners (the “benefit” term in Hamilton’s rule) is the slope of the line fit to *w*(*G*). The direct fitness effect of an individual’s own genotype (the “cost” term in Hamilton’s rule) is the vertical offset between strains in the fitted model.

A common form of multilevel selection theory uses the Price equation to describe how selection acts within and between groups of interacting individuals [19, 20]. If total group fitness is *W* and the within-group fitness difference between strains is Δ*w*, then multilevel selection describes fitness using least squares regressions of the form *W* ∼ *G* and Δ*w* ∼ 1 (Fig. 1, Eqn. S9). The among-group selection gradient is the slope of the line fit to *W* (*G*). The within-group selection gradient is the expected value of Δ*w*.

### Calculations and statistics

We analyzed all data using the R environment for statistical computing [24]. Where applicable, we let wild-type and ancestral strains be the reference genotype and let mutants and laboratory-evolved derivatives be the focal genotype. We fit statistical models by least squares regression (`lm()` command), excluding zeroes for log-transformed quantities.

We evaluated the performance of statistical models using the Akaike Information Criterion (AIC) [25, 26], calculated via `AIC()`. AIC is a common model-selection tool that measures goodness of fit discounted by the number of parameters in the model. 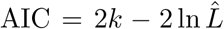, where *k* is the number of parameters in a model and 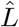 is maximum likelihood. AIC comparisons are meaningful even when models are not nested and have different predictors. A common rule of thumb is models perform substantially differently if |ΔAIC| > 2, equivalent to adding a parameter without improving goodness of fit. We sometimes compared the best-performing models (least AIC) from sets of candidates (Table S3). To compare models fit to linear or log-transformed fitness outcomes, we calculated the AIC of logarithmic models, as measured on the linear scale, as `AIC()` plus the Jacobian of the log transform: 2Σln *y*_*j*_, where the *y*_*j*_ are the fitted values [27].

We calculated unstandardized fitness effect sizes as *e*^*β*^, where *β* are the regression coefficients in “log-linear” models of the form *m* ∼ *g* + *G* + *gG, M* ∼ *G*, and Δ*m* ∼ *G*. In log-linear models, slopes are meaningfully comparable to intercepts because they measure the total effect of group genotype from zero to one. We also calculated scaled effect sizes of the form *β*′ = *β/Σ|β|*, where *β*′ ranges from zero to one and measures effect sizes as a proportion of all effects in a model. We information-theoretic measure analogous to effective species number in ecology as measured by calculated the effective number of parameters in log-linear models as 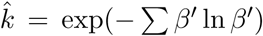, an Shannon diversity. 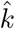 ranges from one when a single parameter explains all the data to the actual number of parameters (four in log-linear models) when all fitness effects have equal size.

To measure the performance of models fit to one set fitness outcomes at predicting the other set, we first fit models of the form *m* ∼ *g* + *G* + *gG, M* ∼ *G*, and Δ*m* ∼ *G* or ∼ ln(*G/*(1 − *G*)). We then used the parameters of these fitted models to calculate the predicted values for the other set of outcomes (Eqns. S11, S12). So that compared models would have the same error structure, we also fit a Gaussian error parameter via maximum likelihood (`mle()` command) given the data and predicted values. We counted *M* and Δ*m* as part of the same model so that neighbor-modulated and multilevel fitness models each used four parameters to explain the same dataset and ΔAIC measured relative goodness of fit 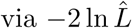.

## Results

To test how useful different theoretical approaches in social evolution are for analyzing data from microbial systems, we conducted kin and multilevel selection analyses of published datasets that share a common “mix experiment” design (Fig. 1A). We found that the canonical mathematical formulations of both approaches were almost always poor analytical tools because of strong, nonadditive fitness effects.

### Strong selection is common in microbial data

Kin and multilevel selection partition the components of fitness differently, but they both describe those partitions in terms of Wrightian fitness *w*—the number of individuals in the descendant population per individual in the ancestral population (Table 1, Eqn. S3). In microbial mix experiments, Wrightian fitness can range over several orders of magnitude (*e.g.* Fig. 2). As a result, the linear scale over which these theories describe social selection is a poor choice for visualizing data because many values cluster near zero. Statistical models fit on this scale also poorly characterize data in regions of low fitness and can predict nonsensical results like negative cell number.

**Figure 2:**
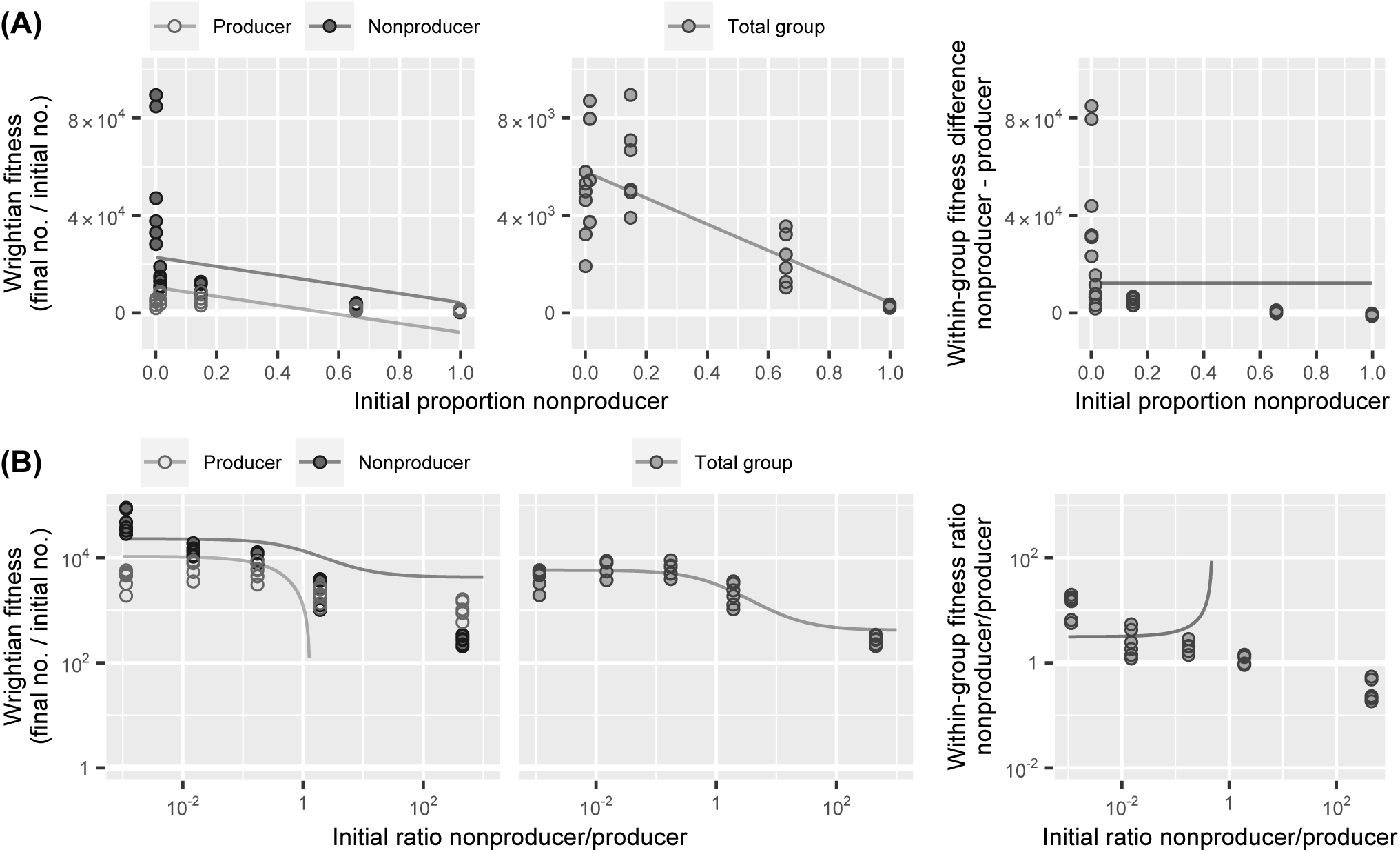
The canonical formulations of both kin and multilevel selection use statistical models that can be poor choices for analyzing microbial data because of strong selection and nonadditive fitness effects. **(A)** Additive fitness models fitted to data from mix experiment with *Pseudomonas aeruginosa* bacteria that do or do not produce a secreted iron-chelating siderophore [28]. **(B)** Data plotted over logarithmic scales to clearly visualize results. Linear models poorly characterize data in regions of low fitness and can predict nonsensical results like negative cell number. When social effects are nonadditive, strains can respond differently to mixing and within-group selection can be frequency-dependent. In all panels, points show experimental data and lines show fitted models. See Table 1 for mathematical notation.

To estimate how common strong fitness effects are in microbial systems, we measured the range of fitness values in published datasets (Fig. 3A). Over the range of mix frequencies investigated, strain and total group fitness typically spanned one or more orders of magnitude. Within groups, the fitness of one strain was often an order of magnitude or more greater than the other. Some systems had relatively weak fitness effects, but in most studies strong selection was the norm.

**Figure 3:**
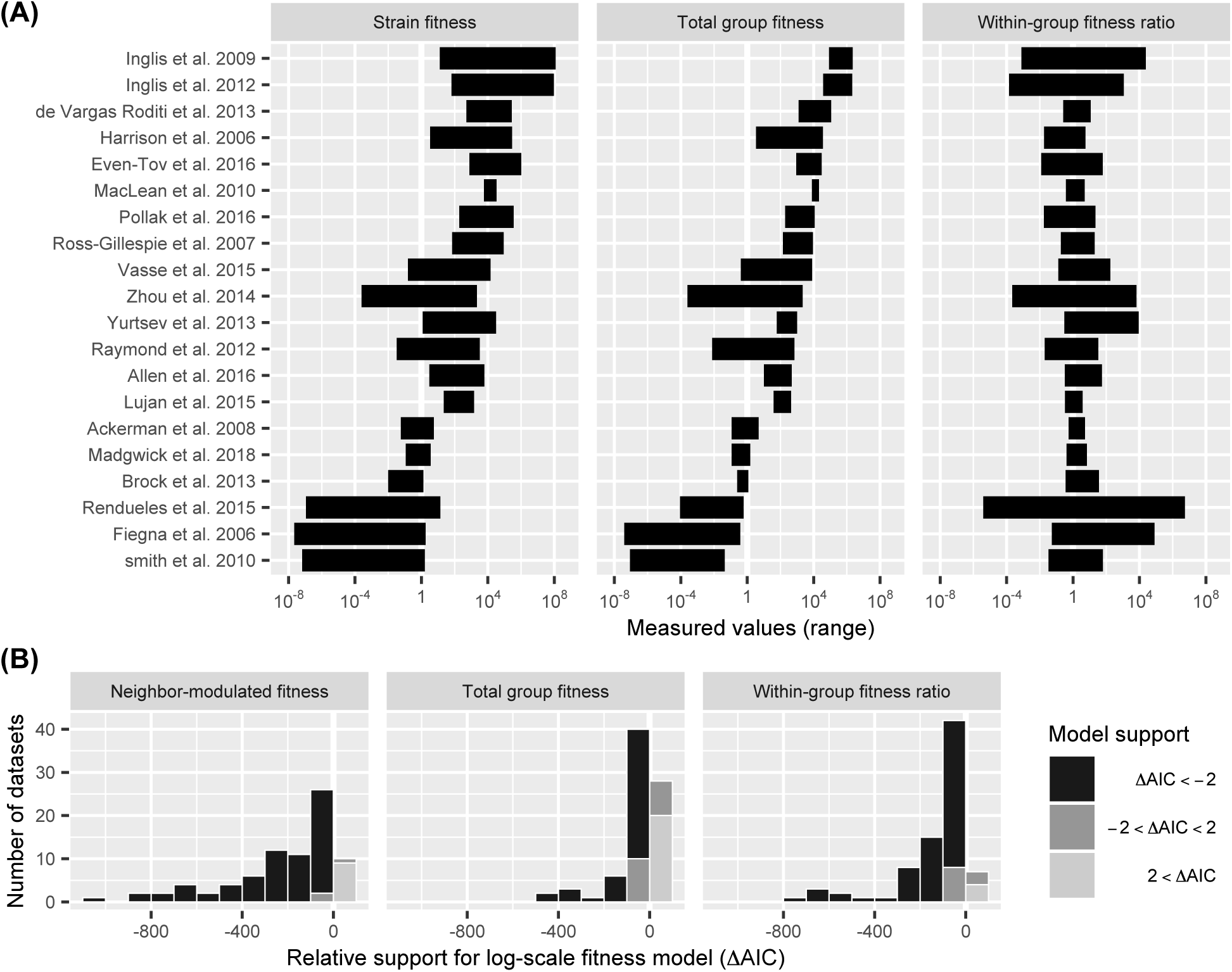
Strong selection is common in microbial mix experiments. **(A)** Wrightian fitness typically spans orders of magnitude. Data show range of measured values in datasets from analyzed studies, excluding zeroes. **(B)** The results of microbial mix experiments are usually best analyzed over logarithmic fitness scales. Data show AIC difference between statistical models fit to either log or linear fitness for the same dataset. Negative ΔAIC indicates better fit for log-fitness model. See Table S3 for model details.

For many of the datasets we examined, microbes underwent several rounds of reproduction. This can compound fitness effects across generations and increase their measured magnitude. But in some datasets with large effects, populations actually decreased in abundance (sporulating *Myxococcus* bacteria, for example). And many datasets concerned cell death, survival under starvation, or cooperative increase in local carrying capacity, so there was often no meaningful sense in which fitness effects could be measured “per generation” or expressed as weak effects on net growth rate.

Many of the studies we examined dealt with strong selection by analyzing experimental outcomes over logarithmic scales (Table S2) to visualize data and fit statistical models over the entire relevant range. To quantify how much better fitness is analyzed over logarithmic rather than linear scales, we compared the performance of statistical models fit to either linear or log-transformed outcomes but with the same predictor and error structure. Log-scale fitness models outperformed linear models for nearly all of the datasets and fitness outcomes we analyzed (Fig. 3B). One drawback to log-scale fitness is that transformed data are undefined when zero or negative. Zero fitness sometimes occured when one strain became undetectable in the final population. In many studies, the fitness of a strain was also sometimes greater and sometimes less than its partner depending on mixing frequency (Fig. 3A), such that the within-group fitness difference Δ*w* was sometimes negative. To analyze frequency-dependent selection over logarithmic scales, then, many studies instead used a within-group fitness measure equal to the fitness ratio *w*_A_*/w*_B_ or the Malthusian fitness difference Δ*m* (Table S2).

### Nonadditive fitness effects are common in microbial data

Another issue with analyzing microbial mix experiments using kin or multilevel selection theory is that these approaches, in their canonical formulations, both fit additive fitness models to data that are often not additive. Fitness effects are additive if they independently and linearly increase Wrightian fitness. Weak effects are always approximately additive. In their canonical formulations, kin and multilevel selection each use two additive parameters to represent fitness effects. Kin selection’s indirect fitness effect of social neighbors corresponds to a statistical model where strain fitness is a linear function of mix frequency (Fig. 1). Multilevel selection’s between-group fitness effect corresponds to a model where whole-group fitness is also a linear function of mix frequency. Kin selection’s direct fitness effect and multilevel selection’s within-group fitness effect both correspond to models where the within-group fitness difference between strains is constant.

In the microbial datasets we analyzed, all these terms were often strongly nonadditive (*e.g.* Fig. 2). Both strain and total group fitness often responded nonlinearly to mixing frequency. Within-group fitness differences between strains also depended on mixing frequency. As a result, the statistical models fitted by the kin and multilevel selection approaches were frequently poor summaries of the data and predicted nonsensical results like negative cell number.

To quantify the extent of nonadditivity, we compared the statistical performance of additive fitness models with those that included nonadditive terms (Fig. 4A). For nearly all datasets, nonadditive models usually described neighbor-modulated and within-group fitness better than additive models, and for total group fitness performed at least as well. There were no datasets for which additive models performed better. Among the microbial datasets we analyzed, fitness nonadditivity was the norm.

**Figure 4:**
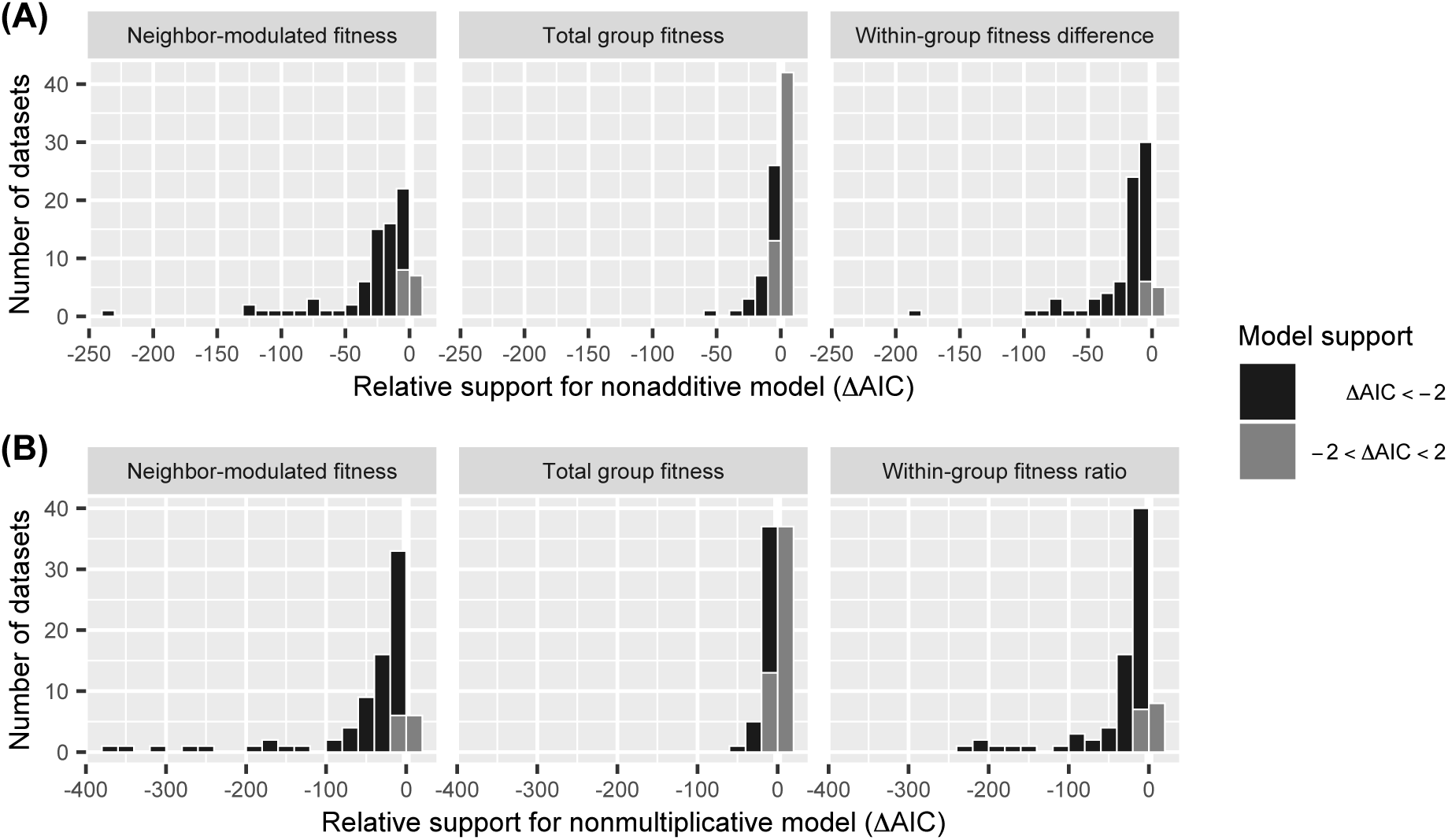
Nonadditive fitness effects are ubiquitous in microbial data. **(A)** Over linear fitness scales, models usually perform better when they allow nonadditive effects on strain fitness and within-group fitness. **(B)** Even over logarithmic fitness scales, models usually perform better when they allow nonmultiplicative effects. Data show AIC difference between models for the same dataset. Negative ΔAIC indicates better fit. See Table S3 for model details.

Strong selection can create fitness effects that are multiplicative rather than additive. To assess how much this contributed to nonadditivity, we compared multiplicative fitness models to those that could include interaction and nonlinear terms. The more-elaborate models usually outperformed the simpler multiplicative models (Fig. 4B). There were no datasets for which multiplicative models performed better. The two fitness predictors used in standard kin and multilevel selection regression were thus insufficient to capture the fitness effects seen in these microbial datasets. Model performance was particularly improved by allowing frequency-dependent selection within groups and interactions between individual and neighbor genotype (Fig. S1A). Allowing quadratic nonlinearities improved model performance more modestly (Fig. S1B).

One of the consequences of nonadditivity is that the strongest fitness effects frequently occured at extreme mixing frequencies, on the order of a percentage or less (*e.g.* Fig. 2B). For visualizing and analyzing fitness effects at extreme mixing frequencies, several studies used a log-ratio measure of frequency like ln(*q*_A_*/q*_B_). To quantitatively evaluate the usefulness of examining data over logarithmic mixing frequencies, we compared the performance of fitness models with the same predictor structure but with either mixing proportion or log mixing ratio as the independent variable (Fig. S2). We found that within-group fitness benefitted most from this approach, with the within-group fitness ratio commonly being a log-log function of mixing ratio (e.g. Fig. 2B). Total group fitness, on the other hand, often responded more linearly to mixing proportion. In some cases, strain and total group fitness showed strong effects at extreme mixing frequencies but were not log-linear (e.g. Fig. S28).

### Strain and group-focused fitness outcomes are both useful

Kin and multilevel selection focus on different fitness outcomes. For microbial mix experiments, kin selection focuses on strain fitness, while multilevel selection focuses on total group fitness and the relative fitness of strains within groups (Fig. 1). Each set of outcomes can be calculated from the other (Eqns. S11, S12). So we asked: what are the consequences of analyzing a particular set of outcomes?

In the datasets we examined, we found cases where one set of outcomes revealed some biological result that the other did not. Analyzing data from multiple perspectives also revealed similarities between systems that would not have been visible otherwise. During sporulation of *Myxococcus* bacteria, for example, evolved strain PX inhibits sporulation of its ancestral wild-type but is itself unaffected by mixing (Fig. 5A). Researchers were aware of this one-way mixing effect (G. J. Velicer, *pers. comm.*), but because they only reported multilevel fitness outcomes it is not apparent in the original study [29], nor is its similarity to later findings in *Dictyostelium* amoebae (Fig. S15) [30]. To quantify the extent to which effects seen in one set of fitness outcomes were apparent in the other set, we fit neighbor-modulated and mutlilevel fitness models to each dataset and then compared analogous effect sizes (Fig. 5B). Overall, strain and multilevel fitness outcomes mostly told similar stories, with the notable exception that mixing frequently affected strain fitness much more than total group fitness. We found similar results when comparing effect sizes scaled as a fraction of the total so that the relationships would be less dominated by extreme values (Fig. S3, S7B). Strain and total group fitness are only expected to respond similarly to mixing when within-group selection and the fitness effects of individual genotype are small relative to the effects of group and neighbor genotype (Eqns. S17, S19). In many datasets, however, there was strong selection within groups but little change in total group fitness, a situation known as soft selection [31, 32, 33]. Soft selection can happen when total population size is determined by some limited resource over which strains compete—glucose in a flask of media, for example.

**Figure 5:**
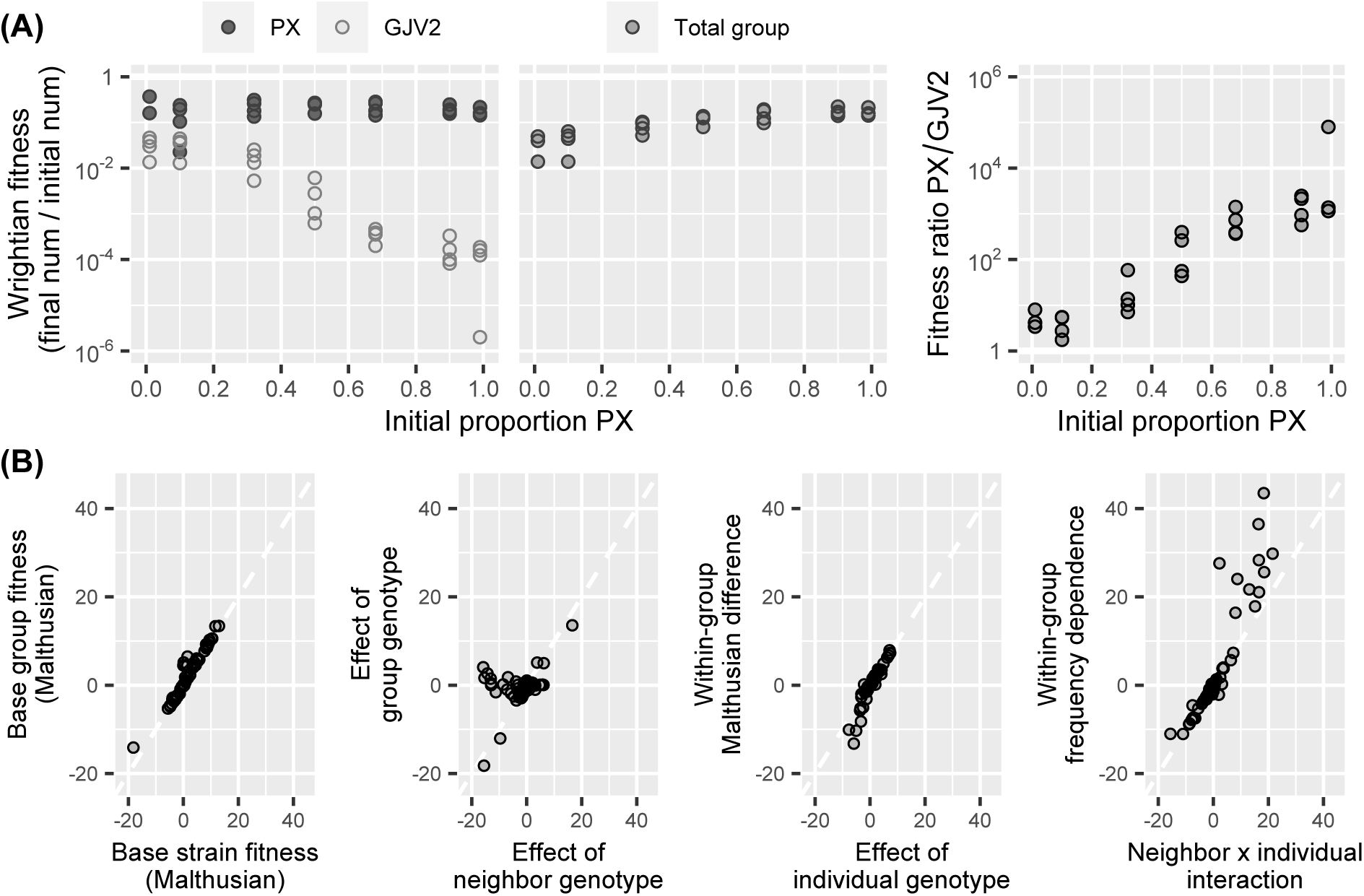
Analyzing both strain and group-focused fitness outcomes can reveal results that would otherwise go unseen. **(A)** Example from *Myxococcus* bacteria. Mixing reduces the fitness of wild-type strain GJV2 but has no effect on lab-evolved strain PX. This one-way effect is not apparent in the original study [29] because it only reports multilevel fitness outcomes. **(B)** Across datasets, kin and multilevel selection analyses mostly tell similar stories, but mixing frequently affects strain and total group fitness differently. Data compare analogous Malthusian effect sizes in statistical models fit to the same dataset.

Some fitness outcomes might be more convenient to analyze because they are more linear over logarithmic scales. If strain fitness is a log-linear function of mixing proportion, for example, then total group fitness is log-nonlinear, and vice versa (Eqns. S17, S19). To address this issue, we examined how well log-linear models fit to one set of fitness outcomes predicted the other set. We found that the best-performing model was sometimes neighbor-modulated fitness and sometimes multilevel fitness, depending on the dataset (Fig. S4).

A theoretical framework might capture microbial fitness effects more succinctly, with fewer meaningful parameters, if a fitness outcome is near-constant in one framework but there are no constant outcomes in the other framework, like in the one-way mixing example above. To address this issue, we fit comparable neighbor-modulated and multilevel fitness models to data then measured how evenly distributed fitness effects were distributed among model parameters (Figs. S6, S7). Multilevel selection consistently described fitness effects with fewer effective parameters (paired *t*(79) = 6.23, *P* < 0.001), in part because of widespread soft selection. Multilevel selection models can capture pure soft selection with a single within-group term, while kin selection models identify a further effect of neighbor genotype [31, 33].

Strain and multilevel fitness outcomes visualize data differently. Strain outcomes can be presented in a single compact plot but can also be hard to parse when strains have similar fitness (*e.g.* Fig. 2). Multilevel fitness outcomes require separate plots but have fewer issues with overlapping data. Strain fitness naturally illustrates the relative magnitudes of direct and indirect fitness effects, which is also possible with multilevel outcomes but requires deliberately matched axes. Within-group selection is only defined for mixed-genotype groups (not strains by themselves), while strain and total-group fitness are defined for both single-strain and mixed-genotype groups.

Strain and multilevel fitness outcomes were both effective for quantitatively comparing social selection in different datasets. The fitness impact of social interaction varied considerably among systems (Fig. 6, S5). In datasets concerning multicellular sporulation of *Myxococcus* bacteria or *Dictyostelium* amoebae, for example, most fitness effects were social. The effects of shared external goods like siderophores or secreted enzymes, on the other hand, were often minor compared to base strain or group fitness. In most datasets there were interactions between neighbor and individual genotype and frequency-dependent selection within groups (Fig. S7). These nonadditive fitness effects were not just minor complications—they were typically a substantial proportion of all social effects and could be larger than base within-group fitness or the direct effect of individual genotype.

**Figure 6:**
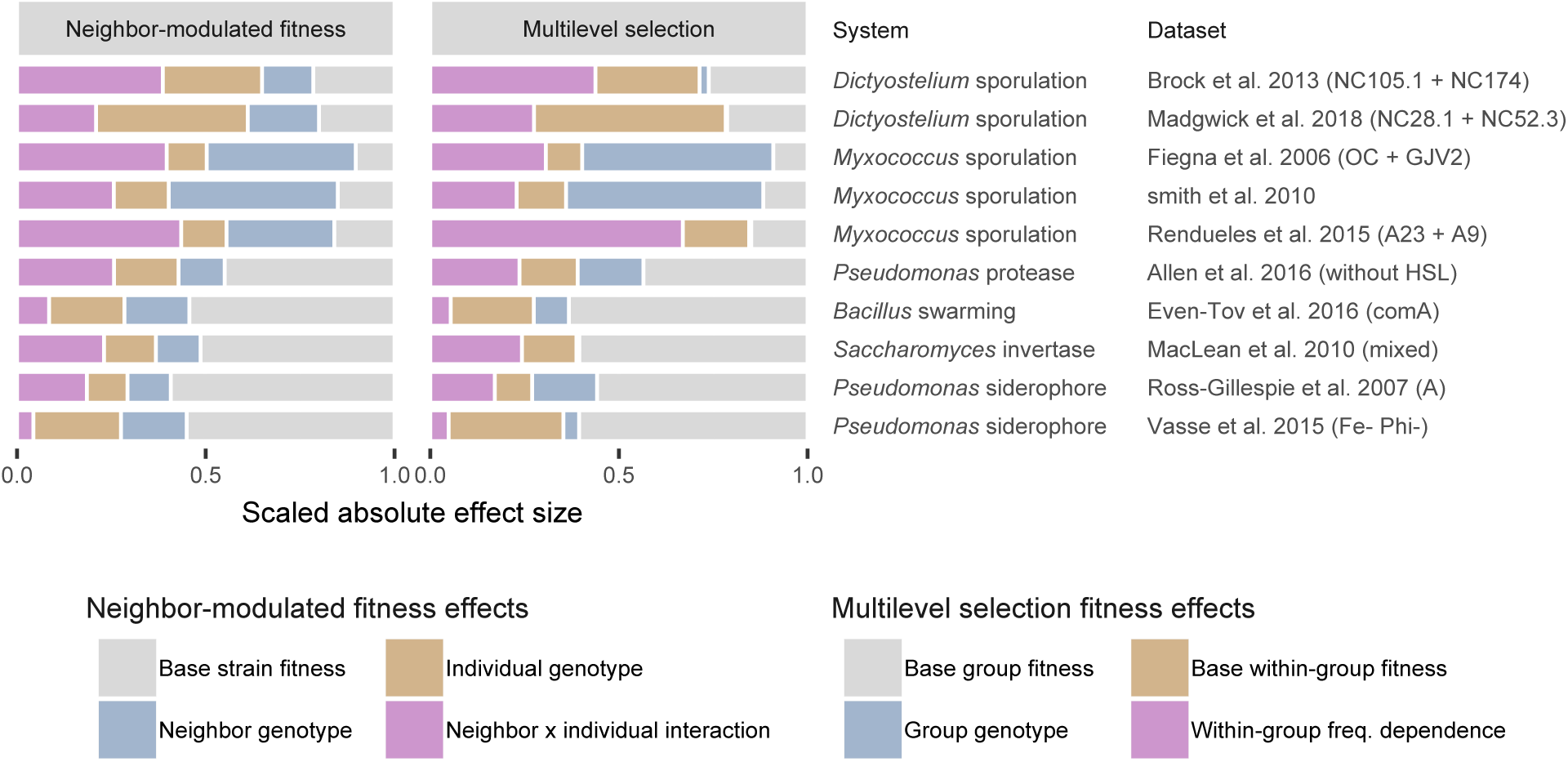
Social interactions have more impact in some systems than in others. In many *Dictyostelium* and *Myxococcus* datasets, for example, most fitness effects are social. In other datasets, the largest effect is base fitness. Data show Malthusian effect sizes for select datasets, scaled as a fraction of all observed effects. Colors indicate analogous effects. See also Fig. S5.

A theoretical approach might be more intuitive if its terms more directly correspond to the raw data collected by researchers. For the datasets we examined, the experimentally measured quantities were most often total and within-group performance, though strain performance was also common (Table S2, Fig. S8) and the difference was nonsignificant (Fisher exact test *P* = 0.63). Some experimental designs measured all three outcomes simultaneously (counting different colony types on the same plate, for example). Studies that directly measured a fitness outcome were not more likely to report it (Fisher exact test: strain *P* = 1, total group *P* = 0.54, within-group *P* = 0.088). Instead, studies reported relative within-group performance most frequently and strain performance the least (Fisher exact test *P* = 0.015). Strikingly, even studies framed verbally in terms of kin selection sometimes only reported group-focused fitness outcomes [28, 34, 35, 36]. Multilevel selection analyses thus appear to hold substantial intuitive appeal, and there is scope for more extensive use of strain fitness data.

Together, these results indicate it is empirically useful to analyze both strain and group-focused fitness outcomes, then use whichever are most convenient or informative for the data and biological question of interest.

## Discussion

### Disconnect between analytical needs and existing theory

To evaluate the usefulness of social evolution mathematics as tools for analyzing microbial data, we used kin selection and multilevel selection theory to analyze data from published studies that share a common “mix experiment” design. We found that both theoretical approaches, in their standard formulations, are poor analytical tools because they use statistical models that are poor characterizations of microbial data. These models are unable to adequately describe strong selection or nonadditive fitness effects, both of which were ubiquitous in the microbial datasets we analyzed. The fitness terms in the canonical versions of kin and multilevel selection are population regressions, made over individuals and groups in the population they are meant to describe. When the statistical models in these regressions do not match the actual relationships in the data, the measured regression coefficients will not be true for populations with different value distributions— different genotype frequencies, for example, or a different distribution of genotypes among groups— even if experiments exactly replicate natural conditions in all other ways [37, 38, 39]. The fitness terms in these poorly-specified regressions are therefore not very useful summary statistics with which to quantify and compare social selection in microbial systems.

The issues we found were caused by the specific statistical models being fitted, not by the fitness outcomes emphasized under either theoretical framework. Examining each genotype’s fitness separately (as in kin selection theory) or examining fitness within and between groups (as in multilevel selection theory) are both useful approaches for analyzing microbial data. When we re-analyzed data from published experiments, we found that using different fitness outcomes revealed results that would not be otherwise visible. Which analytical approach is most useful will depend on the specific question being investigated and may not be apparent beforehand. So while the proper formulation of social evolution theory is sometimes contentious [21, 22], it is practically useful for empiricists to examine their data from multiple theoretical perspectives [40]. An analogy might be how physics embraces different representations of the same process if they are more convenient or provide deeper insight—different coordinates systems, for example, or classical versus Lagrangian mechanics—even when they make the same predictions.

Microbial fitness effects may be more amenable to analysis with canonical kin and multilevel selection theory when performance outcomes are measured for individual cells or virions, or over shorter timescales, where fitness effects are expected to be weaker. Advances in single-cell microscopy and other measures of expression and secretion by microbial individuals make this approach promising [41, 42]. But it seems unlikely that the fitness consequences of social interactions among microbes will always be expressible as weak additive effects on growth and survival rate. The primary benefit of shared external goods like iron-chelating siderophores, for example, may be to increase local carrying capacity. And there may be no meaningful measure of instantaneous fitness during elaborately coordinated processes like fruiting body formation in *Myxococcus* or *Dictyostelium*.

For many research questions, general social evolution theory may not be “the right tool for the job”. General theory is best used for comparing different systems on the same terms and deriving general principles (*e.g.* [43]). For answering specific biological questions, on the other hand, it is often best to use models tailored to biology of a system [44, 45]. Biologically-specific models can predict the magnitude and frequency dependence of fitness outcomes in mix experiments, for example, and explain the sources of nonadditivity [46, 47, 48, 33]. Following the time dynamics of mix experiments can be especially informative, as the specificity and richness of time-series data allows strong tests of hypotheses that would otherwise be indistinguishable based on the initial and final time points alone [33].

### Reformulating social evolution theory: challenges and prospects

If we wanted social evolution theory to be a more useful tool for quantitative analysis of microbial data, what does it need to do? Kin and multilevel selection are both statistical descriptions of social selection. Like statistical mechanics in physics, they describe the behavior of aggregate properties of a system (allele frequency, genotype fitness, *etc.*) in terms of the probabilistic behavior of its individual parts. So we can phrase the theoretical challenge here in statistical terms: How should we specify the statistical model describing social selection? What is the appropriate functional form of the model, and what variables should it include?

The main problem with the canonical formulations of kin and multilevel selection is their quantitative genetic approach based on the Price Equation and linear least-squares regression— appropriate for macroscopic phenotypes like sex ratio, perhaps, but not for microbial fitness. At present, practically-minded microbiologists simply construct statistical models with fitness scales, functional responses, and error distributions that reflect their actual data (Poisson regression or generalized linear models, for example). But then interpreting what these ad hoc analyses tell us about social evolution in microbes is a verbal exercise in which theory is just a heuristic. The theoretical challenge is to formulate mathematical expressions of social selection that can handle the complexities of real microbial fitness data while also enabling quantitative inferences about selection’s net effect in a structured population with nonrandom interactions among genotypes. Key to this challenge are strong selection and non-additive fitness effects. Analytically useful formulations of social selection in microbes would be able to generically allow fitness effects that span orders of magnitude and have non-linear, genotype-dependent responses to genotype frequency among social interactants. They would also be expressible in terms of either strain or multilevel fitness outcomes as needed, be robust to different types of biological interaction, and allow quantitative comparisons across very different systems.

One approach is to extend existing theory, adding terms to Hamilton’s rule for example to better handle nonadditive fitness effects [49, 37]. This kind of elaborative approach still requires ad hoc data analyses, though, which then need to be translated into the terminology of Wrightian fitness. Another approach is to reformulate theory in terms more easily applied to and more meaningful for microbial data. For this approach, a promising guide might be the analytical practices used by experimental researchers. Most prominent and ubiquitous of these is modelling fitness over logarithmic scales, either as transformed values like log10 *w* or as Malthusian fitness *m*. Within-group relative fitness can be analyzed over logarithmic scales when expressed as a fitness ratio *w*_A_*/w*_B_ or as a difference in Malthusian fitnesses Δ*m*. Modelling genotype frequency over logarithmic scales is also useful, since the fitness effects of social neighbors can happen at extreme frequencies. Fisher [50] advocated using log-frequency ratios like ln(*q*_*A*_*/q*_*B*_) because they allow one to easily distinguish genotype frequencies even when one type is very rare, respond linearly to simple forms of selection, behave symmetrically around zero whether a genotype is rare or common, and have a simple statistical interpretation as the log-odds of sampling one genotype relative to another. One challenge for logarithmic-scale analyses, though, are the zeroes that occur in single-strain groups and when a genotype becomes undetectible in the final population, both of which occured in the datasets we analyzed.

Theory can be especially useful for interpreting what nonadditivity means biologically. What biological processes can generate the functional form seen in the data? Curve shape can be a rich source of data with which to make strong tests of scientific hypotheses [51], but for social selection there is relatively little theory to draw upon. Some cases have a straight-forward interpretation, like the one-way mixing effects discussed above, but others are not as obvious. Multiple levels of specificity are possible, from quantitative models tailored to the biology and parameter values of specific systems [46, 48, 33] to heuristic models of general biological processes that make simple qualitative predictions [52, 28]. A useful conceptual guide to different types of nonadditivity might lie in evolutionary game theory [53, 54]. In this theoretical framework, genotypes interact to create fitness payoffs whose structure defines the game being played: “prisoner’s dilemma”, “snowdrift”, *etc*. There are versions of game theory that can deal with nonadditive payoffs and games with many simultaneous players, though combining these features with structured populations remains an area of theoretical development. Ideally, theory would provide not only a set of tools and models but also a concise, general, quantitative statement about evolution under social selection.

## Conclusions

Mathematical theory can play many roles in biology. It can be a conceptual or heuristic guide. It can be a proof of concept to test the verbal logic of hypotheses [55]. It can also be an analytical tool that provides quantitative insight into the biology of a system—epidemiology’s *R*_0_, for example. Here we have evaluated how kin and multilevel selection theory perform in this last analytical role. The two theories emphasize different sets of fitness outcomes, and we found that analyzing both sets can help clarify the biology of selection. But the canonical versions of both theories use statistical models poorly suited to the strong selection and nonadditivity frequently seen in microbial data, severely limiting their usefulness as tools for quantitative analysis and interpretation.

Both kin and multilevel selection have always been envisioned as extremely general processes in evolution. But while some of their early conceptual development explicitly discussed microbial systems [56, 57], their mathematical formulation and empirical study have mostly focused on metazoa. Applying theory developed for one set of taxa to another very different set can act as a test of that theory’s explanatory power. By this measure, both kin and multilevel selection have room for improvement. It is useful to have theory that speaks in same terms as the data, which is why mathematical formulations of social evolution theory have been tailored to the biology and methods of “organismal” biology—quantitative genetics for the study of measurable phenotypes with weak selective effects and unknown genetic causes [3, 58], and ESS models for comparative studies and tests of facultatively adaptive behavior [4]. But microbial science often deals with very different methodologies and data—defined genotypic changes that have large effects on growth and survival under experimentally defined conditions, for example. Fitness is notoriously difficult to measure in metazoa and plants [9], but for microbes it is one of the most common forms of experimental data. Microbial phenotypes, in fact, can be more difficult to measure. Perhaps it is not surprising that theory developed to understand biological phenomena from one set of taxa loses much of its usefulness when applied to another set with very different biology.

We are not claiming that kin or multilevel selection theory are conceptually incorrect, nor that they make incorrect assumptions. It is still possible to fit their canonical fitness models to data from microbial mix experiments and correctly interpret the parameters of those models in terms of average effects in the studied population [2, 39]. But because the fitness measures and statistical models are poorly specified for microbial data, those average effects are not terribly useful—they are not expected to be the same for other populations with the same biology, nor even for the same population at another time [37, 38, 39]. It is no surprise that empirical microbiologists use other fitness measures and other statistical models to analyze and interpret their data.

There is still plenty of room to develop mathematical formulations of social evolution theory that better reflect the practical realities of microbial science. We have identified some of the complexities in microbial data that would need to be addressed and have highlighted some of the promising analytical practices used in empirical research. Taking a quantitative, data-driven approach to theory can help us move beyond verbal and conceptual arguments to find a productive path where both kin and multilevel selection are useful analytical tools for understanding social evolution in all branches of life.

## Acknowledgements

Many thanks to R. Allen, D. Brock, S. Brown, S. Diggle, L. de Vargas Roditi, A. Eldar, J. Gore, Gudelj, W.-D. Hardt, A. Jousset, F. Harrison, R. Kümmerli, A. Luján, B. Raymond, O. Rendueles, A. Ross-Gillespie, M. Vasse, G. J. Velicer, J. Wolf, and E. Yurtsev for generously sharing data. Thanks to A. Traulsen for discussion and comments on the manuscript. js and RFI were supported by startup funds from the University of Missouri–St. Louis. This material is partly based upon work supported by the National Science Foundation under grant nos. DEB1204352, IOS1256416, and DEB1146375 to J. E. Strassmann and D. C. Queller (Washington Univ. in St. Louis).

## Author contributions

Conceived project: js. Designed analyses: js, RFI. Collected data from published studies: js, RFI. Analyzed data: js. Interpreted results and organized their presentation: js, RFI. Wrote manuscript: js, RFI.

