## Supplemental Material for "Kin and group selection are both flawed but useful data analysis tools"

September 30, 2019

**Table S1:** Analyzed datasets

| **Study** | **Species** | **Trait*** | **Datasets available** | **Datasets used** |
|---|---|---|---|---|
| Ackermann *et al.* 2008 [1] | *Salmonella* Typhimurium | Virulence factor | 1 | 1 |
| Allen *et al.* 2016 [2] | *Pseudomonas aeruginosa* | Quorum sensing | 2 | 2 |
| Brock *et al.* 2013 [3] | *Dictyostelium discoideum* | Multicellular sporulation | 8 | 8 |
| de Vargas Roditi *et al.* 2013 [4] | *Pseudomonas aeruginosa* | Swarming | 2 | 2 |
| Even-Tov *et al.* 2016) [5] | *Bacillus subtilis* | Quorum sensing | 2 | 2 |
| Fiegna *et al.* 2003 [6] | *Myxococcus xanthus* | Multicellular sporulation | 3 | 3 |
| Harrison *et al.* 2006 [7] | *Pseudomonas aeruginosa* | Siderophore | 1 | 1 |
| Inglis *et al.* 2009 [8] | *Pseudomonas aeruginosa* | Bacteriocin | 1 | 1 |
| Inglis *et al.* 2012 [9] | *Pseudomonas aeruginosa* | Bacteriocin/Siderophore | 1 | 1 |
| Luján *et al.* 2015 [10] | *Pseudomonas fluorescens* | Siderophore | 2 | 2 |
| MacLean *et al.* 2010 [11] | *Saccharomyces cerevisiae* | Secreted invertase | 2 | 2 |
| Madgwick *et al.* 2018 [12] | *Dictyostelium discoideum* | Multicellular sporulation | 34 | 12 |
| Pollak *et al.* 2016 [13] | *Bacillus subtilis* | Quorum sensing | 2 | 2 |
| Raymond *et al.* 2012 [14] | *Bacillus thuringiensis* | Virulence factor | 1 | 1 |
| Rendueles *et al.* 2015 [15] | *Myxococcus xanthus* | Multicellular sporulation | 14 | 12 |
| Ross-Gillespie *et al.* 2007 [16] | *Pesudomonas aeruginosa* | Siderophore | 3 | 3 |
| smith *et al.* 2010 [17] | *Myxococcus xanthus* | Multicellular sporulation | 1 | 1 |
| Vasse *et al.* 2015 [18] | *Pesudomonas aeruginosa* | Siderophore | 4 | 4 |
| Yurtsev *et al.* 2013 [19] | *Escherichia coli* | Antibiotic resistance | >50 | 12 |
| Zhou *et al.* 2014 [20] | *Bacillus thuringiensis* | Quorum sensing | 8 | 8 |

* Quorum sensing can control multiple downstream traits.

**Table S2:** Fitness outcomes in original studies

| | Measured outcome | | | Reported outcome* | | |
|---|---|---|---|---|---|---|
| **Study** | **Strain** | **Total group** | **Within group** | **Strain** | **Total group** | **Within group** |
| Ackermann *et al.* 2008 | ✓ | ✓ | - | $\lg n'$ | - | - |
| Allen *et al.* 2016 | - | ✓ | ✓ | $\ln(w-1)$ | $N'$ | $\Delta m$ |
| Brock *et al.* 2013 | ✓ | ✓ | ✓ | $w$ | - | - |
| de Vargas Roditi *et al.* 2013 | ✓ | ✓ | ✓ | - | $N'$ † | $q'-q$ |
| Even-Tov *et al.* 2016 | - | ✓ | ✓ | - | - | $w_A/w_B$ † |
| Fiegna *et al.* 2006 | ✓ | - | - | - | - | $\lg(w_A/w_B)$ |
| Harrison *et al.* 2006 | ✓ | ✓ | ✓ | - | $M/\ln 2$ | $m_A/m_B$ |
| Inglis *et al.* 2009 | ✓ | ✓ | ✓ | - | - | $\Delta m/m_B$ |
| Inglis *et al.* 2012 | ✓ | ✓ | ✓ | - | $N'$ | $\Delta m/m_B$ |
| Luján *et al.* 2015 | ✓ | ✓ | ✓ | - | $N'$ † | $w_A/w_B$ |
| MacLean *et al.* 2010 | - | ✓ | ✓ | - | $\lg N'/\lg N'_{max}$ | $m_A/m_B$ |
| Madgwick *et al.* 2018 | - | ✓ | ✓ | $(q\,n'_{alone})/n'$ | $\overline{n'_{alone}}/N'$ | $w_A/w_B$ |
| Pollak *et al.* 2016 | - | ✓ | ✓ | - | $N'$ | $w_A/w_B$ † |
| Raymond *et al.* 2012 | ✓ | ✓ | ✓ | $\lg w$ | $\lg N'$ | $w_A/w_B$ † |
| Rendueles *et al.* 2015 | ✓ | - | - | $\lg(w/w_{alone})$ | $\lg(N'/\overline{n'_{alone}})$ | $\lg(w_A/w_B)$ |
| Ross-Gillespie *et al.* 2007 | ✓ | ✓ | ✓ | - | $N'$ † | $w_A/w_B$ † |
| smith *et al.* 2010 | ✓ | - | - | $\lg w$ | - | - |
| Vasse *et al.* 2015 | ✓ | ✓ | ✓ | - | - | $\lg(w_A/w_B)$ |
| Yurtsev *et al.* 2015 | - | ✓ | ✓ | - | - | $q'$ |
| Zhou *et al.* 2014 | ✓ | ✓ | ✓ | $\lg n'$ | $\lg N'$ | $w_A/w_B$ † |

* See Table 1 for mathematical notation. lg = $\log_{10}$. $x_{alone}$ = value in single-strain experiments. Studies may report different but mathematically equivalent calculation. Reported outcomes included if applied to mixed-strain experiments. Studies may not report all outcomes for all experiments.
† Reported on logarithmic scale

### Supplemental mathematics

#### Kin and multilevel selection analysis of microbial mix experiments

Kin and multilevel selection are theoretical frameworks for understanding natural selection when individuals interact. They describe evolutionary change in different terms. Here we briefly outline their mathematical formulations to show how one would use microbial mix experiments as assays for their fitness terms.

Contemporary formulations of both kin and multilevel selection are based on the Price equation [21, 22]. The Price equation describes evolution using two terms, one relating fitness to genotype and one describing how genotypes are transmitted across generations. The discrete-time version of the Price equation is

$$\overline{w}\Delta\overline{g} = \mathrm{Cov}(w, g) + \mathrm{E}(w\Delta g) \tag{S1}$$

where $\overline{w}$ is population-mean fitness, $\Delta\overline{g}$ is the change in mean genotype between the initial and final population, $\Delta g$ is the change in mean genotype between an individual and its offspring, and the covariance and expectation are taken across all individuals in the initial population. The Price equation is not a model—it is a mathematical identity that holds for definitions of fitness that satisfy

$$q' = q\frac{w}{\overline{w}}\,. \tag{S2}$$

We can rewrite Eqn. S2 as

$$\begin{aligned}\frac{n'}{N'} &= \frac{qw}{\sum qw} \\ &= \frac{wn/N}{\sum wn/N}\end{aligned}$$

and

$$\frac{n'}{\sum n'} = \frac{wn}{\sum wn}\,, \tag{S3}$$

which makes clear that $w = n'/n$ (absolute Wrightian fitness). The Price equation is no longer true, for example, if you substitute $w$ with Malthusian fitness $m = \ln(n'/n)$.

In the absence of mutation, infectious gene transfer, or other processes that affect the genotype of offspring, $\mathrm{E}(w\Delta g) = 0$. We can then rewrite Eqn. S1 to isolate the fitness terms:

$$\Delta\overline{g} = \beta_{wg}\mathrm{Var}(g)/\overline{w}\,, \tag{S4}$$

where $\beta_{wg}$ is the coefficient of a least-squares regression of $w$ on $g$ over all individuals in the population. Eqn. S4 is an expression of evolution due to selection commonly used in quantitative genetics.

The most commonly used form of kin selection analysis is neighbor-modulated fitness, which describes how an individual's fitness is affected by its own genotype and that of its social partners [23, 24]. In its most-general formulation, it expands the total regression of fitness on genotype into a multiple regression on individual genotype $g$ and mean neighbor genotype $G$:

$$\beta_{wg} = \beta_{wg|G} + \beta_{wG|g}\,\beta_{Gg}\,. \tag{S5}$$

$\beta_{wg|G}$ is the partial regression coefficient of fitness on individual genotype, holding neighbor genotype constant. It measures the direct effect of an individual's genotype on its own fitness. $\beta_{wG|g}$ is the partial regression coefficient of fitness on neighbor genotype, holding individual genotype constant. It measures the indirect effect of neighbor genotype on an individual's fitness. $\beta_{Gg}$ is the regression coefficient of neighbor genotype on individual genotype. It measures the statistical similarity of interacting individuals relative to the whole population. To obtain Hamilton's rule for the evolution of cooperation [25, 24], let the direct fitness effect of a cooperative genotype be its cost $-C = \beta_{wg|G}$, the indirect fitness effect be its benefit $B = \beta_{wG|g}$, and the regression coefficient of neighbor genotype on individual genotype be Hamiltonian relatedness $R = \beta_{Gg}$. Cooperative genotypes increase in frequency when the total regression of fitness on genotype is positive: $RB - C > 0$

For microbial mix experiments, then, the basic quantity of interest in a neighbor-modulated fitness analysis is a strain's fitness $w$ as a function of its own genotype $g$ and the average genotype of its local social group $G$, expressed as a statistical model of the form

$$w \sim g + G\,. \tag{S6}$$

For the large groups assayed in most microbial experiments, it is easiest and most straight-forward to use the "whole-group" formulation of neighbor-modulated fitness that includes one's self in its social environment [26]. The alternative would be an "other-only" formulation, but since groups are so large in most microbial experiments they are effectively the same.

Multilevel selection describes how selection acts within and between groups. A common version of multilevel selection uses the Price equation to expand itself hierarchically, partitioning selection into one part due to group-mean fitness and another part due to individual within-group fitness [27, 28]. The expanded equation can be written

$$\overline{w}\Delta\overline{g} = \mathrm{Cov}_j(W, G) + \mathrm{E}_j\left[\,\mathrm{Cov}_{i|j}(w, g)\,\right], \tag{S7}$$

where $i$ and $j$ subscripts indicate where covariances and expectations are taken over individuals and groups, respectively. Note that

$$\begin{aligned} W &= \sum qw \\ &= \sum \frac{n}{N}\frac{n'}{n} \\ &= \sum n'/N \\ &= N'/N\,, \end{aligned}$$

so group-mean fitness is equal to total group fitness. Rewriting Eqn. S7 to isolate the fitness terms,

$$\overline{w}\Delta\overline{g} = \beta_{WG}\,\mathrm{Var}(G) + \mathrm{E}_j\left[\,\beta_{wg|j}\mathrm{Var}_{i|j}(g)\,\right]. \tag{S8}$$

Multilevel selection describes how genotypes are distributed among groups using variance terms. Eqn. S8 partitions fitness into two components: a least-squares regression of group-mean fitness on group-mean genotype with coefficient $\beta_{WG}$ and a least-squares regression of individual fitness on individual genotype with coefficient $\beta_{wg|j}$. In mix experiments with only two genotypes $g_{\mathrm{A}} = 1$ and $g_{\mathrm{B}} = 0$, the within-group regression term can be written $\beta_{wg|j} = (w_A - w_B)/(g_{\mathrm{A}} - g_{\mathrm{B}}) = \Delta w$. The basic quantities of interest in a multilevel selection analysis of microbial mix experiments are thus $W$ and $\Delta w$, represented as statistical models of the form

$$\begin{aligned} W &\sim G \\ \Delta w &\sim 1\,. \end{aligned} \tag{S9}$$

Contextual analysis is another kind of multilevel selection analysis [29]. It is also a quantitative-genetics approach, based on a multiple least-squares regression of fitness on individual and group traits. Contextual analysis does not uniquely specify what traits are relevant, but when the individual trait of interest is individual genotype and the group trait of interest is group-mean genotype, it uses the same fitness model as neighbor-modulated fitness.

#### Strain and group fitness outcomes

Kin and multilevel selection focus on different fitness outcomes. Neighbor-modulated fitness uses one outcome ($w$) as function of two independent variables ($g, G$) while multilevel fitness uses two outcomes ($W, \Delta w$) that are a function of one independent variable ($G$). For microbial mix experiments, it's convenient to write strain fitness as two outcomes ($w_\mathrm{A}, w_\mathrm{B}$), each a function of one independent variable ($G$). With strong selection, it's also convenient to analyze Malthusian fitness outcomes. So we have two outcomes for neighbor-modulated fitness ($m_\mathrm{A}, m_\mathrm{B}$) and two for multilevel fitness ($M, \Delta m$).

To obtain ($M, \Delta m$) given ($m_\mathrm{A}, m_\mathrm{B}$), expand total group fitness as

$$\begin{aligned} M &= \ln W \\ &= \ln(G w_\mathrm{A} + (1-G) w_\mathrm{B})) \\ &= \ln(G e^{m_\mathrm{A}} + (1-G) e^{m_\mathrm{B}})\,. \end{aligned} \tag{S10}$$

We can thus calculate multilevel fitness from strain fitness as

$$\begin{aligned} M &= \ln[G e^{m_\mathrm{A}} + (1-G) e^{m_\mathrm{B}}] \\ \Delta m &= m_\mathrm{A} - m_\mathrm{B}\,. \end{aligned} \tag{S11}$$

To obtain ($m_\mathrm{A}, m_\mathrm{B}$) given ($M, \Delta m$), substitute $m_\mathrm{A} = m_\mathrm{B} + \Delta m$ into Eqn. S11 so that

$$\begin{aligned} M &= \ln\big(G e^{m_\mathrm{B}+\Delta m} + (1-G) e^{m_\mathrm{B}}\big) \\ &= \ln\big(G e^{m_\mathrm{B}} e^{\Delta m} + (1-G) e^{m_\mathrm{B}}\big) \\ &= \ln\big(e^{m_\mathrm{B}}(1 - G + G e^{\Delta m})\big) \\ &= \ln e^{m_\mathrm{B}} + \ln\big(1 - G + G e^{\Delta m}\big) \\ &= m_\mathrm{B} + \ln\big(1 - G + G e^{\Delta m}\big)\ . \end{aligned}$$

We can thus calculate strain fitness from multilevel fitness as

$$\begin{aligned} m_\mathrm{A} &= M - \ln(1 - G + G e^{\Delta m}) + \Delta m \\ m_\mathrm{B} &= M - \ln(1 - G + G e^{\Delta m})\,. \end{aligned} \tag{S12}$$

If neighbor-modulated fitness is a log-linear function of mixing proportion we can write its functional form as

$$m_i = m_0 + \beta_1 G + \beta_2 g_i + \beta_3 G g_i \,. \tag{S13}$$

With $g_{\mathrm{A}} = 1$ and $g_{\mathrm{B}} = 0$, strain fitness is

$$\begin{aligned} m_{\mathrm{A}} &= m_0 + \beta_2 + (\beta_1 + \beta_3)G \\ m_{\mathrm{B}} &= m_0 + \beta_1 G \end{aligned} \tag{S14}$$

and

$$\Delta m = m_{\mathrm{A}} - m_{\mathrm{B}} = \beta_2 + \beta_3 G \,. \tag{S15}$$

So if strain fitness is a log-linear function of $G$ then the within-group fitness ratio will be, too. Rearranging Eqn. S12, we can write total group fitness as

$$M = m_{\mathrm{B}} + \ln(1 - G + Ge^{\Delta m}) \tag{S16}$$

$$= m_0 + \beta_1 G + \ln(1 - G + Ge^{\Delta m}) \tag{S17}$$

which is in general not a linear function of $G$. The $\ln(1 - G + Ge^{\Delta m})$ term ranges from zero as $G \to 0$ to $\Delta m$ as $G \to 1$. So total group fitness is most similar to strain fitness (and thus most log-linear) when the within-group fitness difference is small relative to the effect of group genotype on total group fitness.

Conversely, if total group fitness is a log-linear function of mixing frequency we can write its functional form as

$$M = M_0 + \beta_4 G \,. \tag{S18}$$

Substituting Eqn. S18 into Eqn. S12,

$$\begin{aligned} m_{\mathrm{A}} &= M_0 + \beta_4 G - \ln(1 - G + Ge^{\Delta m}) + \Delta m \\ m_{\mathrm{B}} &= M_0 + \beta_4 G - \ln(1 - G + Ge^{\Delta m}) \,. \end{aligned} \tag{S19}$$

So if multilevel fitness outcomes are log-linear functions of $G$ then strain fitness outcomes are log-nonlinear. Strain fitness approaches log-linearity when the fitness effect of individual genotype is small relative to the effect of neighbor genotype.

**Table S3:** Model comparisons

| Comparison | Reference model(s)* | Focal model(s)** |
|---|---|---|
| Log-scale fitness (Fig. 3B) | $w \sim g + G$<br>$\sim g + G + gG$<br>$\sim g + G + gG + G(1-G)$<br>$\sim g + G + gG + G(1-G) + gG(1-G)$<br>$W \sim G$<br>$\sim G + G(1-G)$<br>$w_\mathrm{A}/w_\mathrm{B} \sim 1$<br>$\sim G$<br>$\sim G + G(1-G)$ | $m \sim g + G$<br>$\sim g + G + gG$<br>$\sim g + G + gG + G(1-G)$<br>$\sim g + G + gG + G(1-G) + gG(1-G)$<br>$M \sim G$<br>$\sim G + G(1-G)$<br>$\Delta m \sim 1$<br>$\sim G$<br>$\sim G + G(1-G)$ |
| Nonadditivity (Fig. 4A) | $w \sim g + G$<br>$W \sim G$<br>$\Delta w \sim 1$ | $w \sim g + G + gG$<br>$\sim g + G + gG + G(1-G)$<br>$\sim g + G + gG + G(1-G) + gG(1-G)$<br>$W \sim G + G(1-G)$<br>$\Delta w \sim G$<br>$\sim G + G(1-G)$ |
| Nonmultiplicativity (Fig. 4B) | $m \sim g + G$<br>$M \sim G$<br>$\Delta m \sim 1$ | $m \sim g + G + gG$<br>$\sim g + G + gG + G(1-G)$<br>$\sim g + G + gG + G(1-G) + gG(1-G)$<br>$M \sim G + G(1-G)$<br>$\Delta m \sim G$<br>$\sim G + G(1-G)$ |
| Linear nonmultiplicativity (Fig. S1A) | $m \sim g + G$<br>$\Delta m \sim 1$ | $m \sim g + G + gG$<br>$\Delta m \sim G$ |
| Quadratic nonmultiplicativity (Fig. S1B) | $m \sim g + G + gG$<br>$M \sim G$<br>$\Delta m \sim G$ | $m \sim g + G + gG + G(1-G)$<br>$\sim g + G + gG + G(1-G) + gG(1-G)$<br>$M \sim G + G(1-G)$<br>$\Delta m \sim G + G(1-G)$ |
| Log mixing ratio (Fig. S2) | $m \sim g + G + gG$<br>$M \sim G$<br>$\Delta m \sim G$ | $m \sim g + \operatorname{logit} G + g \operatorname{logit} G$<br>$M \sim \operatorname{logit} G$<br>$\Delta m \sim \operatorname{logit} G$ |

* See Table 1 for mathematical notation. Where multiple models indicated for an outcome, comparison made using model with least AIC (per dataset).
** $\operatorname{logit} G = \ln(G/(1-G))$

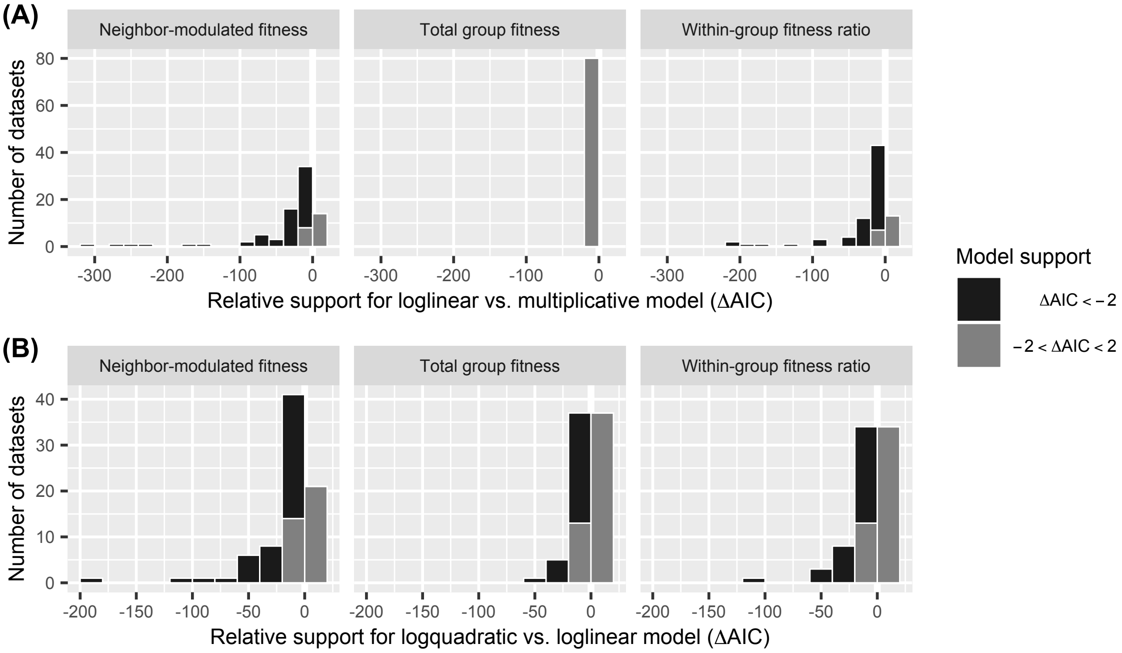

**Figure S1:** **(A)** Fitness models usually perform better when they allow frequency-dependent selection within groups and interactions between individual and neighbor genotype. For total group fitness, multiplicative and loglinear models are identical. **(B)** Allowing Malthusian fitness outcomes to be quadratic functions of mixing proportion can also improve model performance. Data show AIC difference between models for each dataset. Negative ΔAIC indicates better fit. See Table S3 for model details.

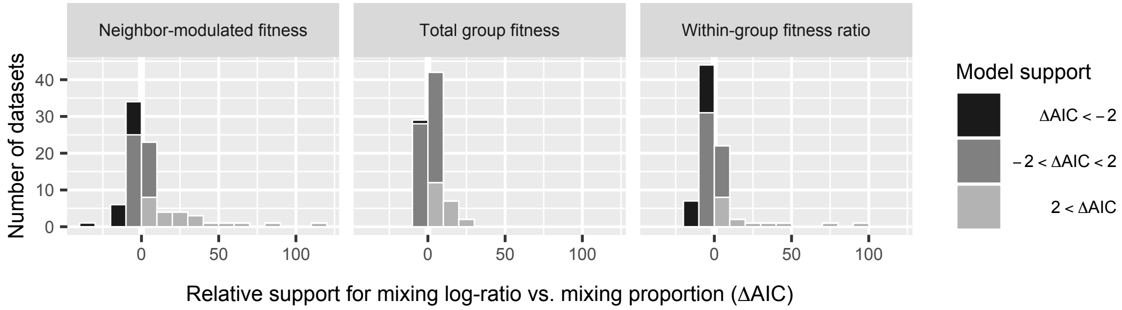

**Figure S2:** Fitness models sometimes perform better as functions of log mixing ratio $\ln(G/(1-G))$ instead of mixing proportion $G$. Data show AIC difference between models for each dataset. Negative ΔAIC indicates better fit for log-ratio model. See Table S3 for model details.

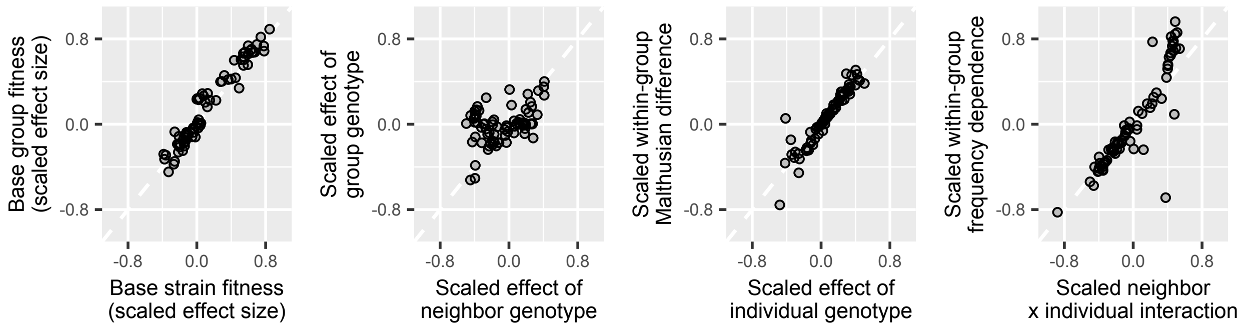

**Figure S3:** Across datasets, kin and multilevel selection analyses mostly tell similar stories, but mixing frequently affects strain and total group fitness differently. Data show scaled Malthusian effect sizes in neighbor-modulated and and multilevel fitness models fit to the same dataset. Plots compare analogous fitness effects.

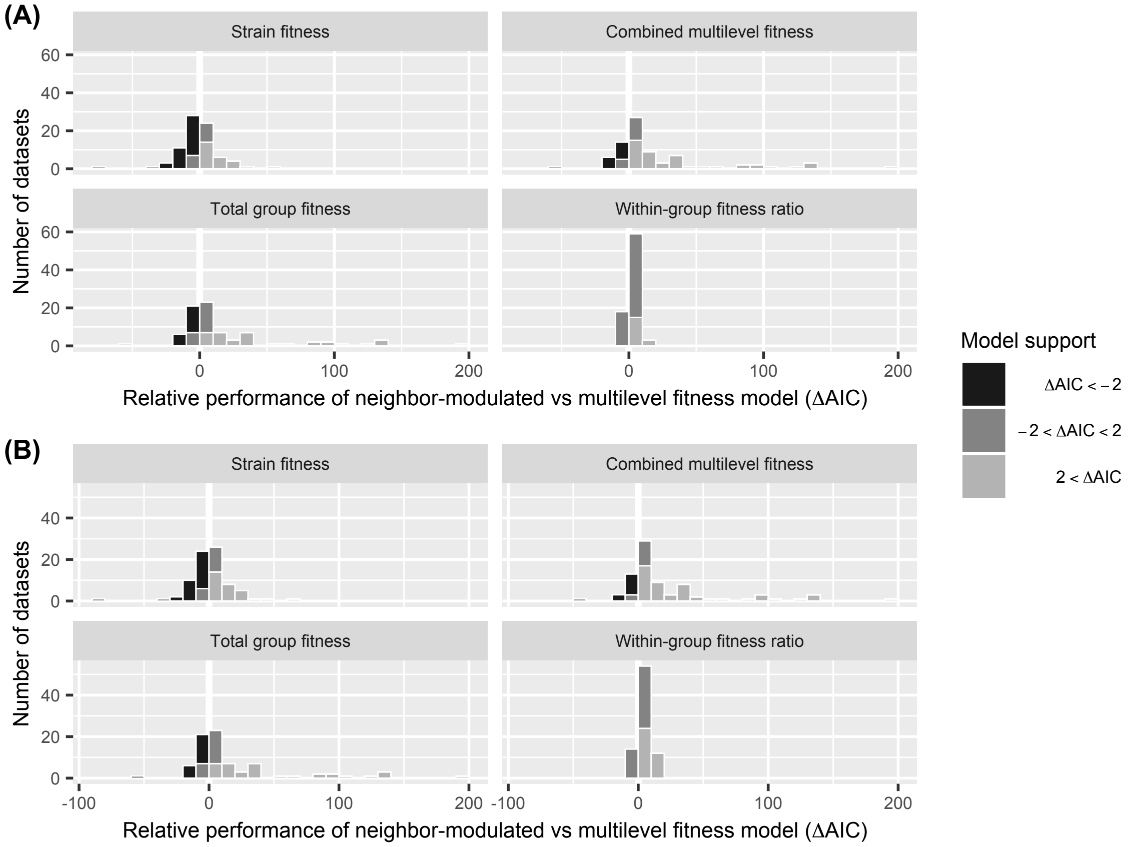

**Figure S4:** Microbial datasets are sometimes more easily analyzed as neighbor-modulated fitness and sometimes as multilevel fitness. Data show relative performance of log-linear models fit to one set of fitness outcomes at predicting the other set. Lower panels show separate components of multilevel model. **(A)** Relative performance when all models are functions of initial mixing proportion. **(B)** Allowing within-group fitness to be a function of log mixing ratio $\ln(q_A/q_B)$ has little overall effect.

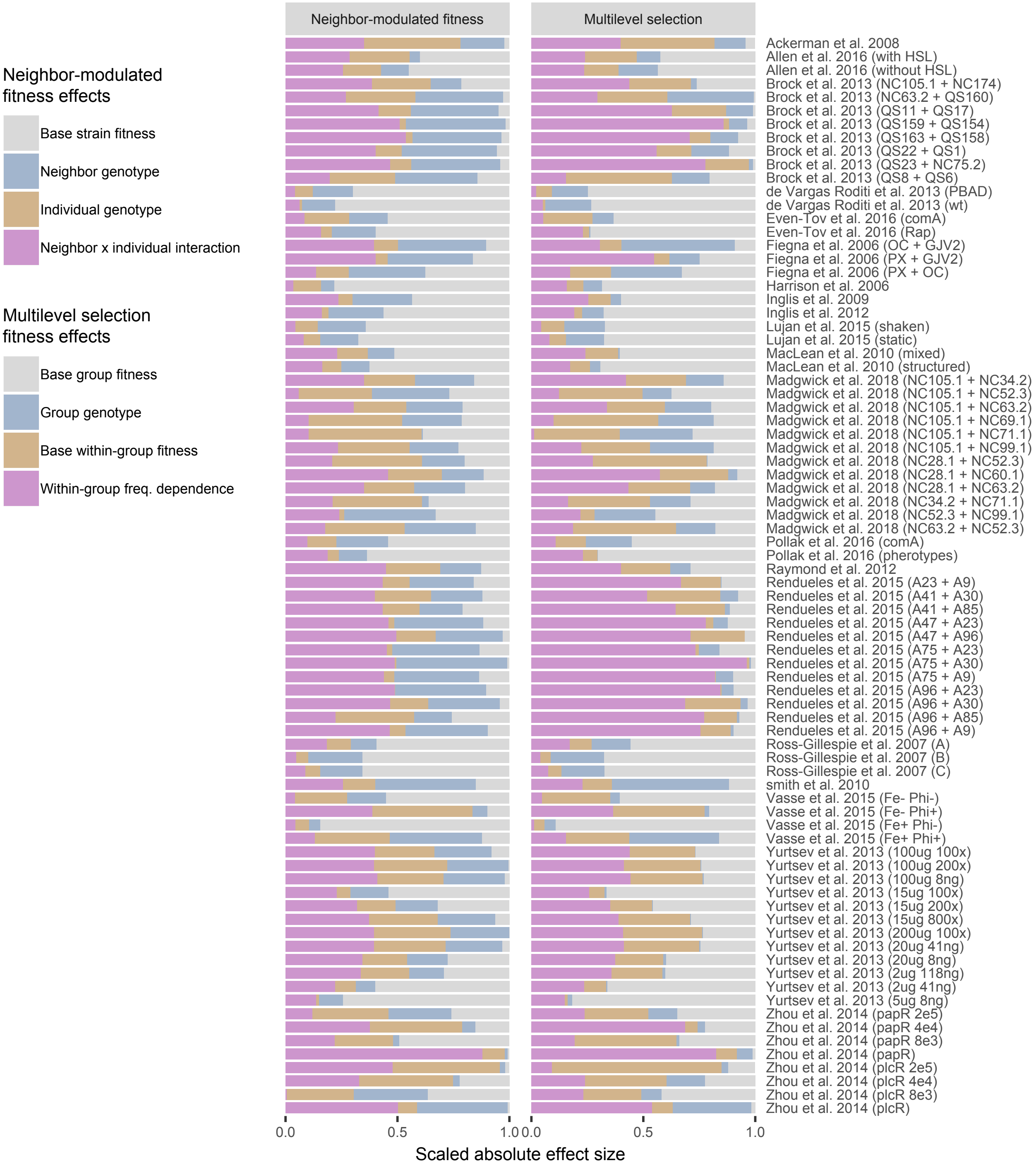

**Figure S5:** Fitness effect sizes in analyzed datasets. Data show absolute Malthusian effect sizes, scaled as a fraction of all observed effects. Colors indicate analogous effects.

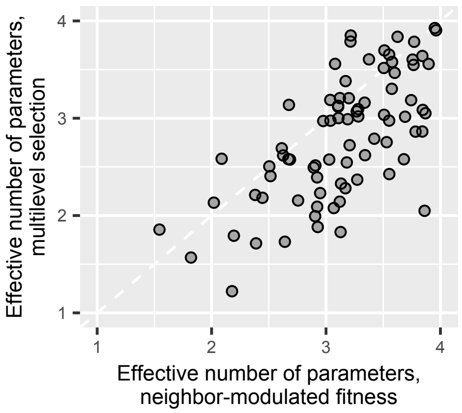

**Figure S6:** Multilevel selection models often describe the results of microbial mix experiments with fewer effective parameters than neighbor-modulated fitness models. Data show effective number of parameters in log-linear models fit to the same dataset.

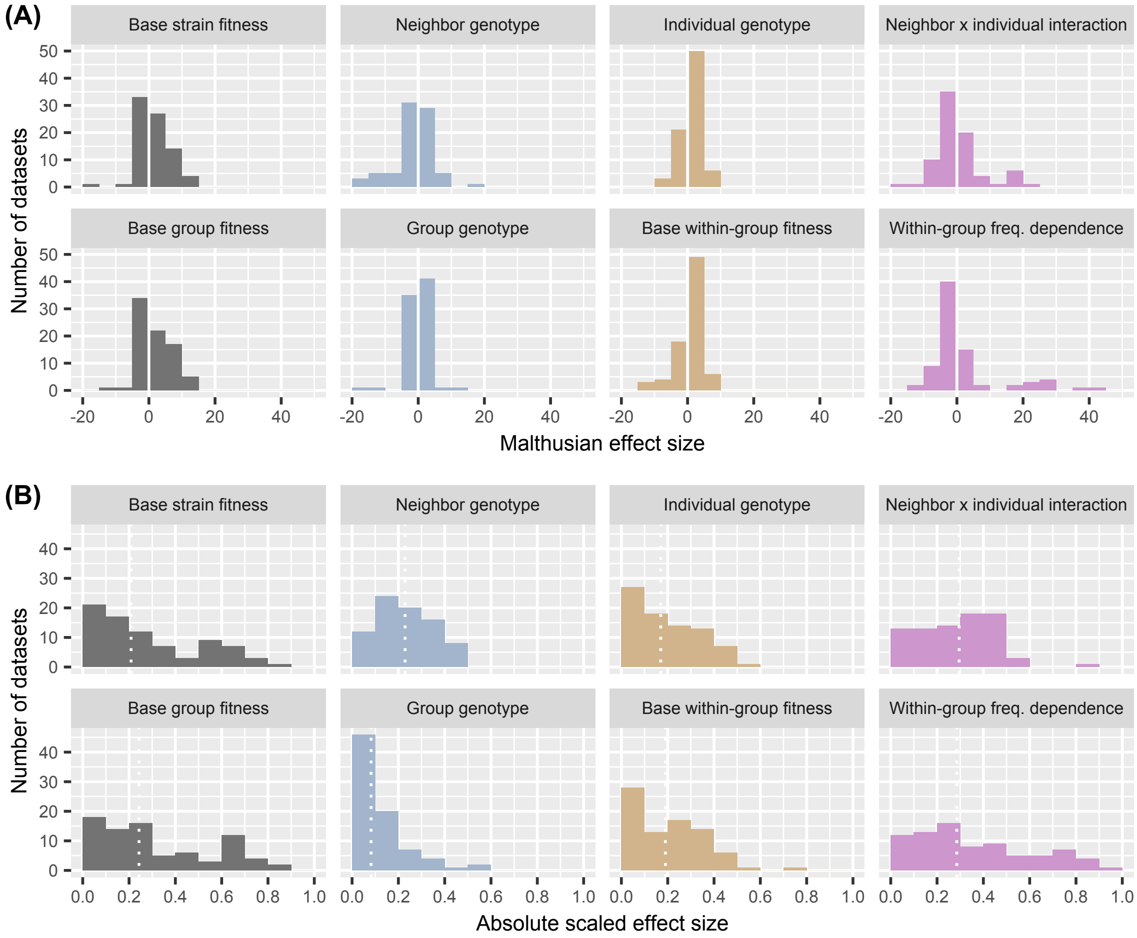

**Figure S7:** **(A)** Distribution of Malthusian effect sizes across datasets for fitted neighbor-modulated and multilevel fitness models. **(B)** Distribution of Malthusian effect sizes scaled as fraction of all effects in fitted models. Within-group frequency dependence and interactions between individual and neighbor genotype are often sizeable. The small effect of group genotype on total group fitness, compared to the effect of neighbor genotype on strain fitness, indicates widespread soft selection. Dashed lines shows medians. Colors indicate analogous effects.

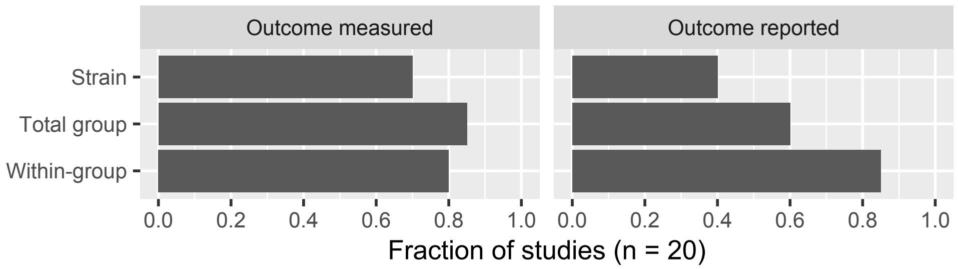

**Figure S8:** Fitness outcomes measured and reported in original studies of analyzed datasets. In principle, studies could measure and analyze all three outcomes. See Table S2 for details of individual papers.

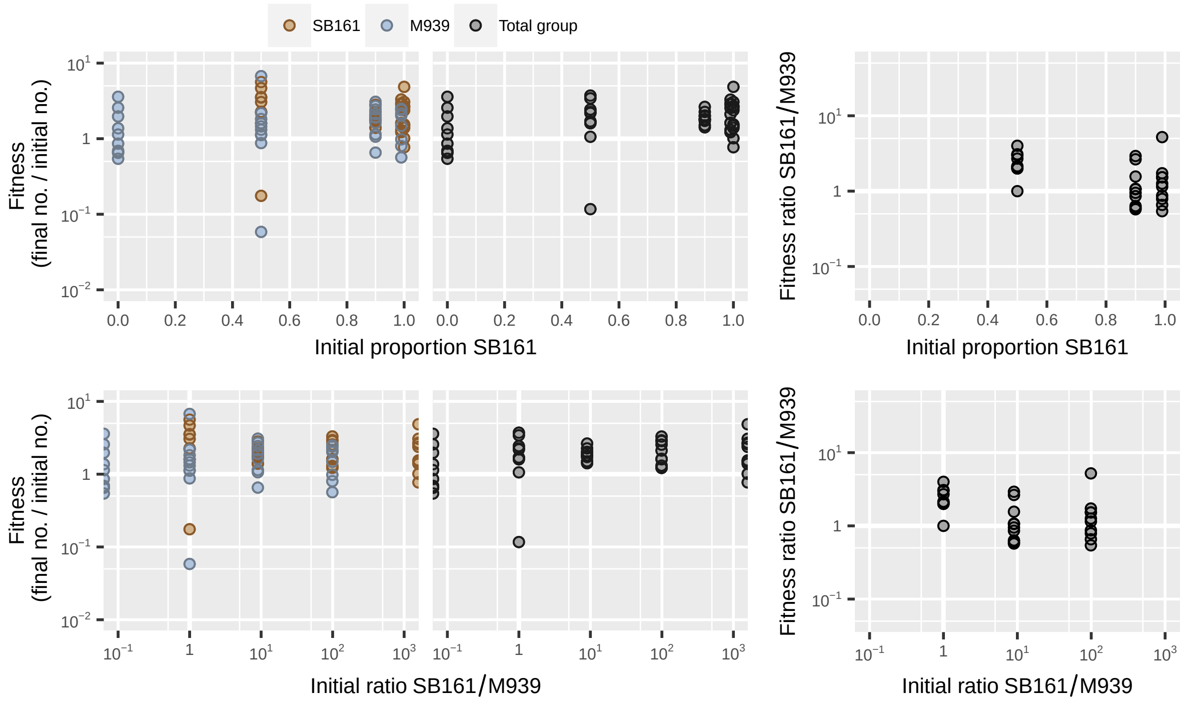

**Figure S9:** Data from Ackermann *et al.* [1] showing inital establishment in mouse ceca of wild-type *Salmonella enterica* serovar Typhimurium strain M939 and ΔSPI-1 virulence factor mutant SB161.

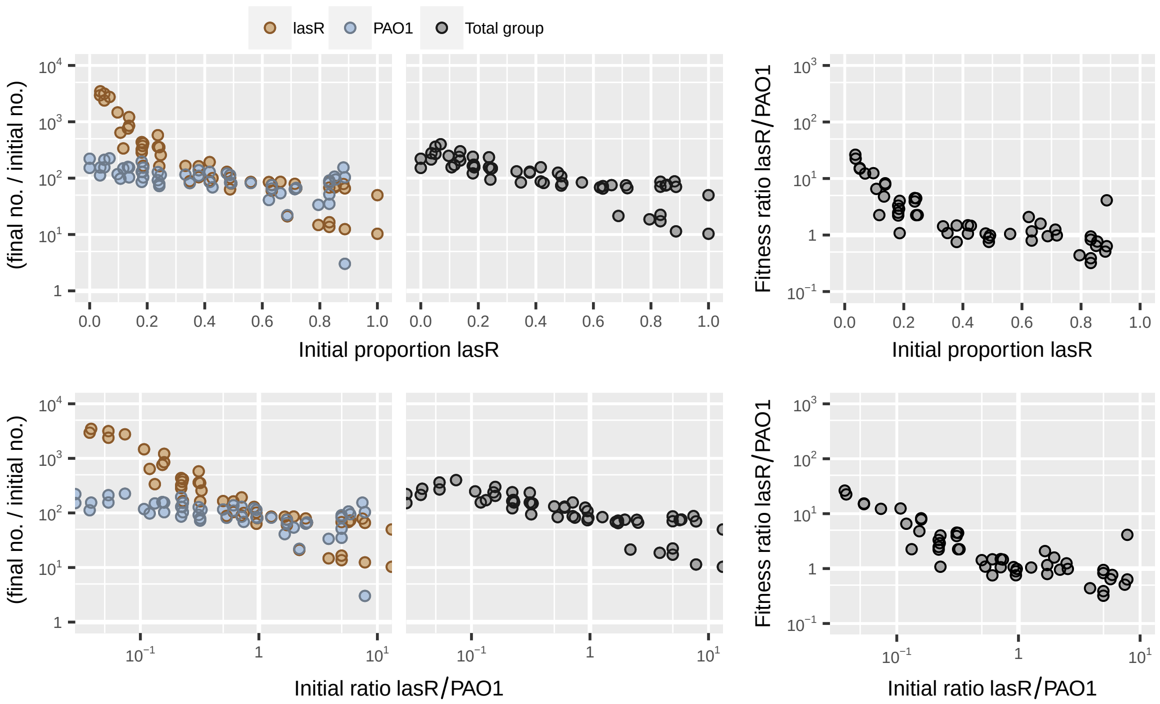

**Figure S10:** Data from Allen *et al.* [2] showing mix experiment with wild-type *Pseudomonas aeruginosa* strain PAO1 and a signal-blind *lasR* mutant.

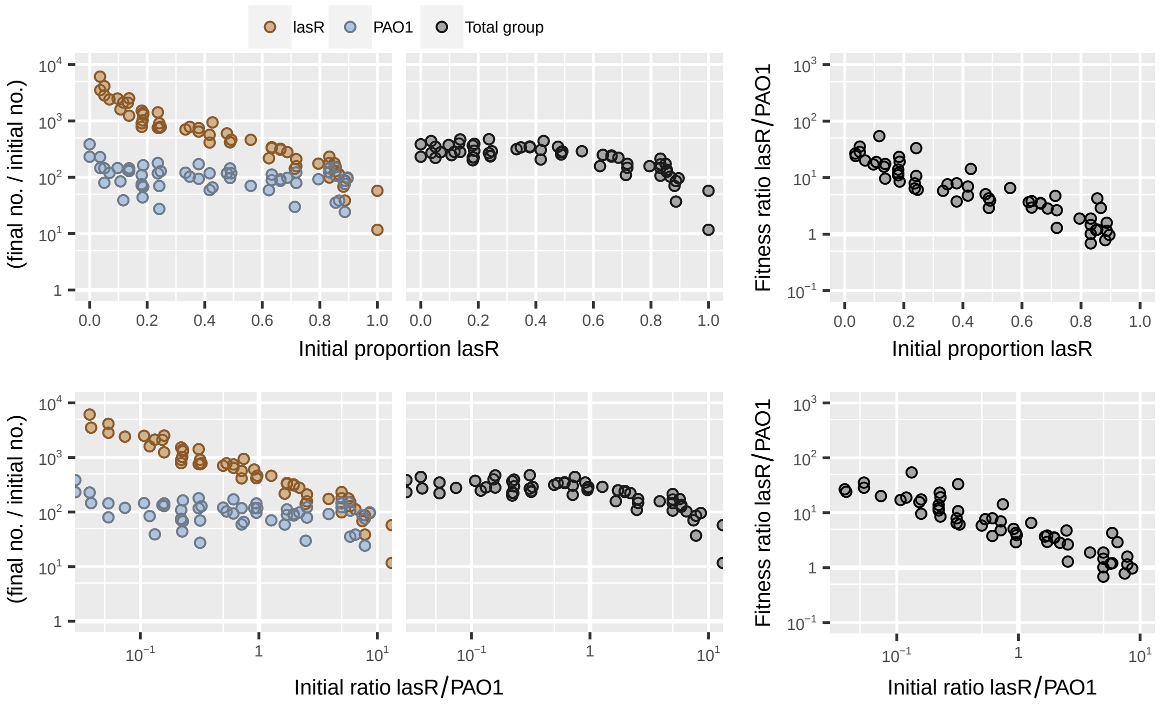

**Figure S11:** Data from Allen *et al.* [2] showing mix experiment with wild-type *Pseudomonas aeruginosa* strain PAO1 and a signal-blind *lasR* mutant, with signalling molecules added to culture media.

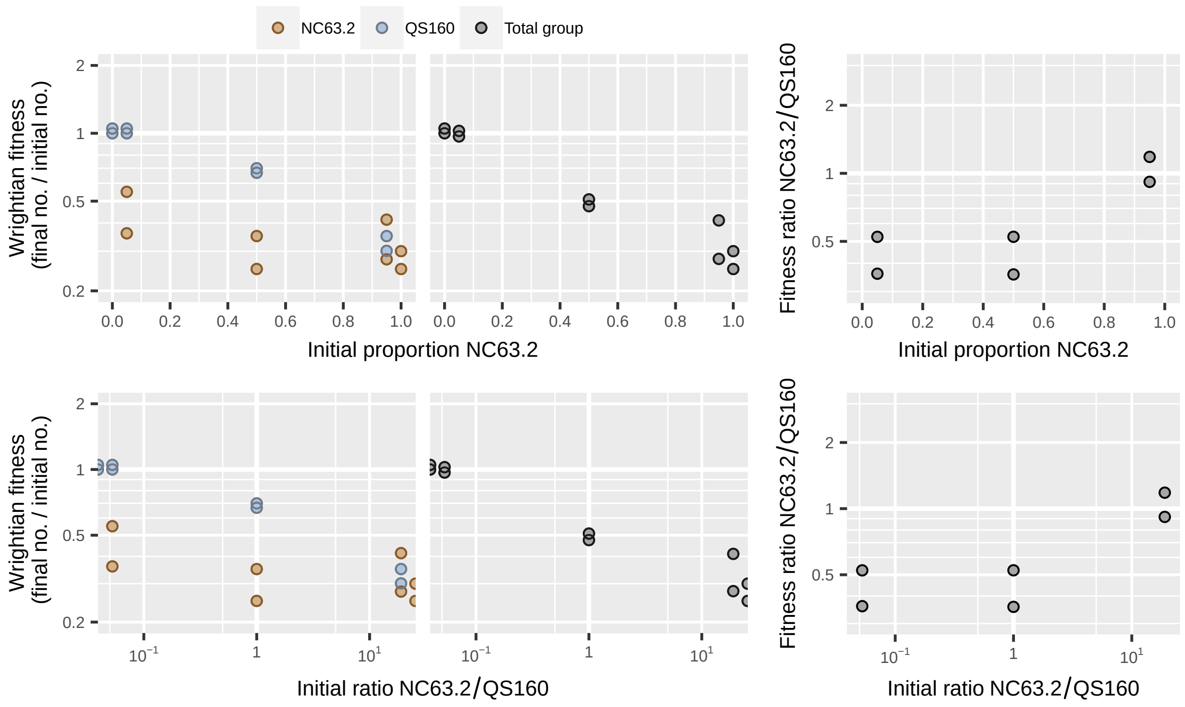

**Figure S12:** Dataset from Brock *et al.* [3] showing codevelopment of *Dictyostelium discoideum* farmer strain NC63.2 and nonfarmer strain QS160.

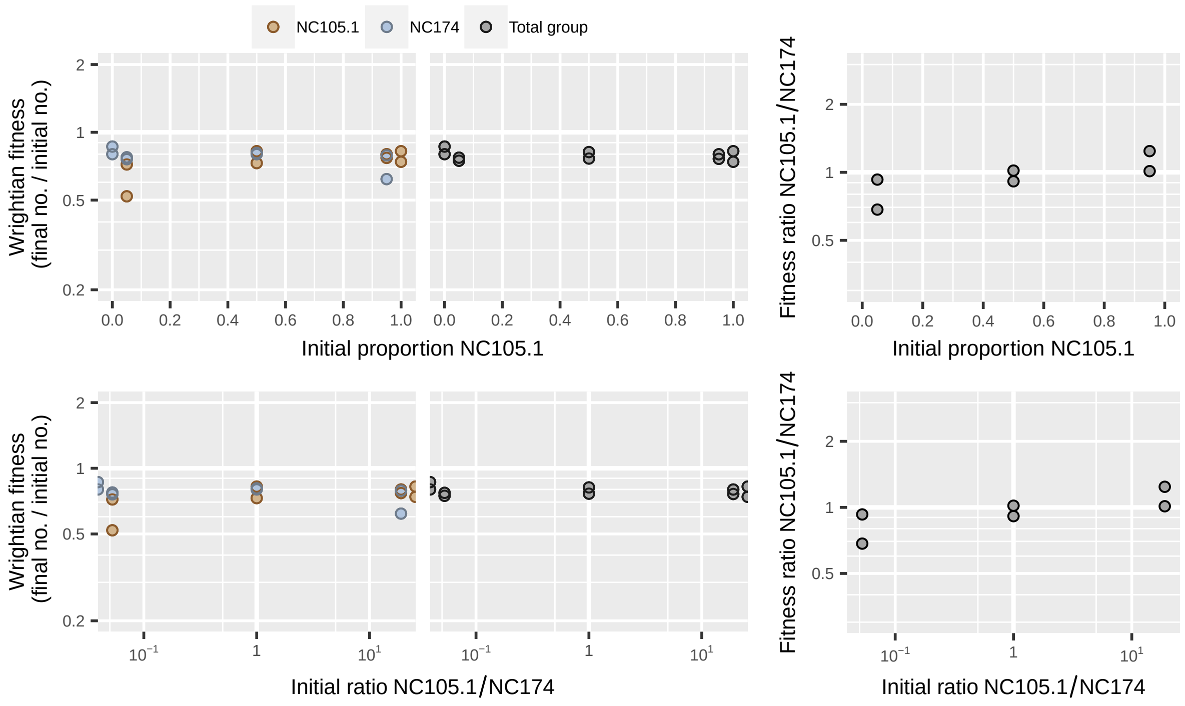

**Figure S13:** Dataset from Brock *et al.* [3] showing codevelopment of *Dictyostelium discoideum* farmer strain NC105.1 and nonfarmer strain NC174.

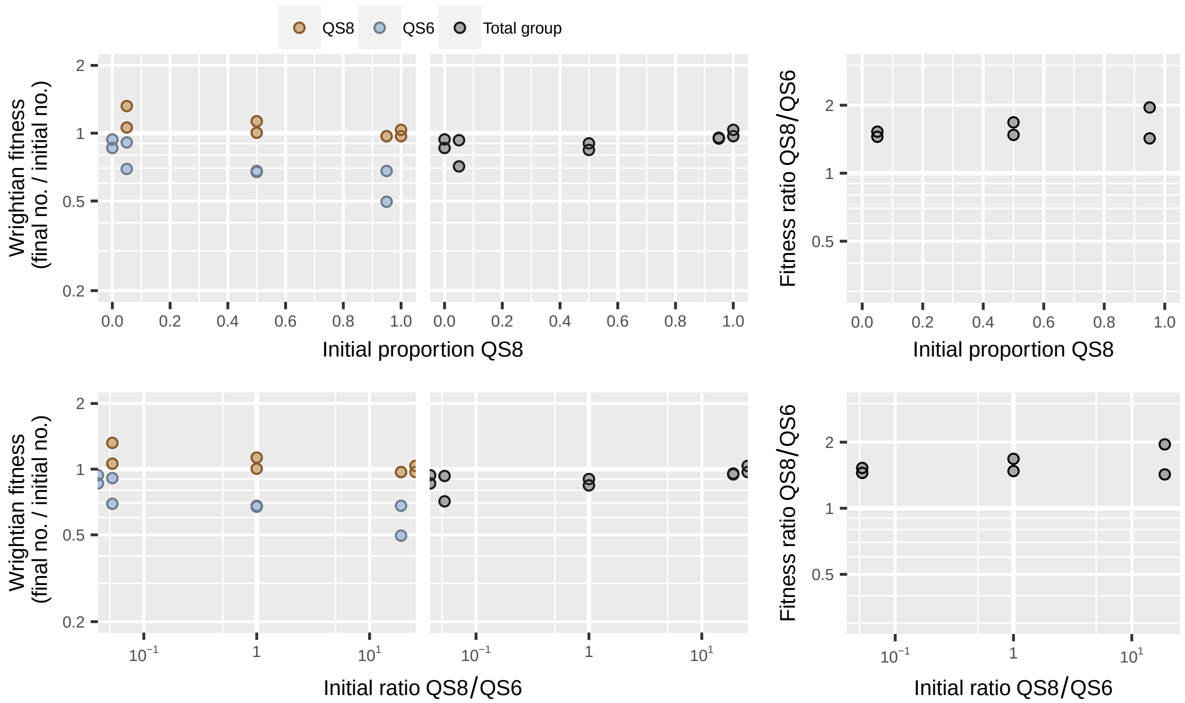

**Figure S14:** Dataset from Brock *et al.* [3] showing codevelopment of *Dictyostelium discoideum* farmer strain QS8 and nonfarmer strain QS6.

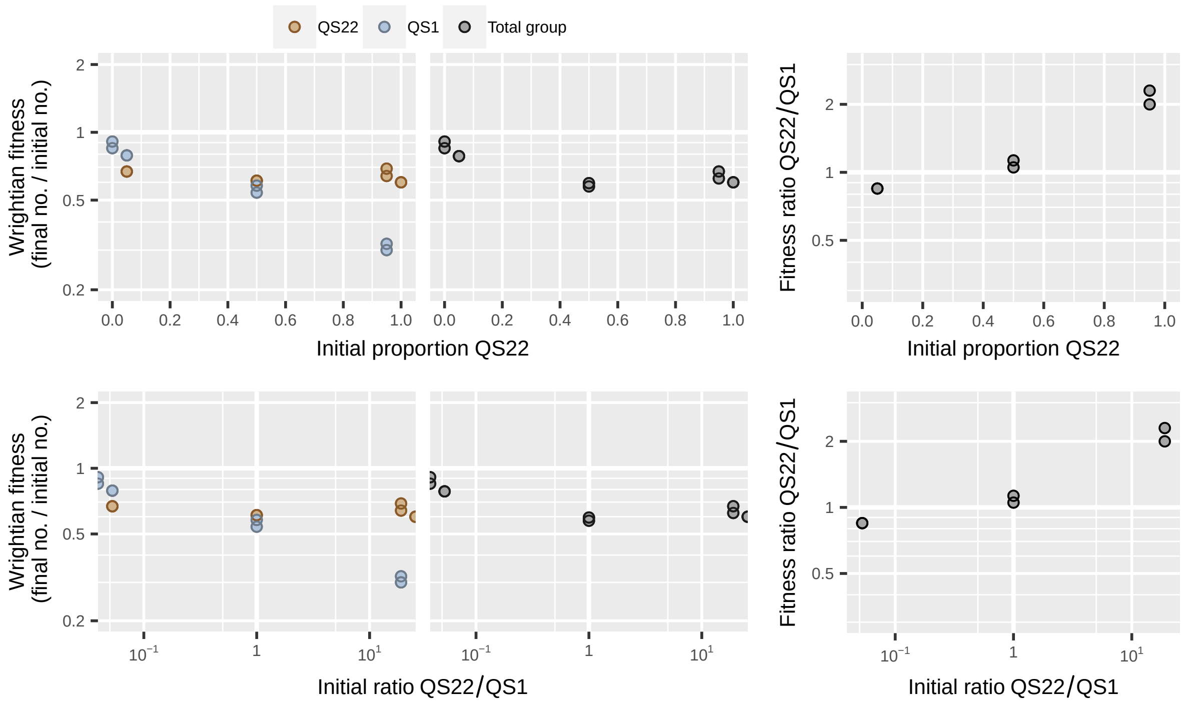

**Figure S15:** Dataset from Brock *et al.* [3] showing codevelopment of *Dictyostelium discoideum* farmer strain QS22 and nonfarmer strain QS1.

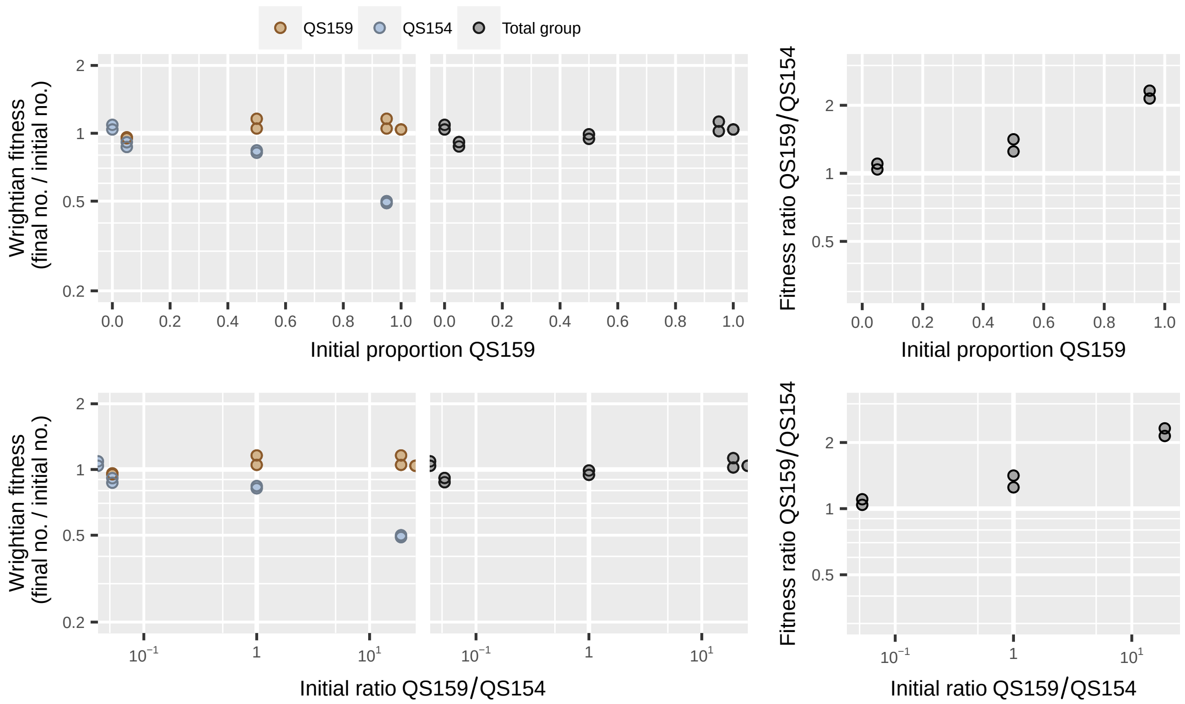

**Figure S16:** Dataset from Brock *et al.* [3] showing codevelopment of *Dictyostelium discoideum* farmer strain QS159 and nonfarmer strain QS154.

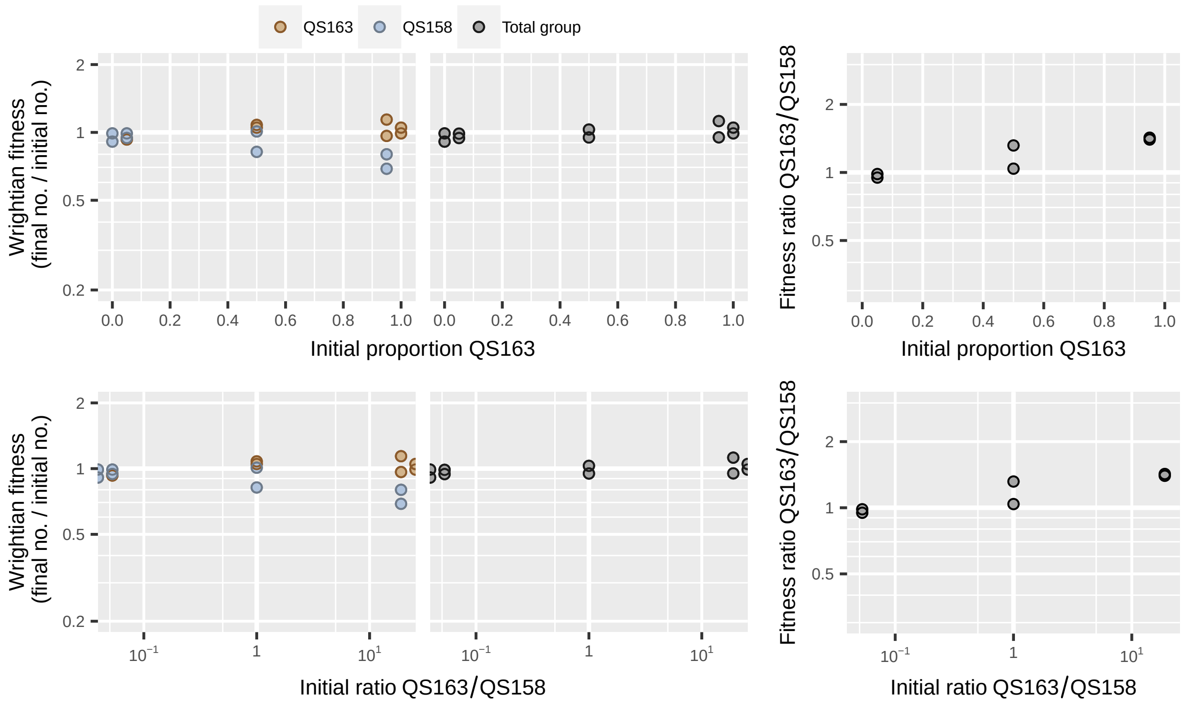

**Figure S17:** Dataset from Brock *et al.* [3] showing codevelopment of *Dictyostelium discoideum* farmer strain QS163 and nonfarmer strain QS158.

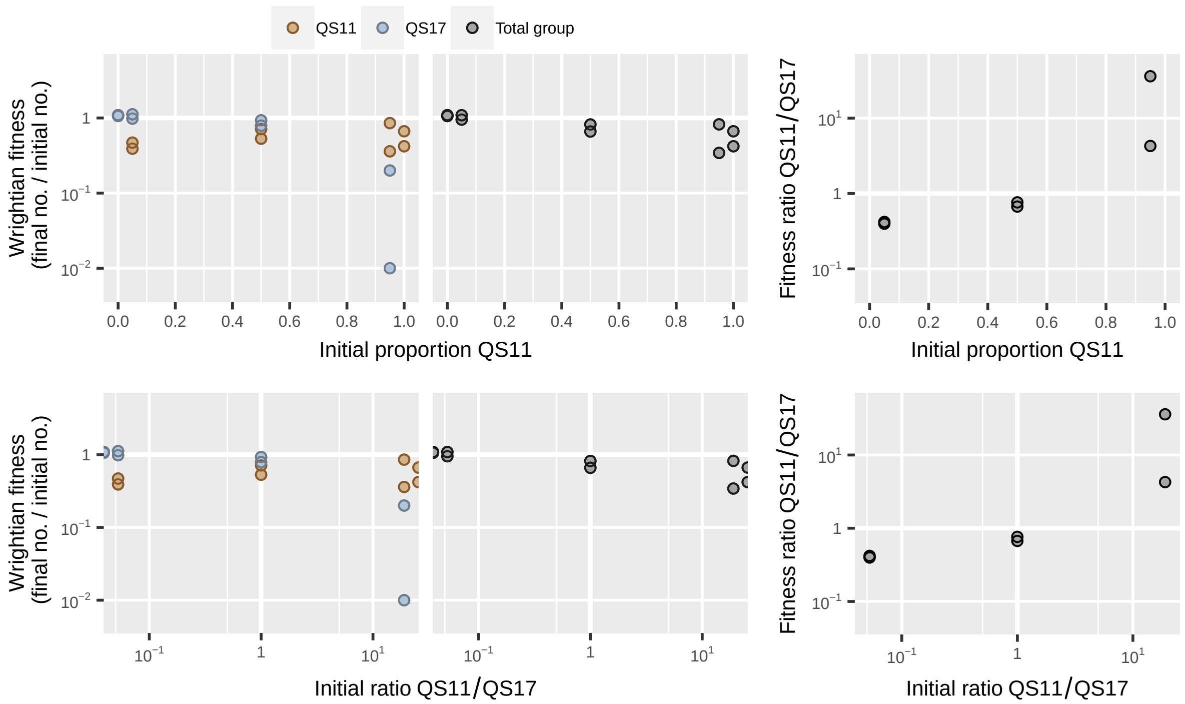

**Figure S18:** Dataset from Brock *et al.* [3] showing codevelopment of *Dictyostelium discoideum* farmer strain QS11 and nonfarmer strain QS17.

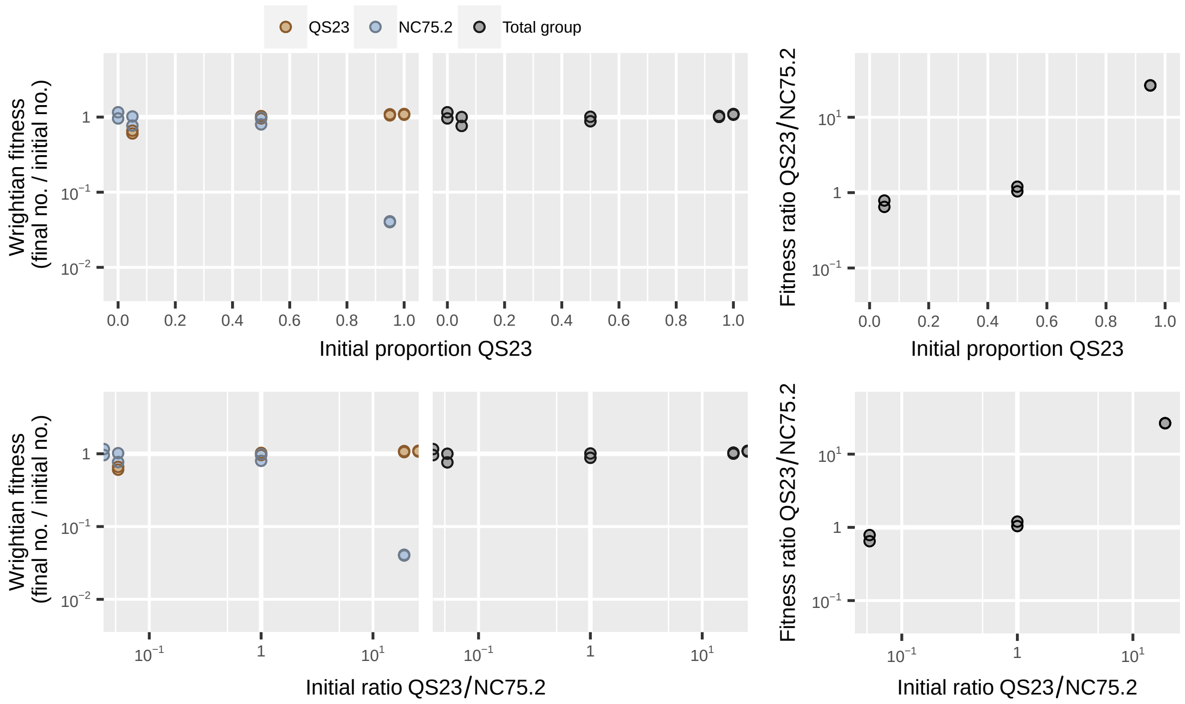

**Figure S19:** Dataset from Brock *et al.* [3] showing codevelopment of *Dictyostelium discoideum* farmer strain QS23 and nonfarmer strain NC75.2.

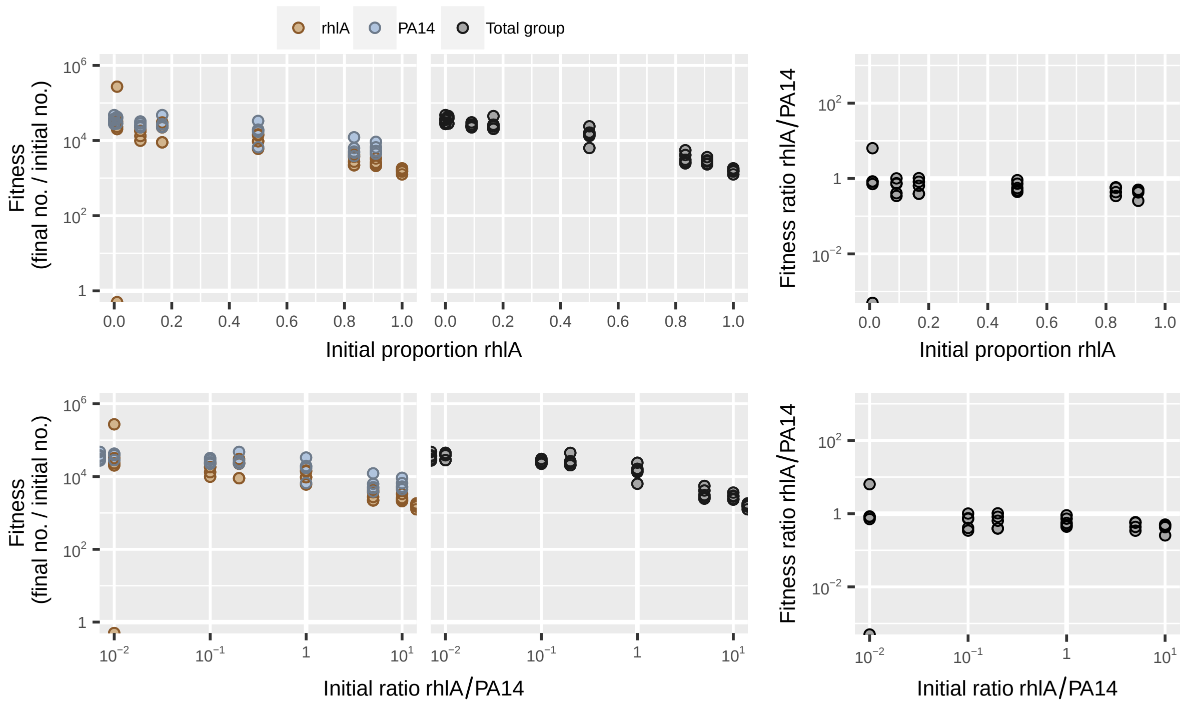

**Figure S20:** Dataset from de Vargas Roditi *et al.* [4] showing mix experiment on swarming media with wild-type *Pseudomonas aeruginosa* strain PA14 and a ΔrhlA biosurfactant nonproducer.

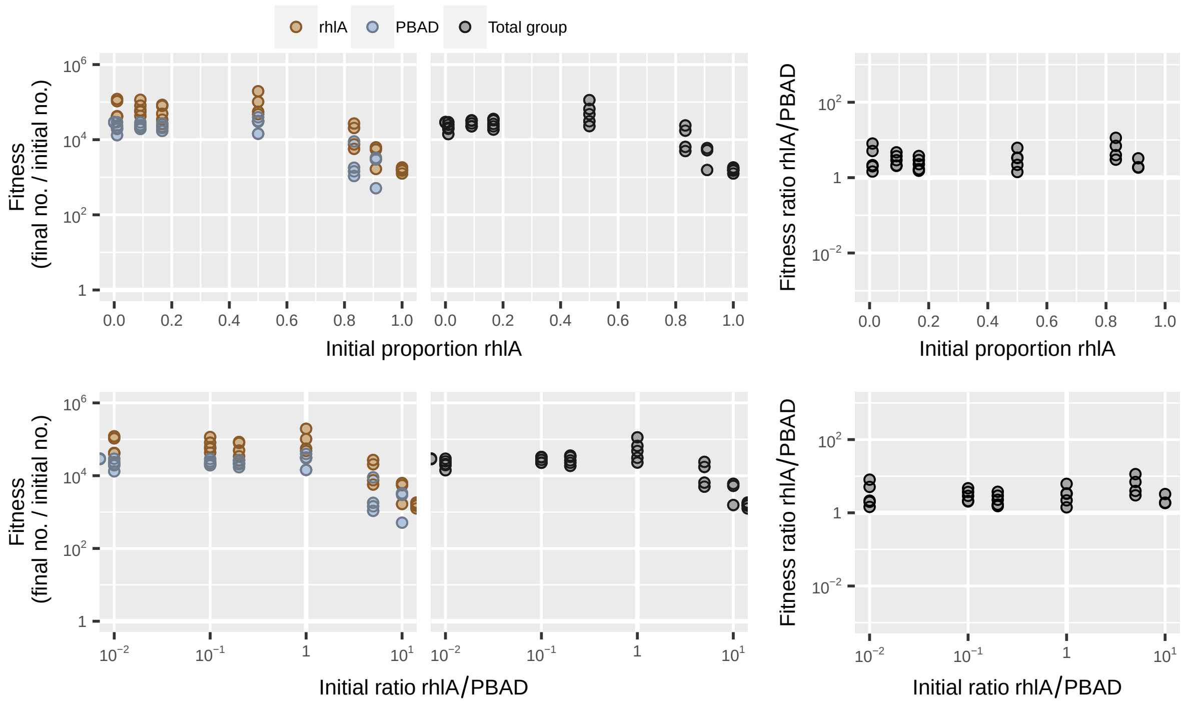

**Figure S21:** Dataset from de Vargas Roditi *et al.* [4] showing mix experiment on swarming media with engineered *Pseudomonas aeruginosa* strain PBAD and a ΔrhlA biosurfactant nonproducer.

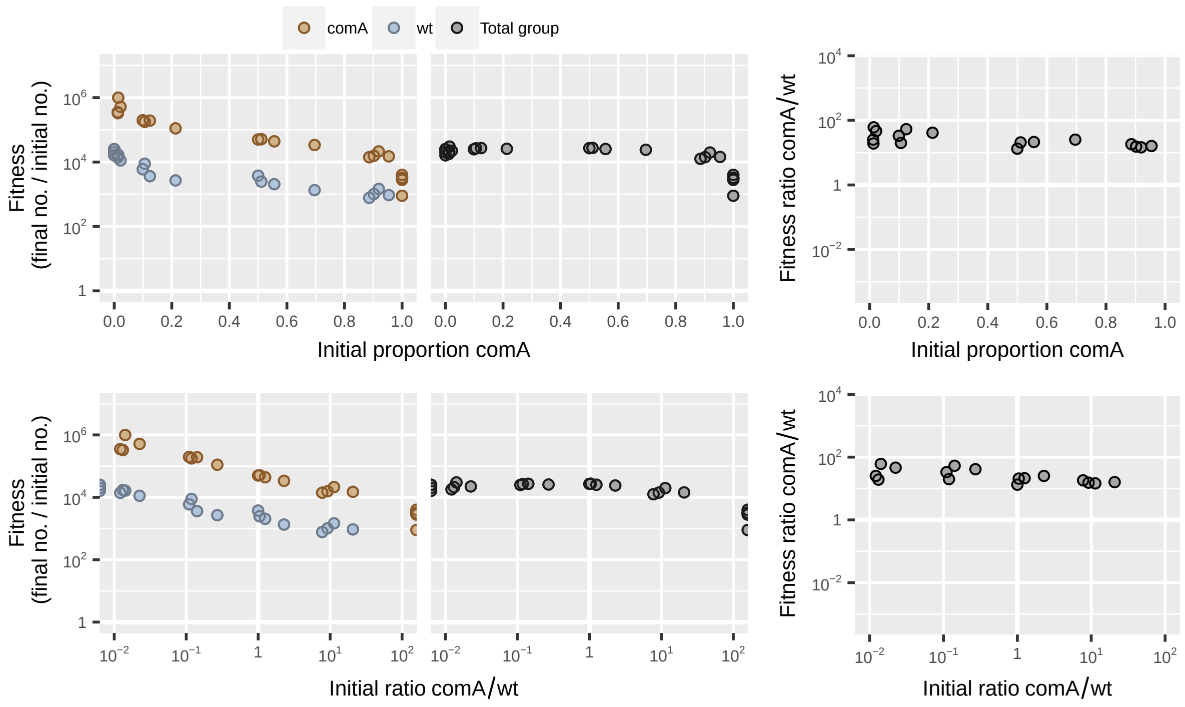

**Figure S22:** Dataset from Even-Tov *et al.* [5] showing mix experiment with wild type *Bacillus subtilis* and a $\Delta$*comA* mutant on swarming media.

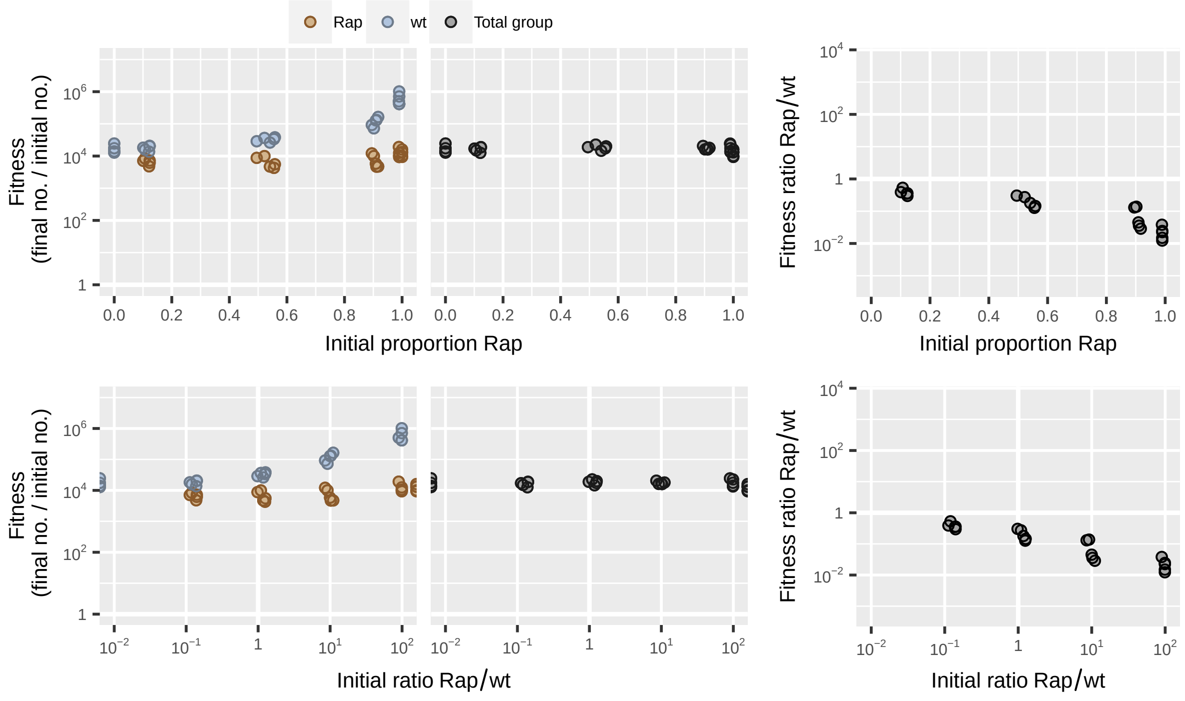

**Figure S23:** Dataset from Even-Tov *et al.* [5] showing mix experiment with wild-type *Bacillus subtilis* and a $\Delta$rapFphrF mutant on swarming media.

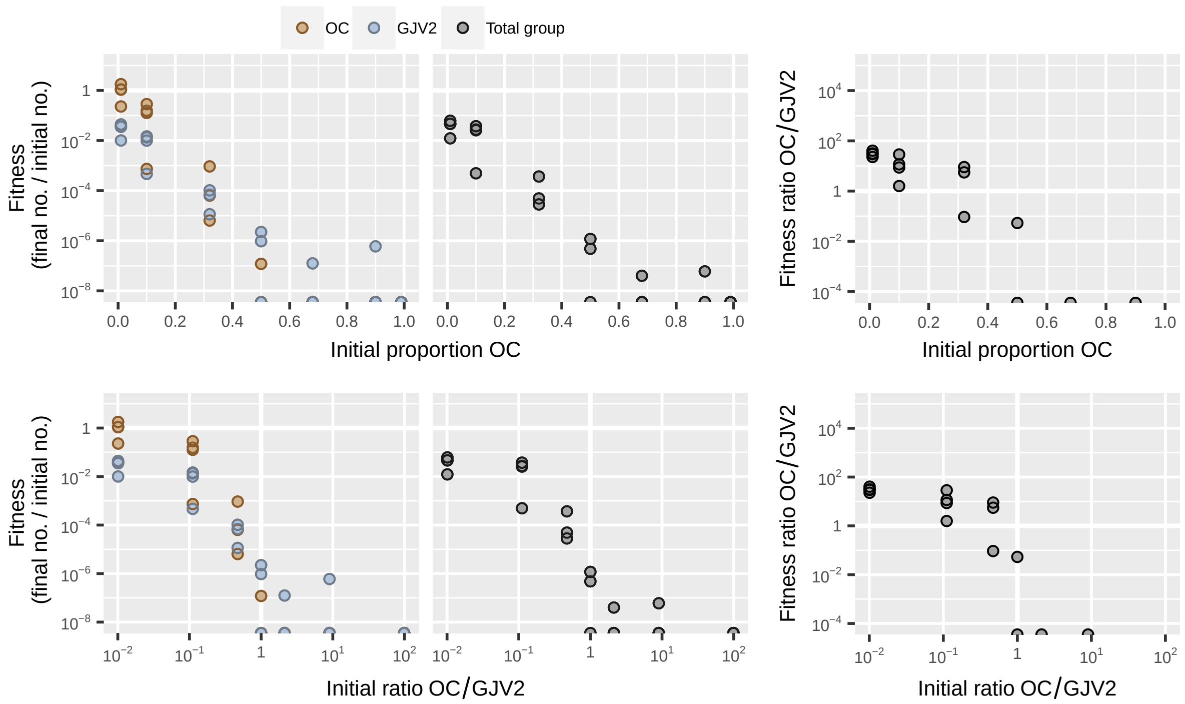

**Figure S24:** Dataset from Fiegna *et al.* [6] showing codevelopment of evolved *Myxococcus xanthus* strain OC and ancestral wild-type strain GJV2.

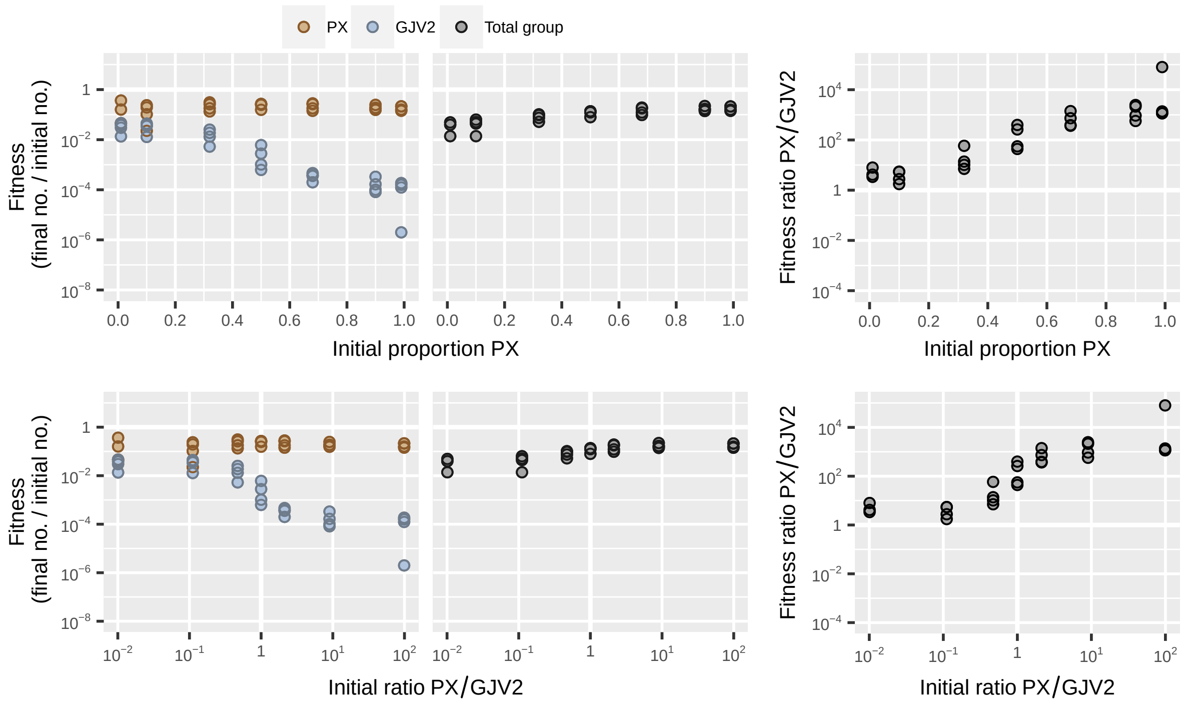

**Figure S25:** Dataset from Fiegna *et al.* [6] showing codevelopment of evolved *Myxococcus xanthus* strain PX and ancestral wild-type strain GJV2.

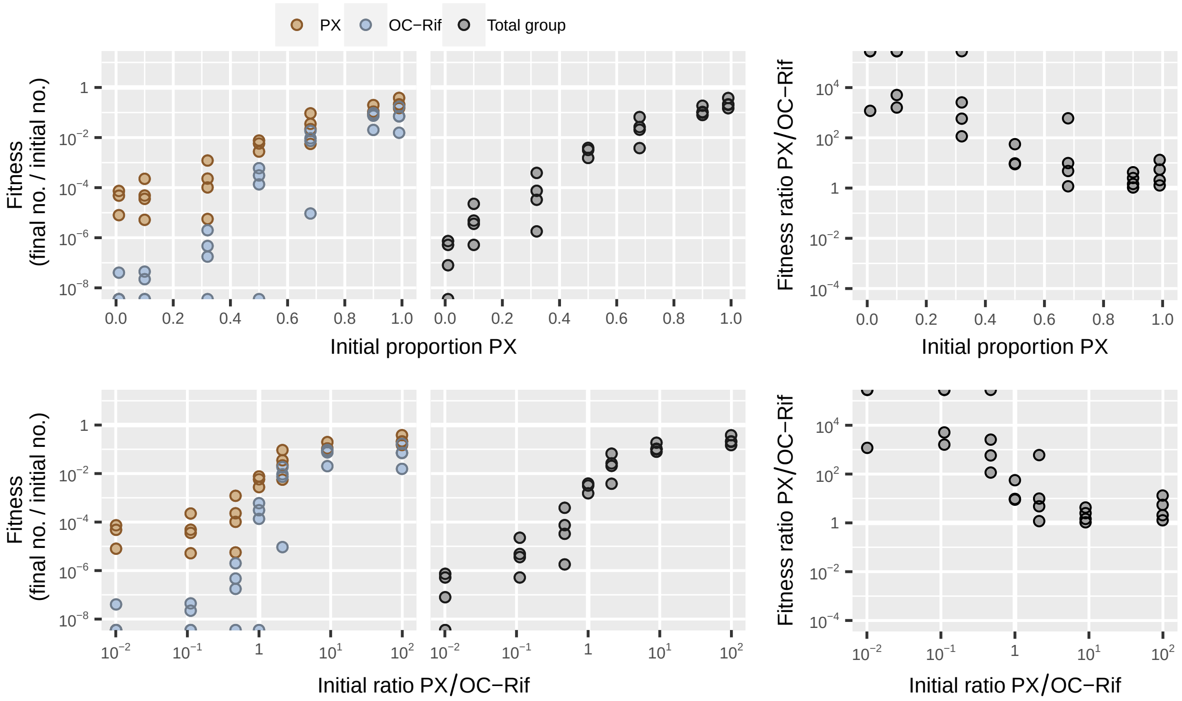

**Figure S26:** Dataset from Fiegna *et al.* [6] showing codevelopment of evolved *Myxococcus xanthus* strain PX and evolved strain OC.

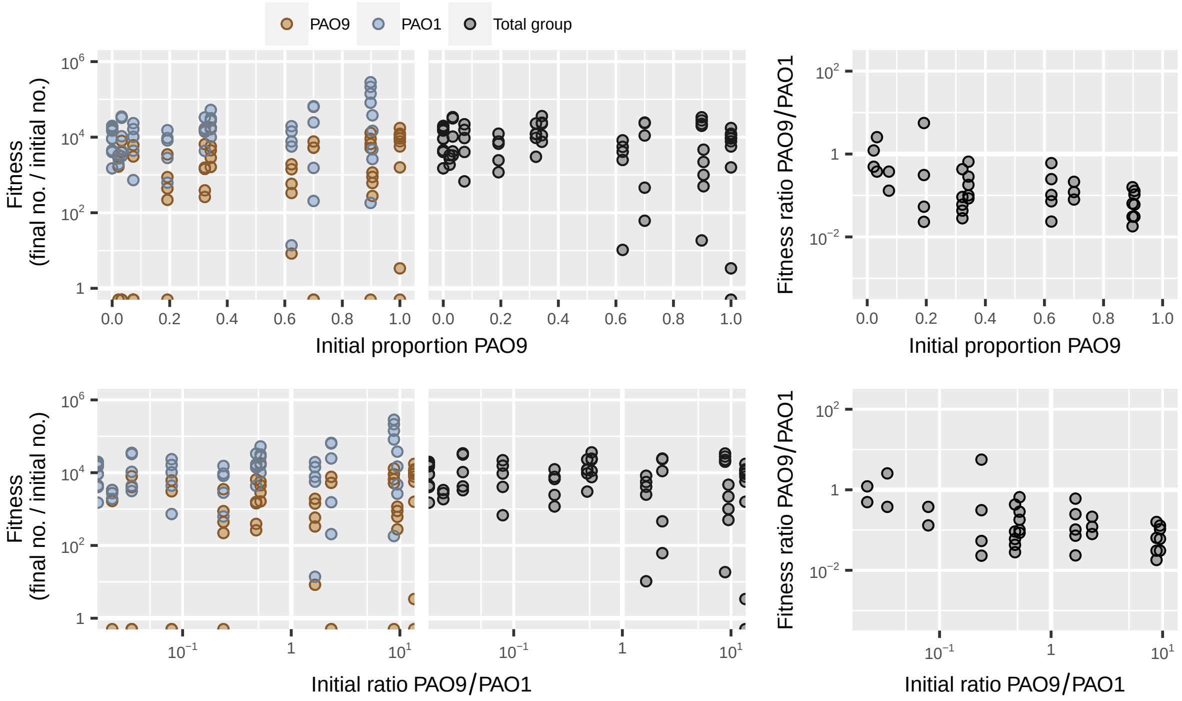

**Figure S27:** Dataset from Harrison *et al.* [7] showing coinfection of waxmoth larvae with wild-type *Pseudomonas aeruginosa* strain PAO1 and siderophore nonproducer PAO9.

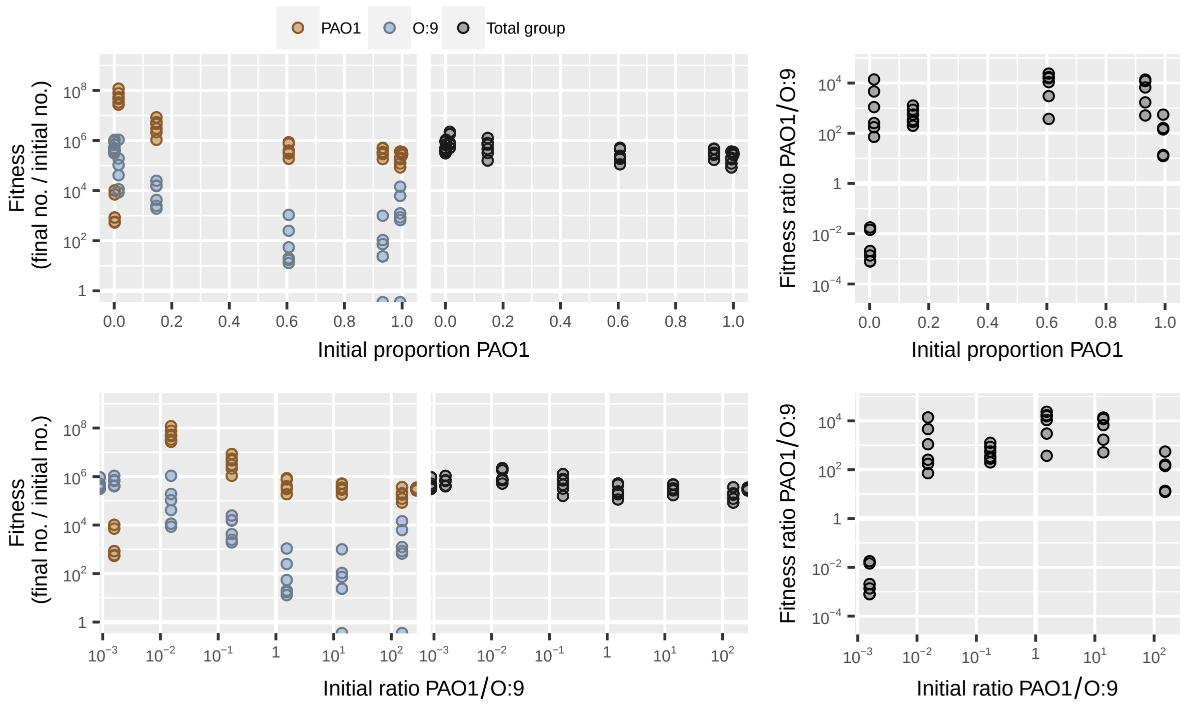

**Figure S28:** Dataset from Inglis *et al.* [8] showing mix experiment with wild-type *Pseudomonas aeruginosa* strain PAO1 and bacteriocin producer O:9.

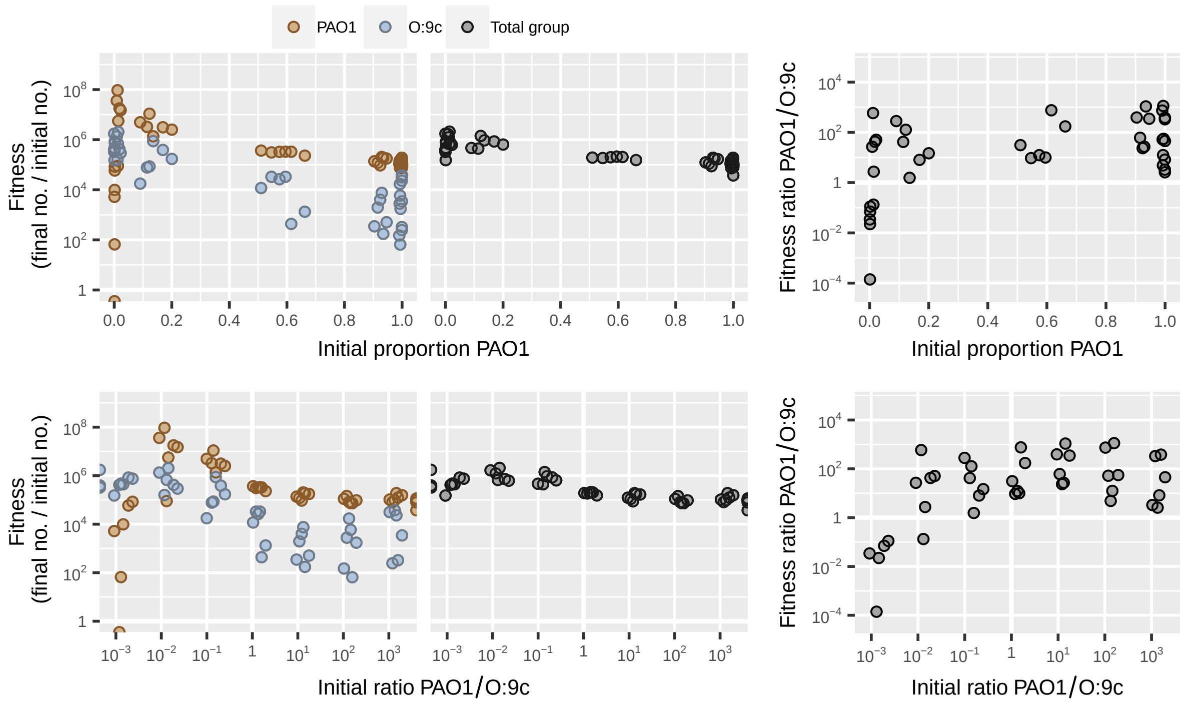

**Figure S29:** Dataset from Inglis *et al.* [9] showing mix experiment with wild-type *Pseudomonas aeruginosa* strain PAO1 and bacteriocin producer O:9c.

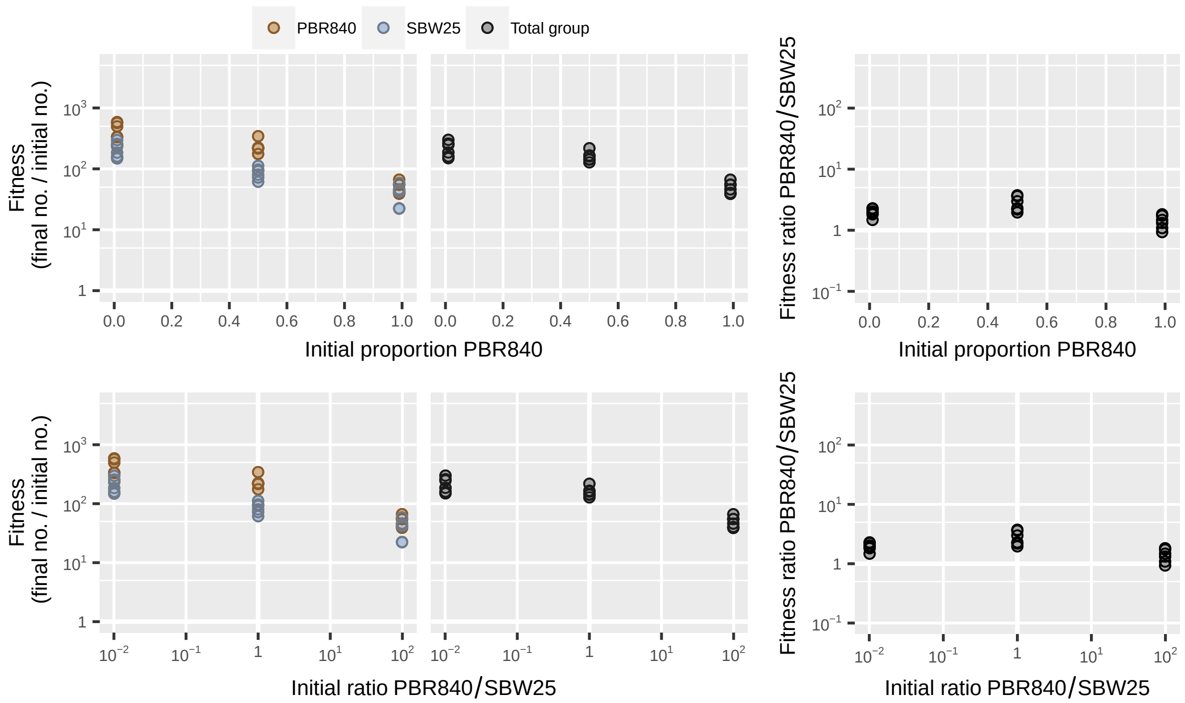

**Figure S30:** Dataset from Luján *et al.* [10] showing mix experiment with *Pseudomonas aeruginosa* strain SBW25 and siderophore nonproducer PBR840 in mixed soil microcosms.

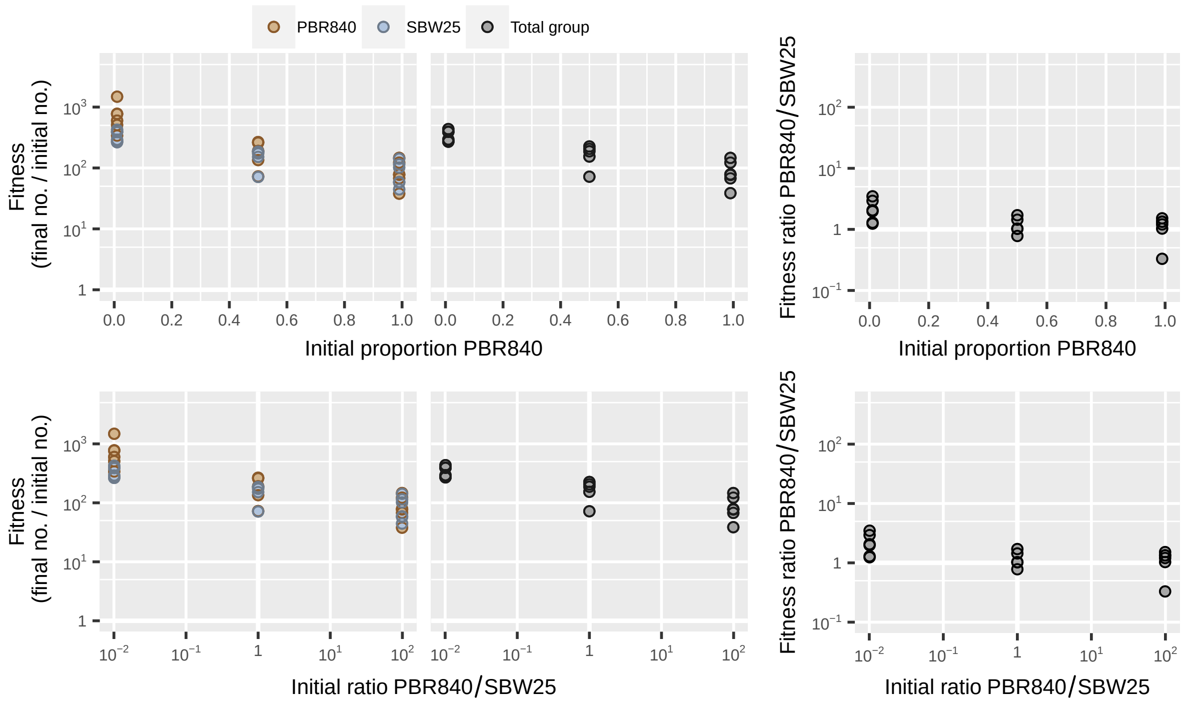

**Figure S31:** Dataset from Luján *et al.* [10] showing mix experiment with *Pseudomonas aeruginosa* strain SBW25 and siderophore nonproducer PBR840 in static soil microcosms.

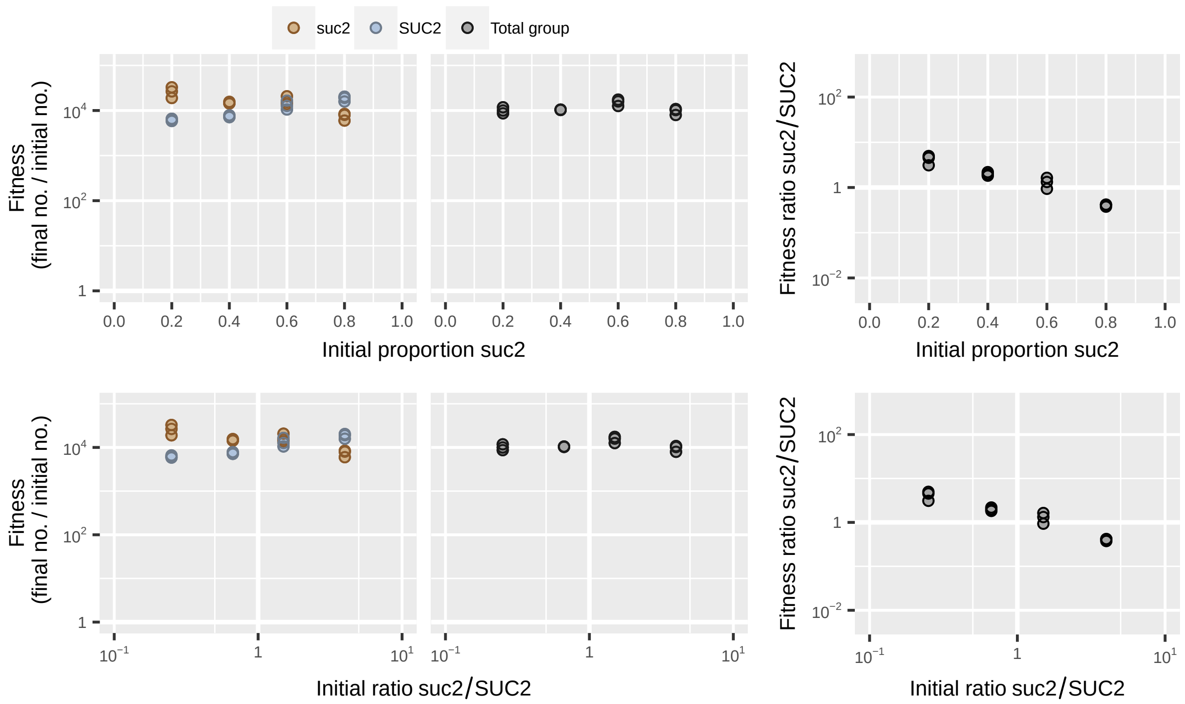

**Figure S32:** Dataset from MacLean *et al.* [11] showing mix experiment with *Sacharomyces cerevisea* strains suc2 and SUC2 without population structure.

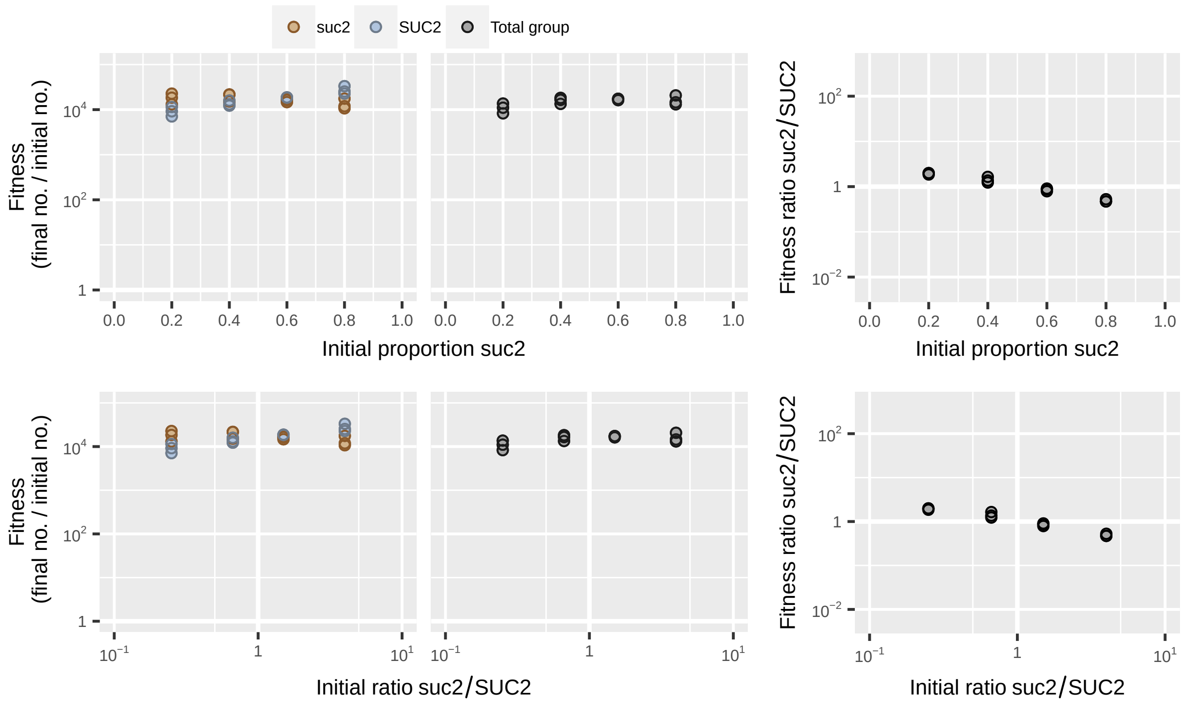

**Figure S33:** Dataset from MacLean *et al.* [11] showing mix experiment with *Sacharomyces cerevisea* strains suc2 and SUC2 with population structure.

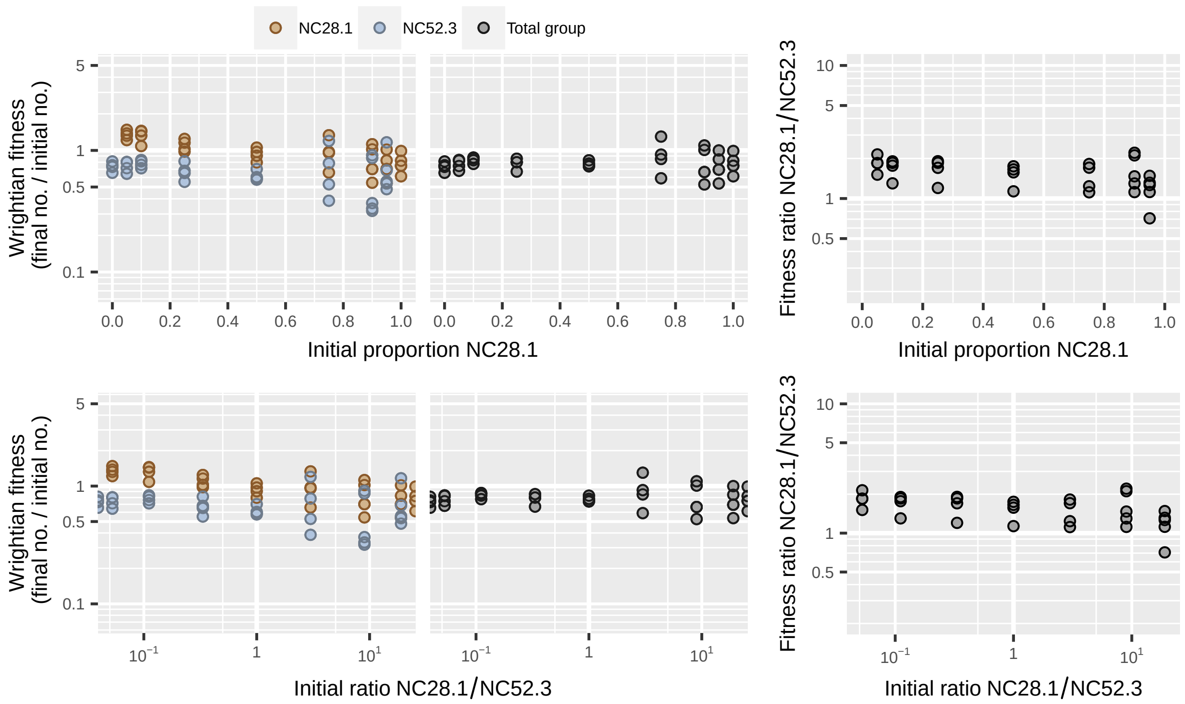

**Figure S34:** Dataset from Madgwick *et al.* [12] showing codevelopment of *Dictyostelium discoideum* strains NC28.1 and NC52.3.

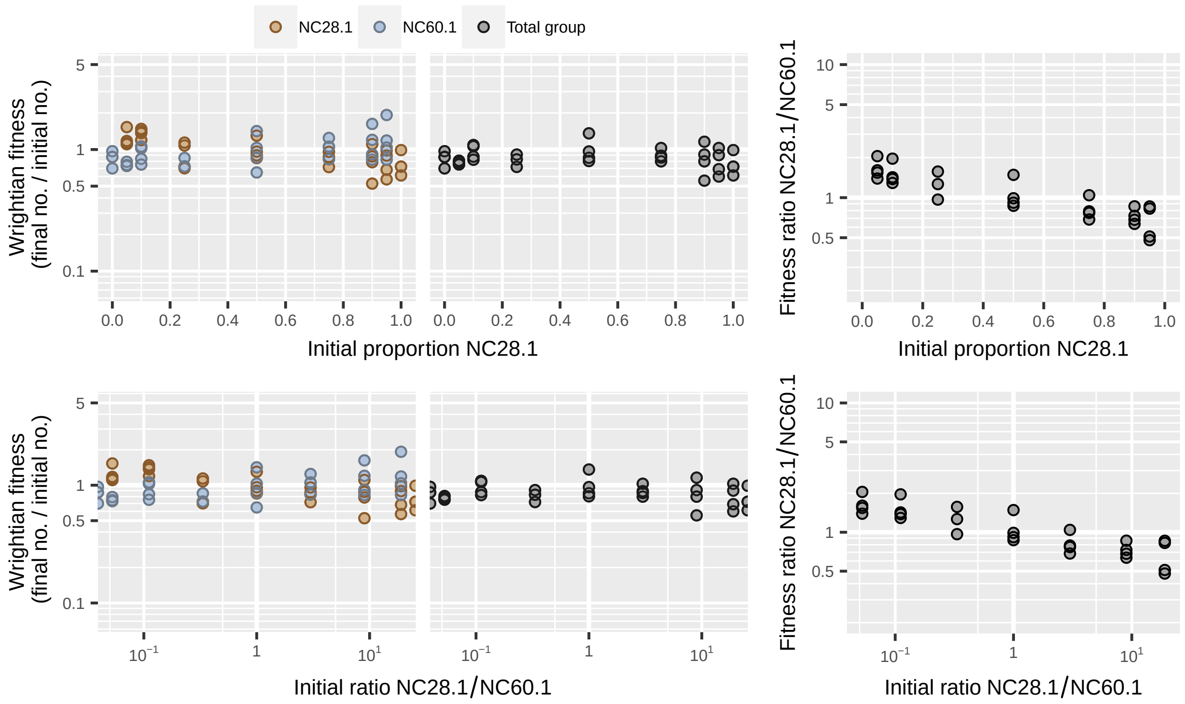

**Figure S35:** Dataset from Madgwick *et al.* [12] showing codevelopment of *Dictyostelium discoideum* strains NC28.1 and NC60.1.

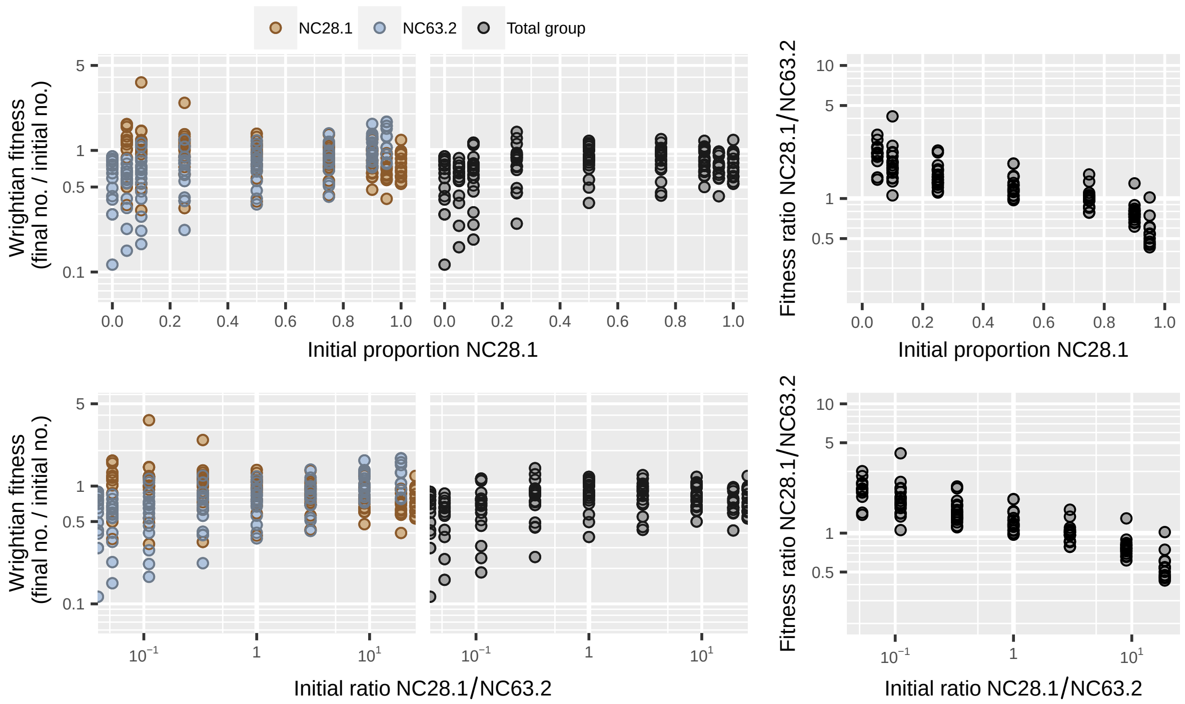

**Figure S36:** Dataset from Madgwick *et al.* [12] showing codevelopment of *Dictyostelium discoideum* strains NC28.1 and NC63.2.

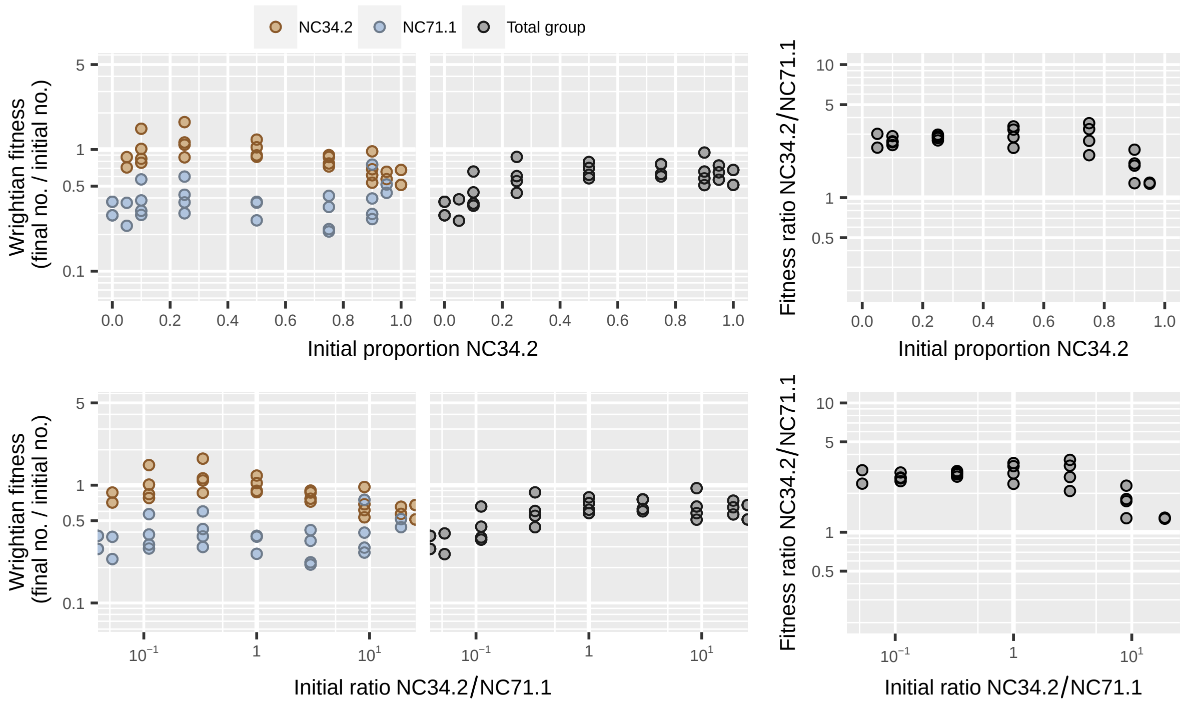

**Figure S37:** Dataset from Madgwick *et al.* [12] showing codevelopment of *Dictyostelium discoideum* strains NC34.2 and NC71.1.

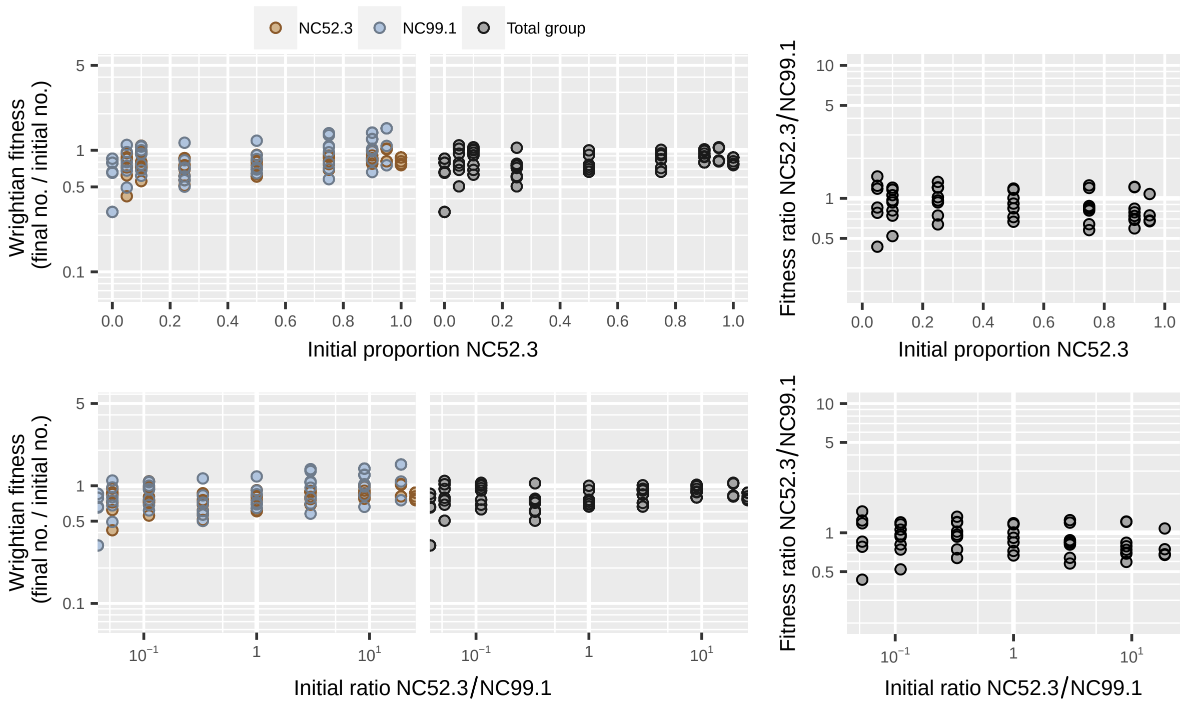

**Figure S38:** Dataset from Madgwick *et al.* [12] showing codevelopment of *Dictyostelium discoideum* strains NC52.3 and NC99.1.

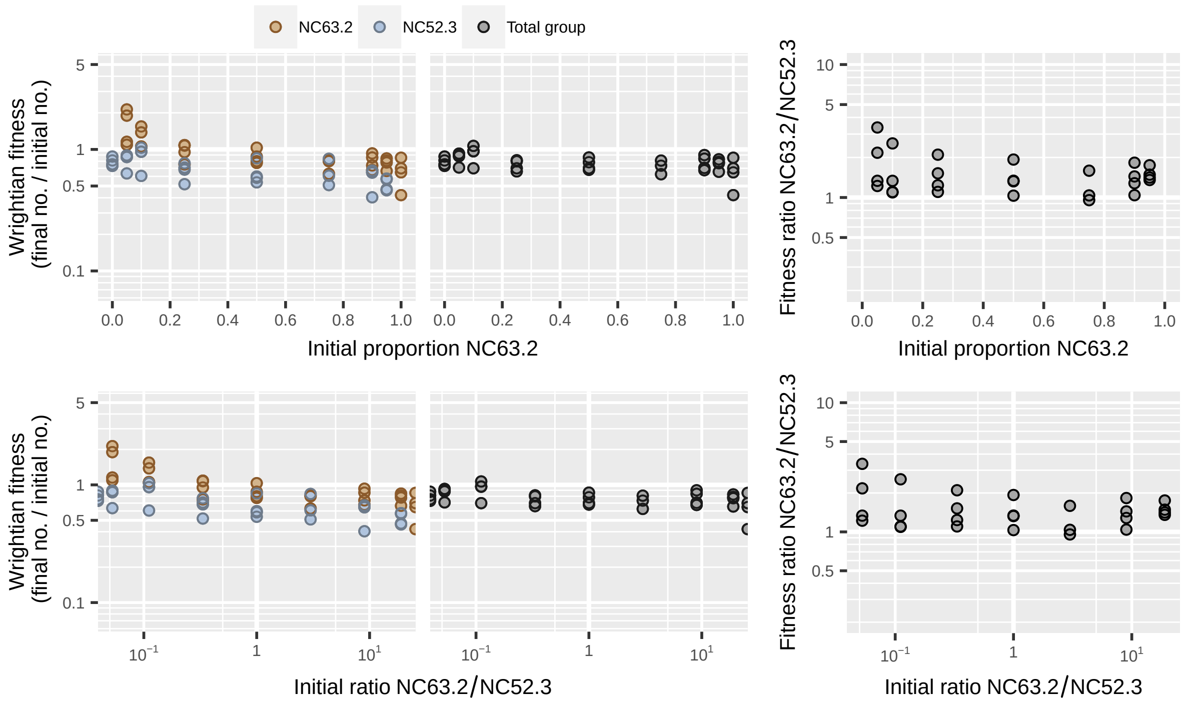

**Figure S39:** Dataset from Madgwick *et al.* [12] showing codevelopment of *Dictyostelium discoideum* strains NC63.2 and NC52.3.

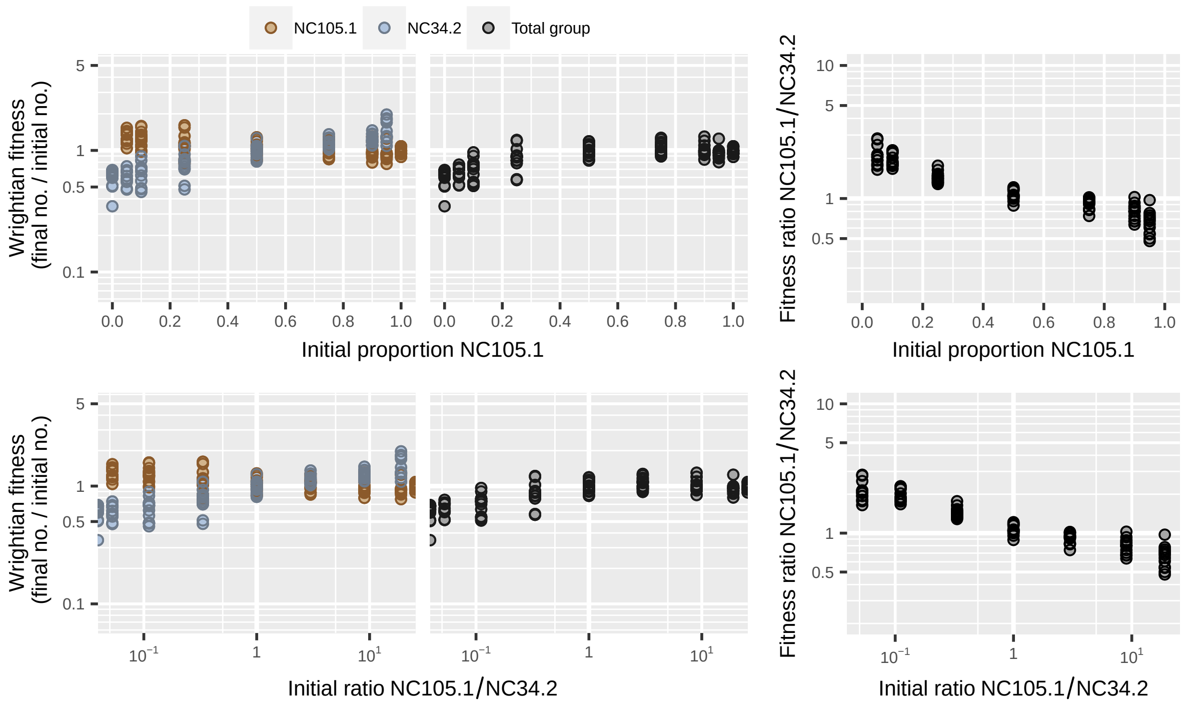

**Figure S40:** Dataset from Madgwick *et al.* [12] showing codevelopment of *Dictyostelium discoideum* strains NC105.1 and NC34.2.

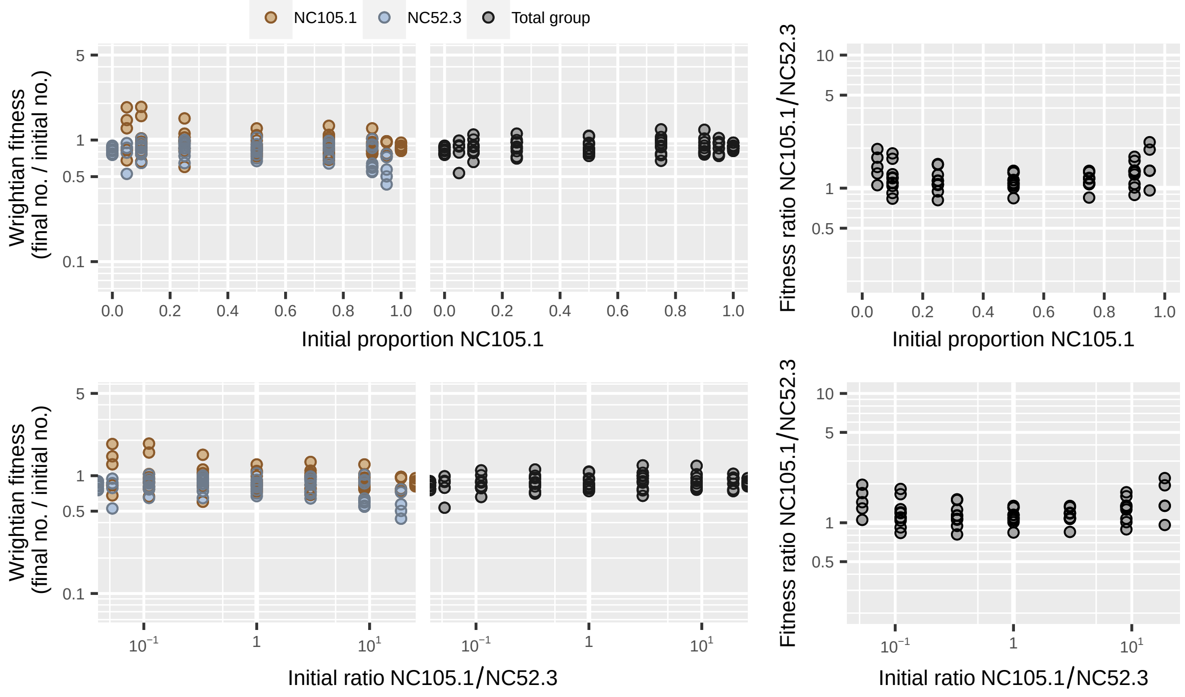

**Figure S41:** Dataset from Madgwick *et al.* [12] showing codevelopment of *Dictyostelium discoideum* strains NC105.1 and NC52.3.

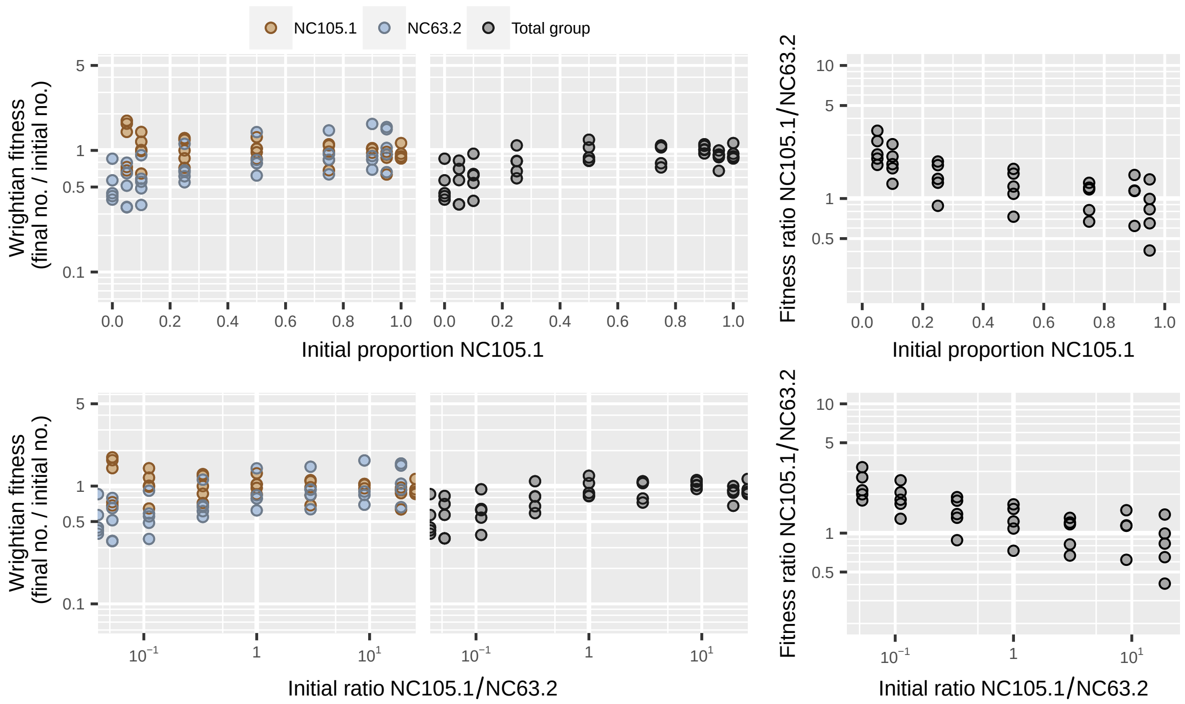

**Figure S42:** Dataset from Madgwick *et al.* [12] showing codevelopment of *Dictyostelium discoideum* strains NC105.1 and NC63.2.

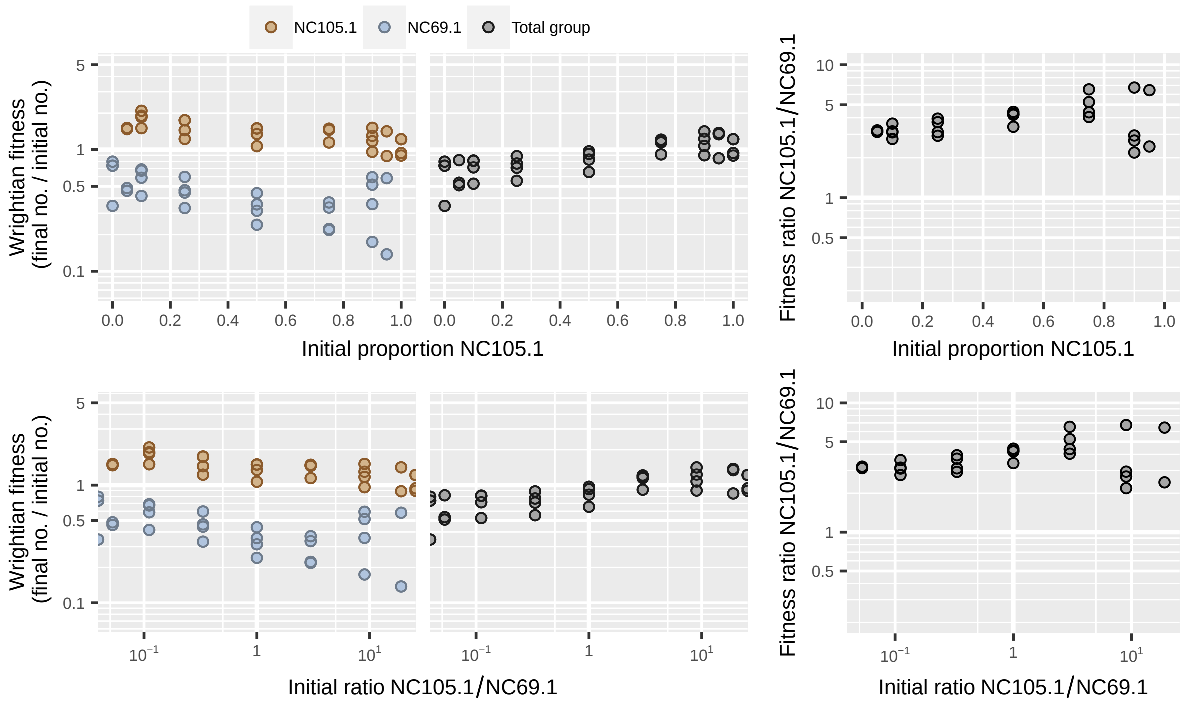

**Figure S43:** Dataset from Madgwick *et al.* [12] showing codevelopment of *Dictyostelium discoideum* strains NC105.1 and NC69.1.

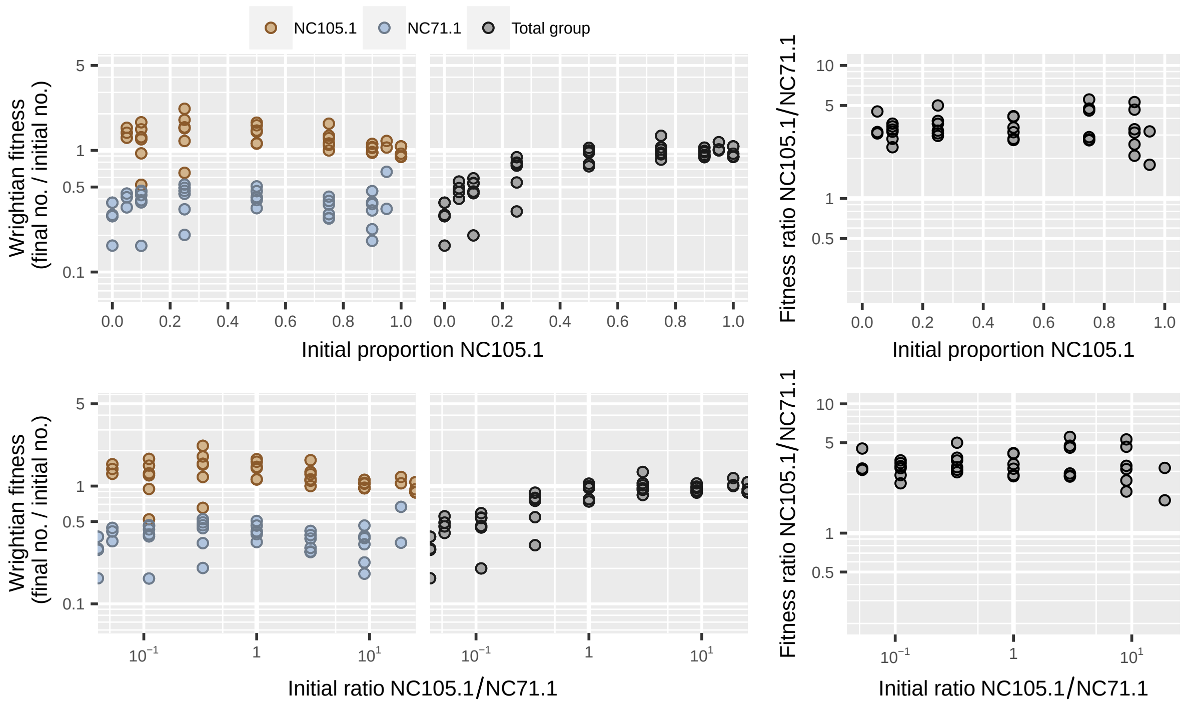

**Figure S44:** Dataset from Madgwick *et al.* [12] showing codevelopment of *Dictyostelium discoideum* strains NC105.1 and NC71.1.

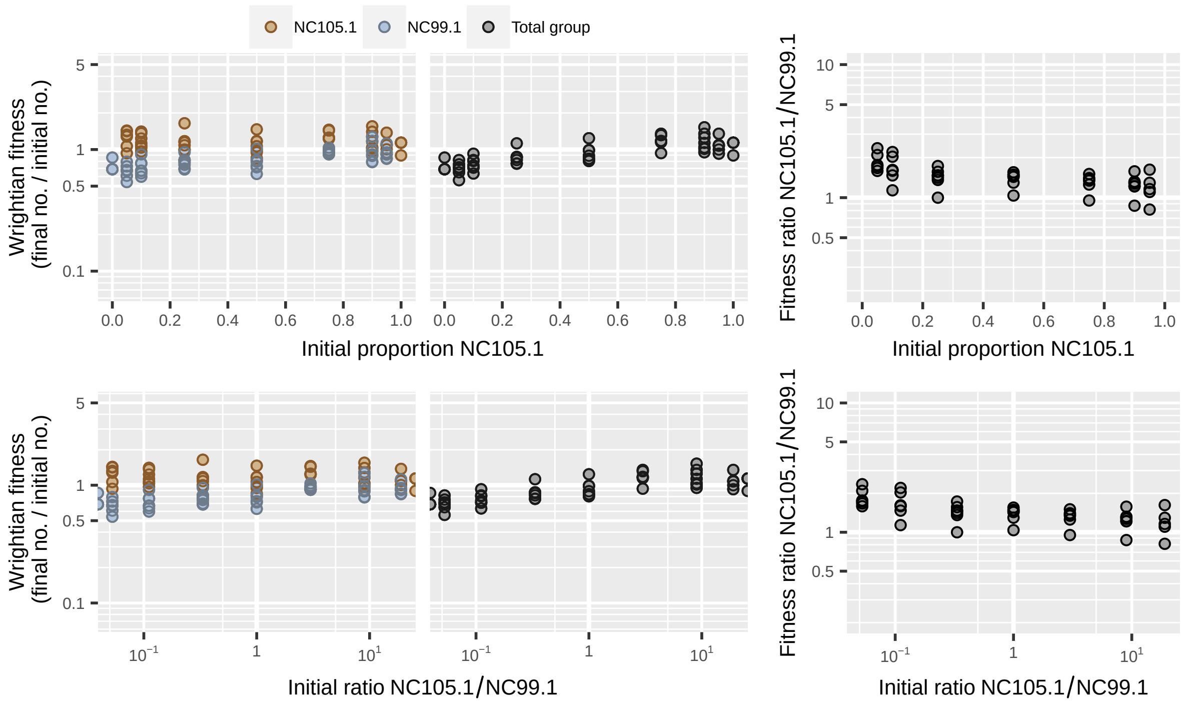

**Figure S45:** Dataset from Madgwick *et al.* [12] showing codevelopment of *Dictyostelium discoideum* strains NC105.1 and NC99.1.

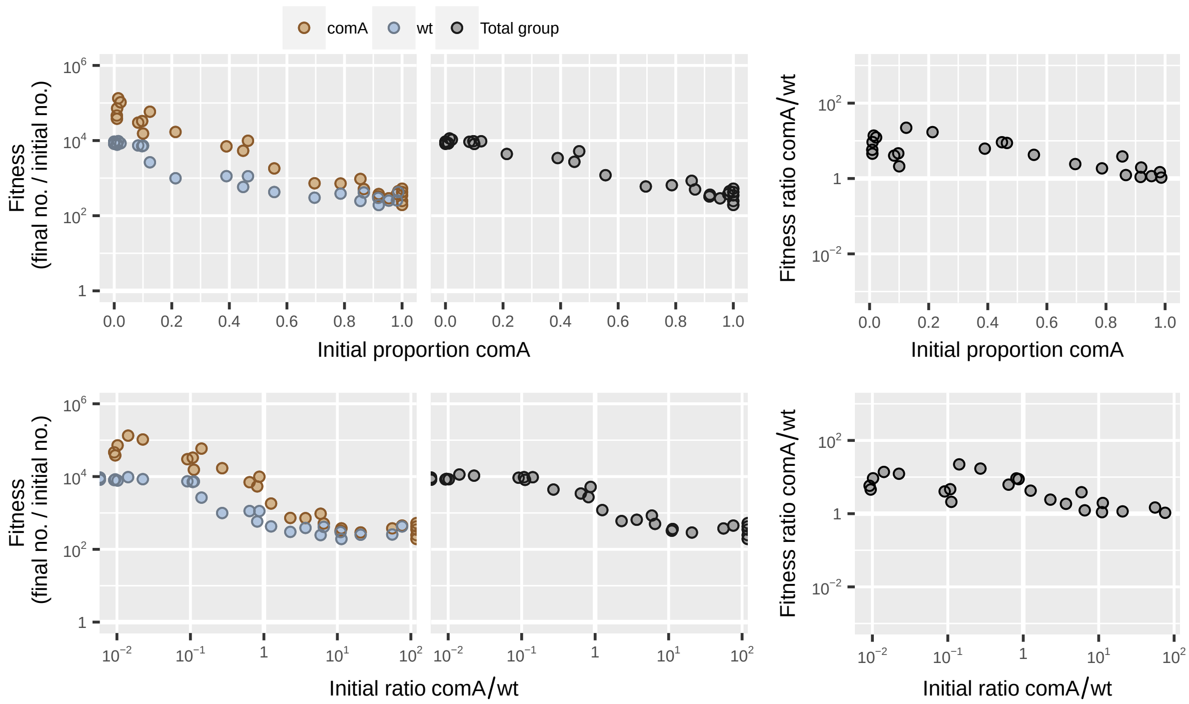

**Figure S46:** Dataset from Pollak *et al.* [13] showing mix experiment with *Bacillus subtilis* Δ*comA* mutant and wild-type on swarming media.

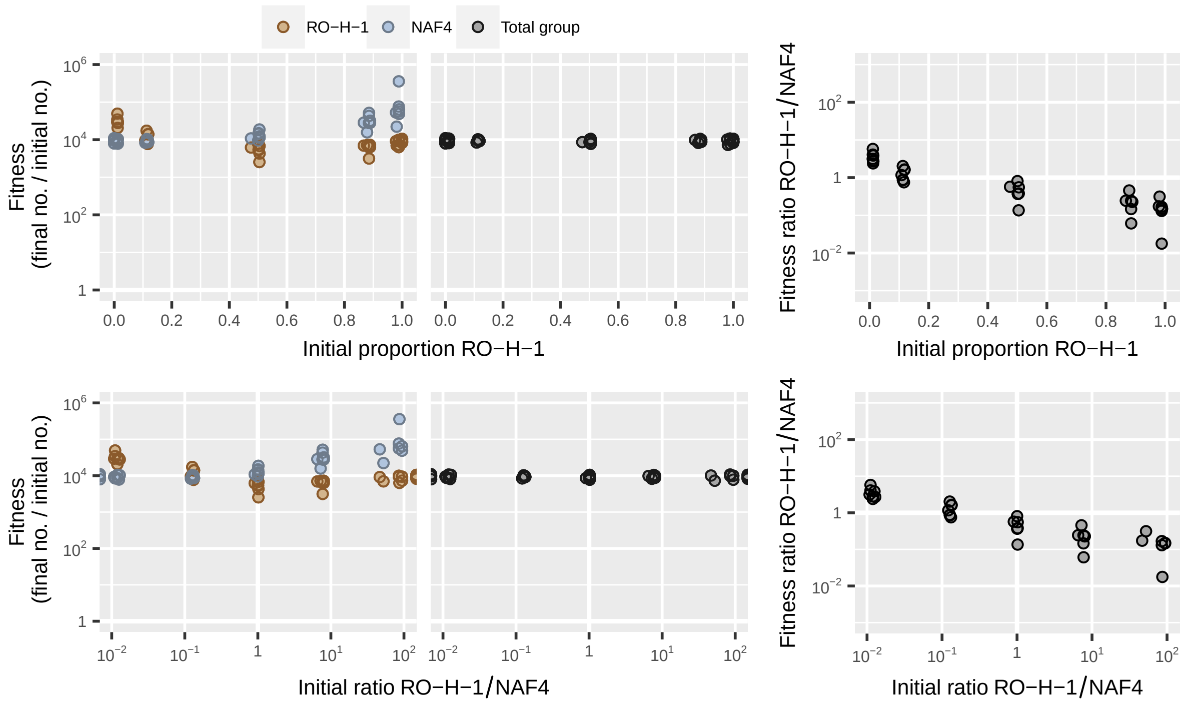

**Figure S47:** Dataset from Pollak *et al.* [13] showing mix experiment with *Bacillus subtilis* strains of different pherotypes on swarming media.

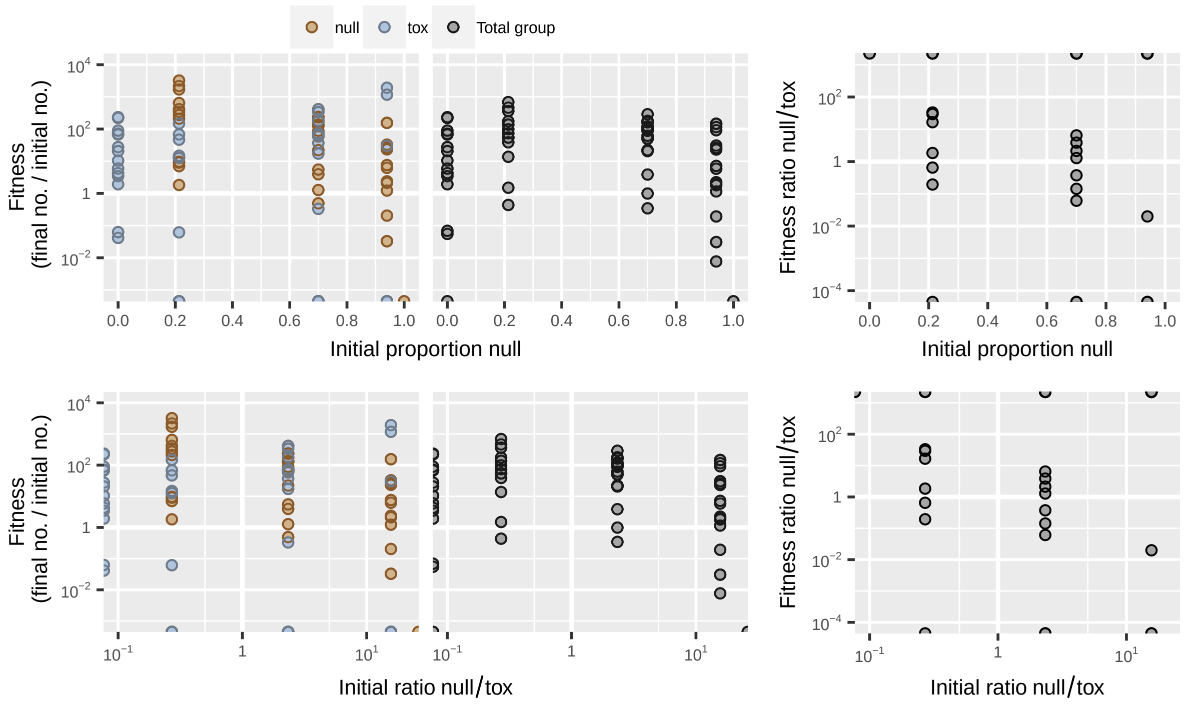

**Figure S48:** Dataset from Raymond *et al.* [14] showing coinfection of caterpillars with a wild-type *Bacillus thuringensis* strain and a mutant that does not produce Cry toxin.

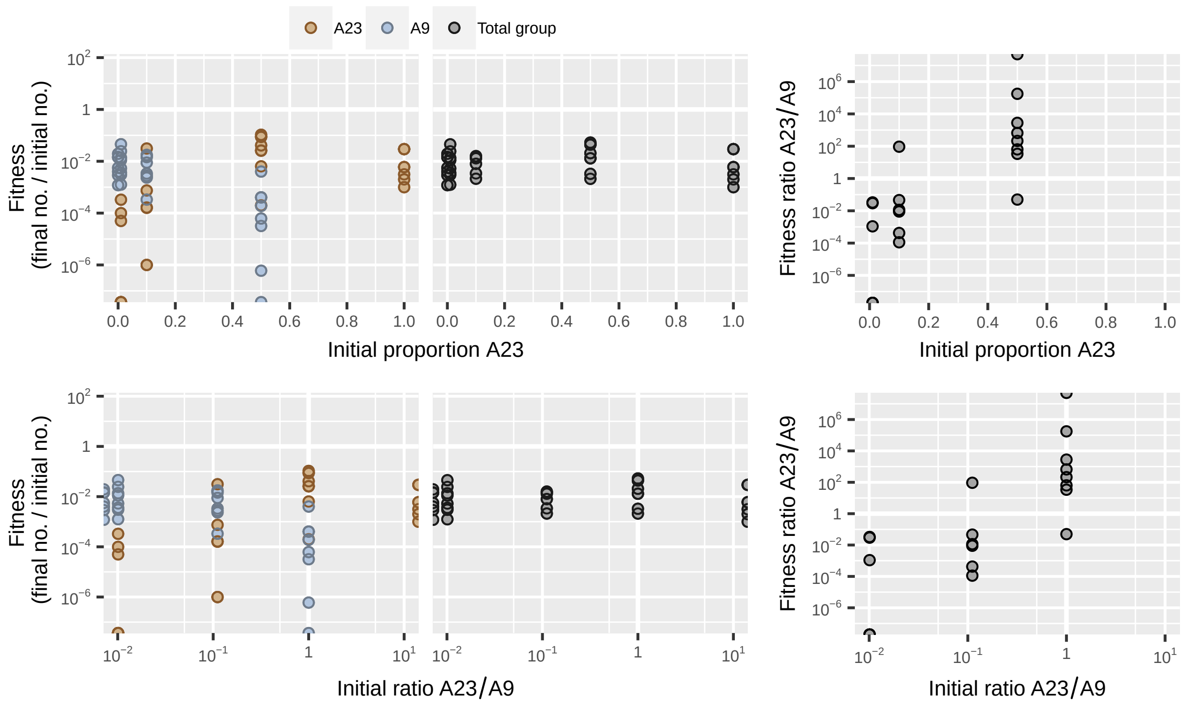

**Figure S49:** Dataset from Rendueles *et al.* [15] showing codevelopment of *Myxococcus xanthus* strains A23 and A9 in which there is contact-dependent interference competition.

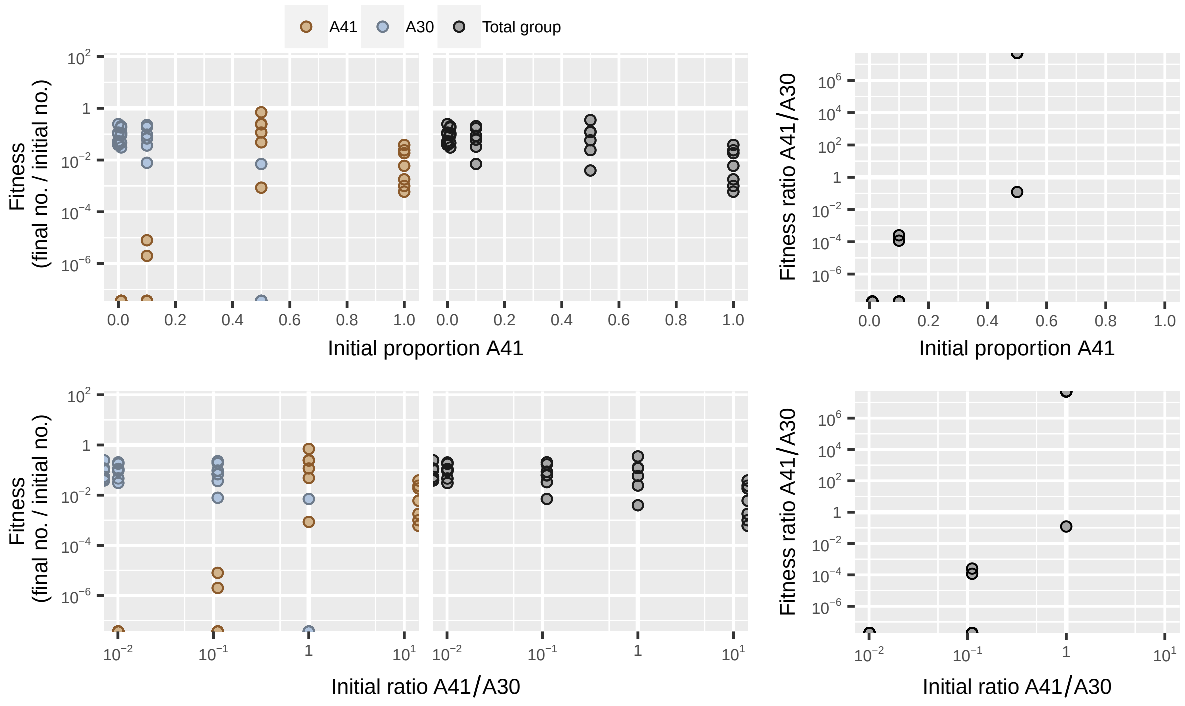

**Figure S50:** Dataset from Rendueles *et al.* [15] showing codevelopment of *Myxococcus xanthus* strains A41 and A30 in which there is contact-dependent interference competition.

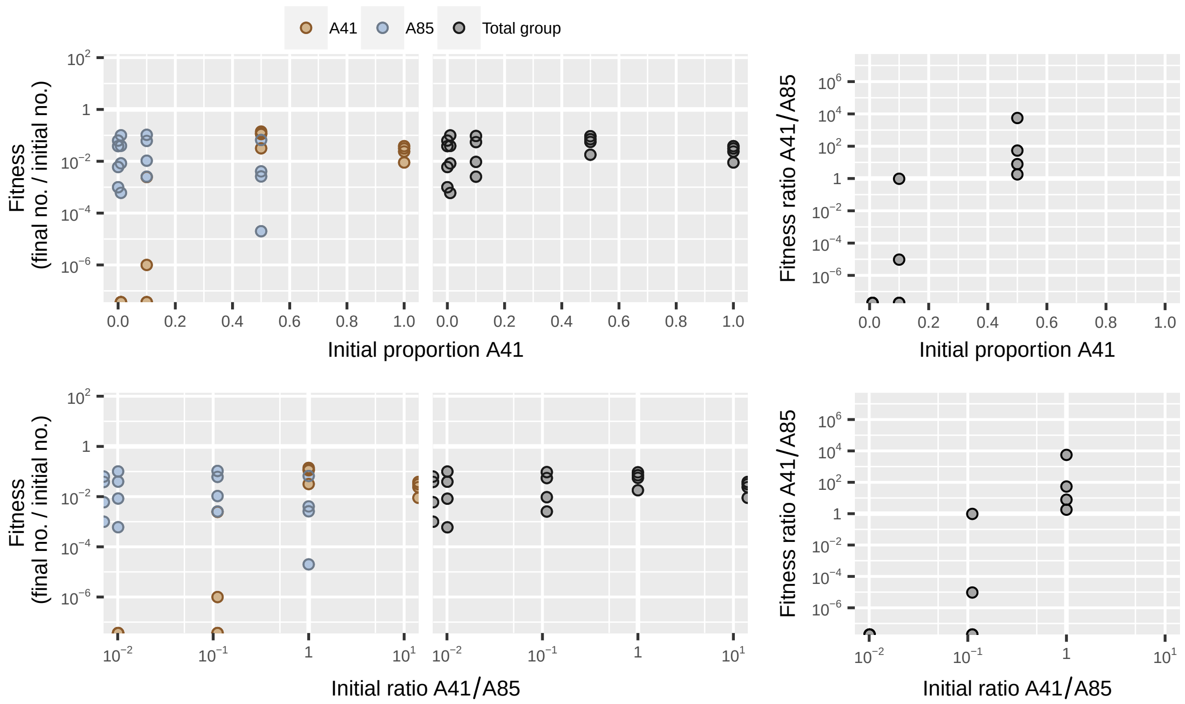

**Figure S51:** Dataset from Rendueles *et al.* [15] showing codevelopment of *Myxococcus xanthus* strains A41 and A85 in which there is contact-dependent interference competition.

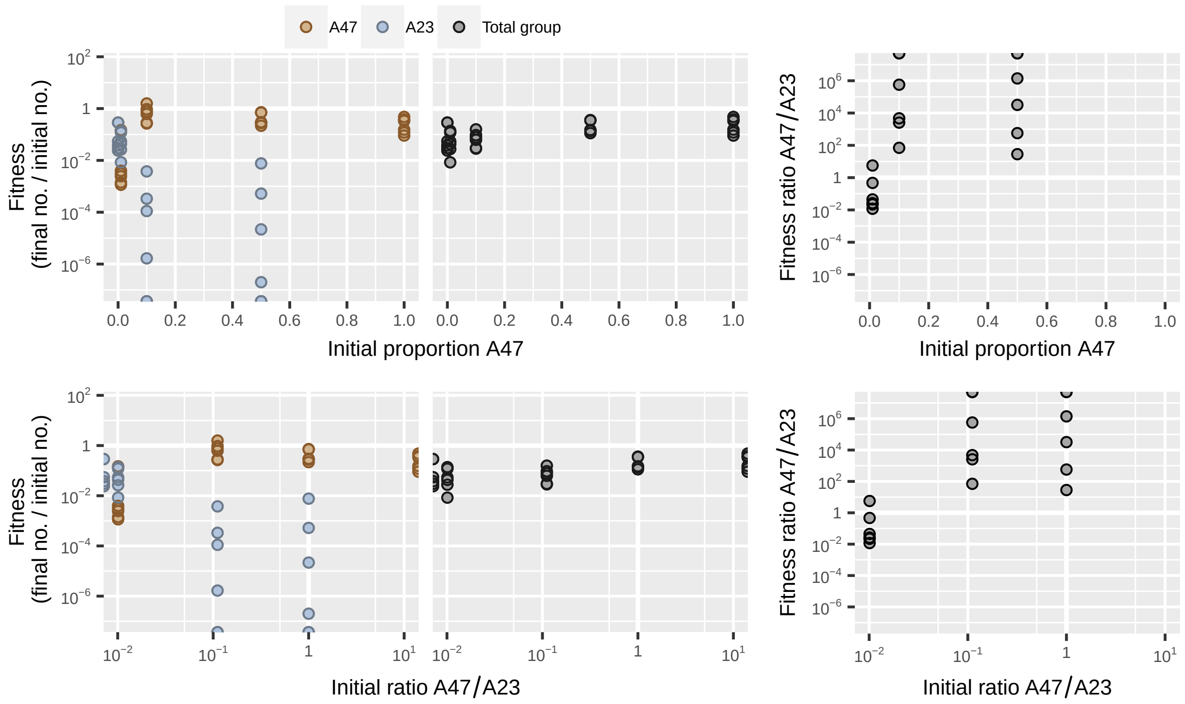

**Figure S52:** Dataset from Rendueles *et al.* [15] showing codevelopment of *Myxococcus xanthus* strains A47 and A23 in which there is contact-dependent interference competition.

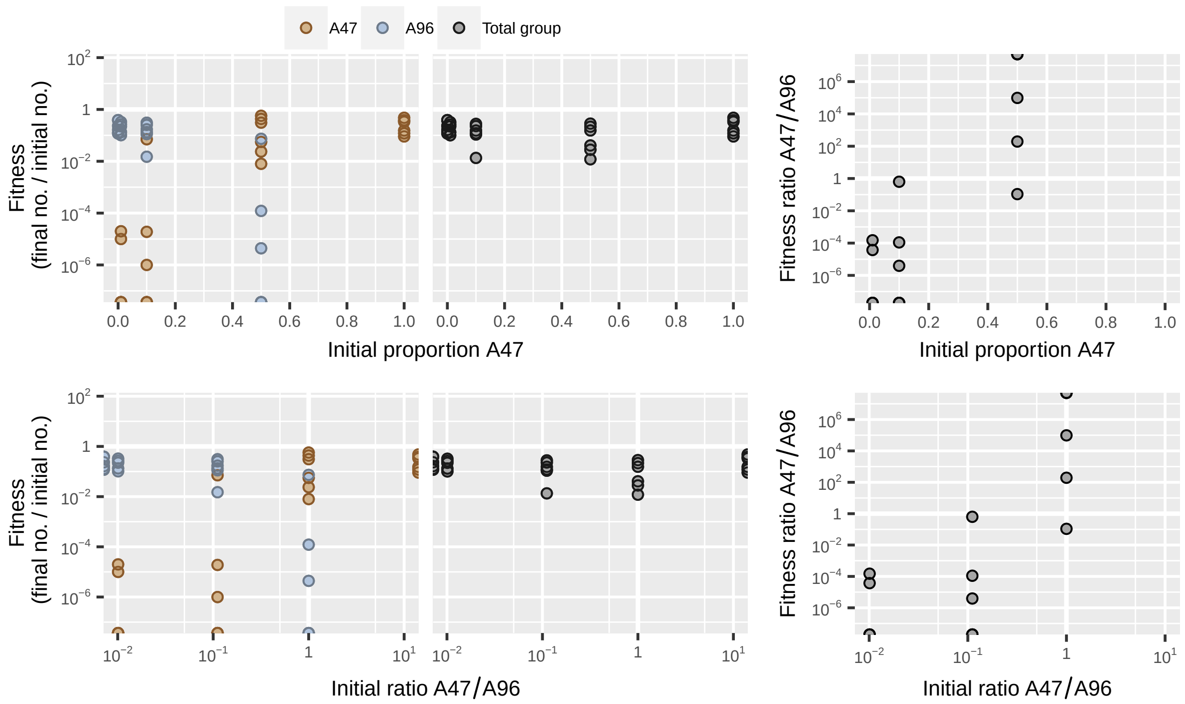

**Figure S53:** Dataset from Rendueles *et al.* [15] showing codevelopment of *Myxococcus xanthus* strains A47 and A96 in which there is contact-dependent interference competition.

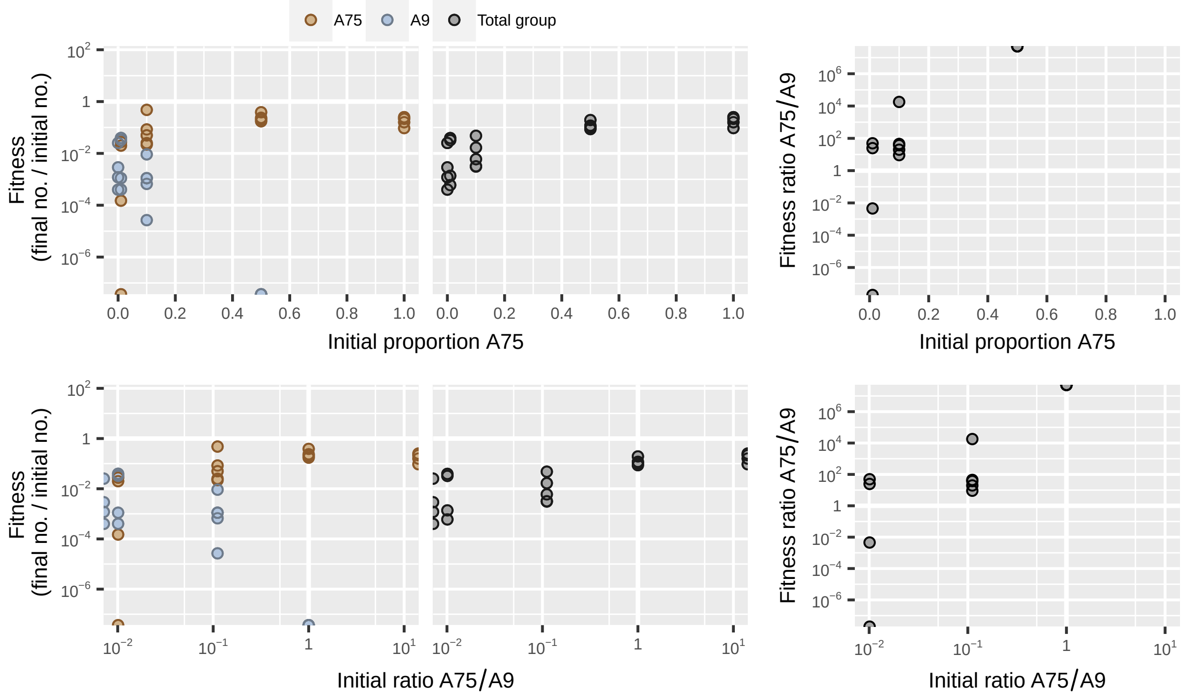

**Figure S54:** Dataset from Rendueles *et al.* [15] showing codevelopment of *Myxococcus xanthus* strains A75 and A9 in which there is contact-dependent interference competition.

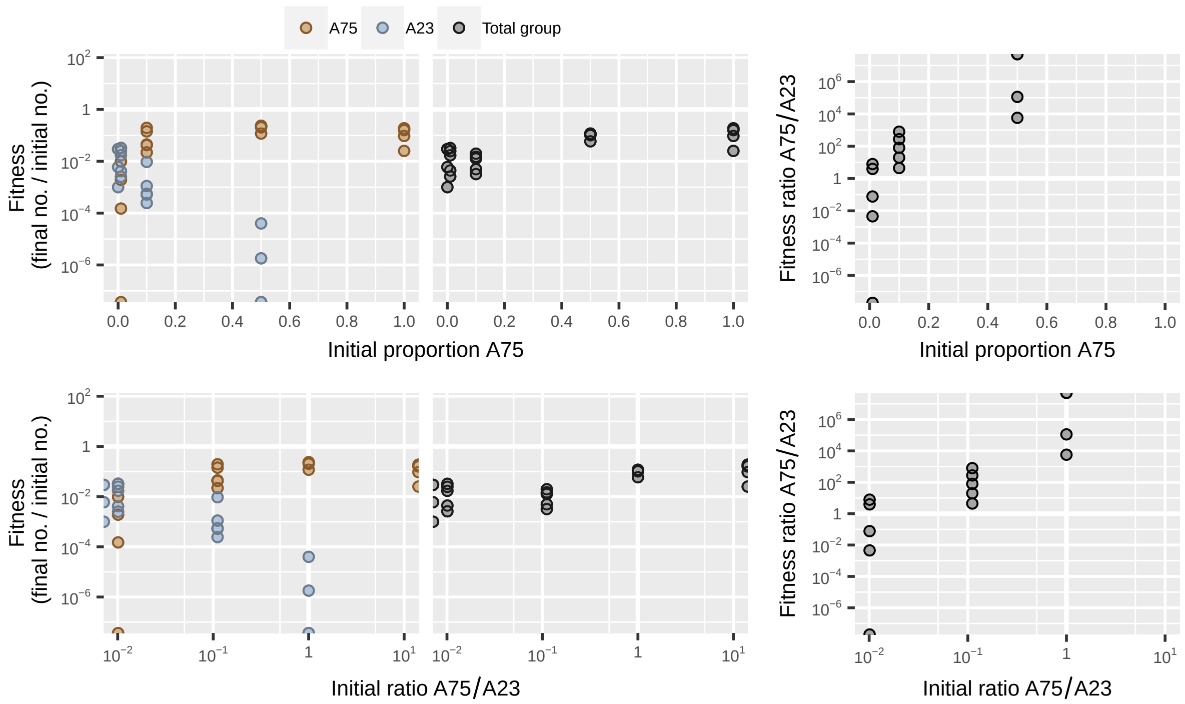

**Figure S55:** Dataset from Rendueles *et al.* [15] showing codevelopment of *Myxococcus xanthus* strains A75 and A23 in which there is contact-dependent interference competition.

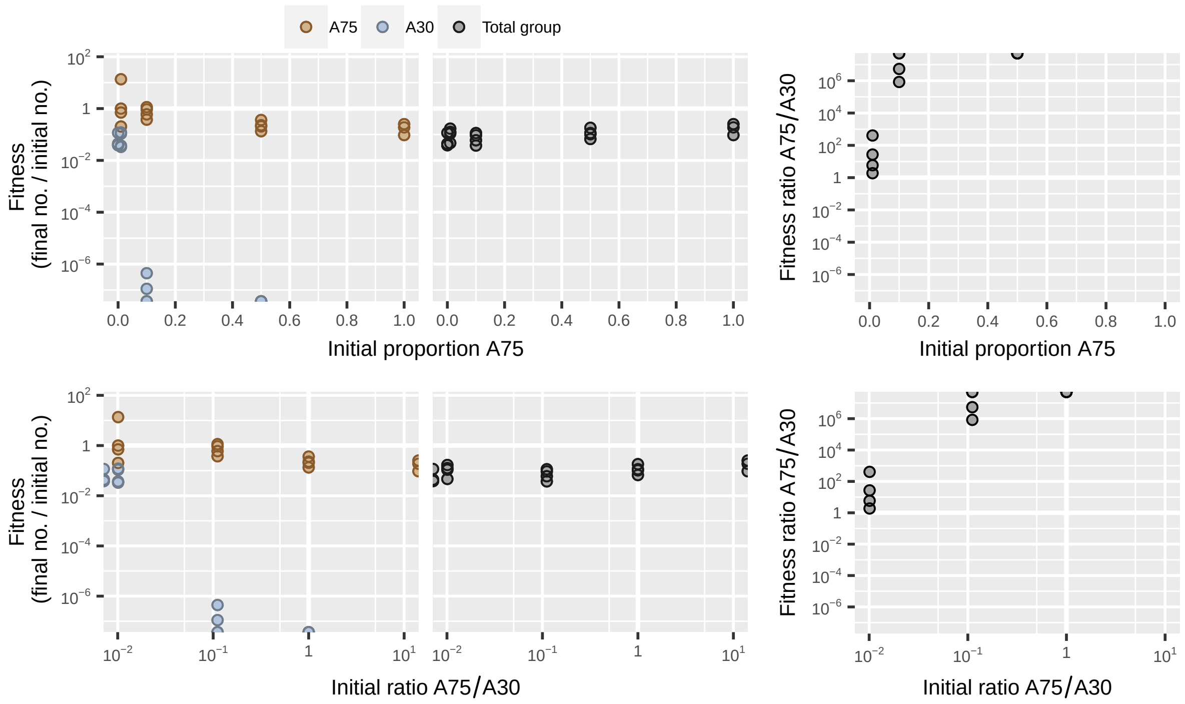

**Figure S56:** Dataset from Rendueles *et al.* [15] showing codevelopment of *Myxococcus xanthus* strains A75 and A30 in which there is contact-dependent interference competition.

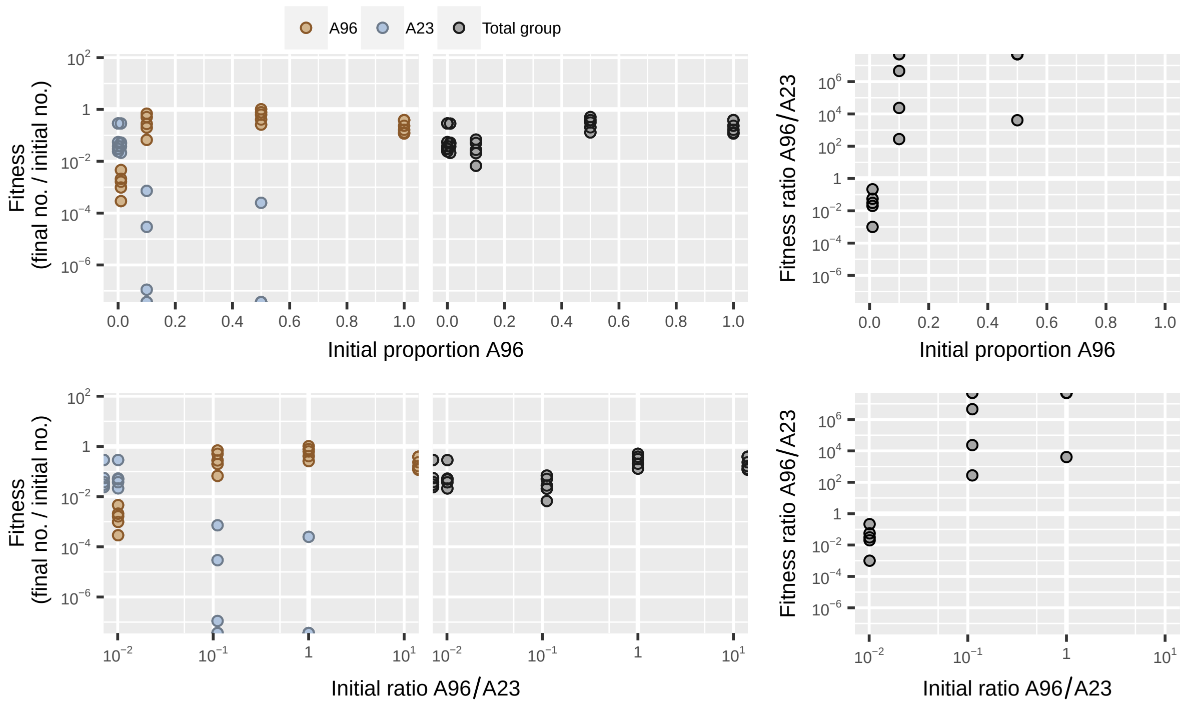

**Figure S57:** Dataset from Rendueles *et al.* [15] showing codevelopment of *Myxococcus xanthus* strains A96 and A23 in which there is contact-dependent interference competition.

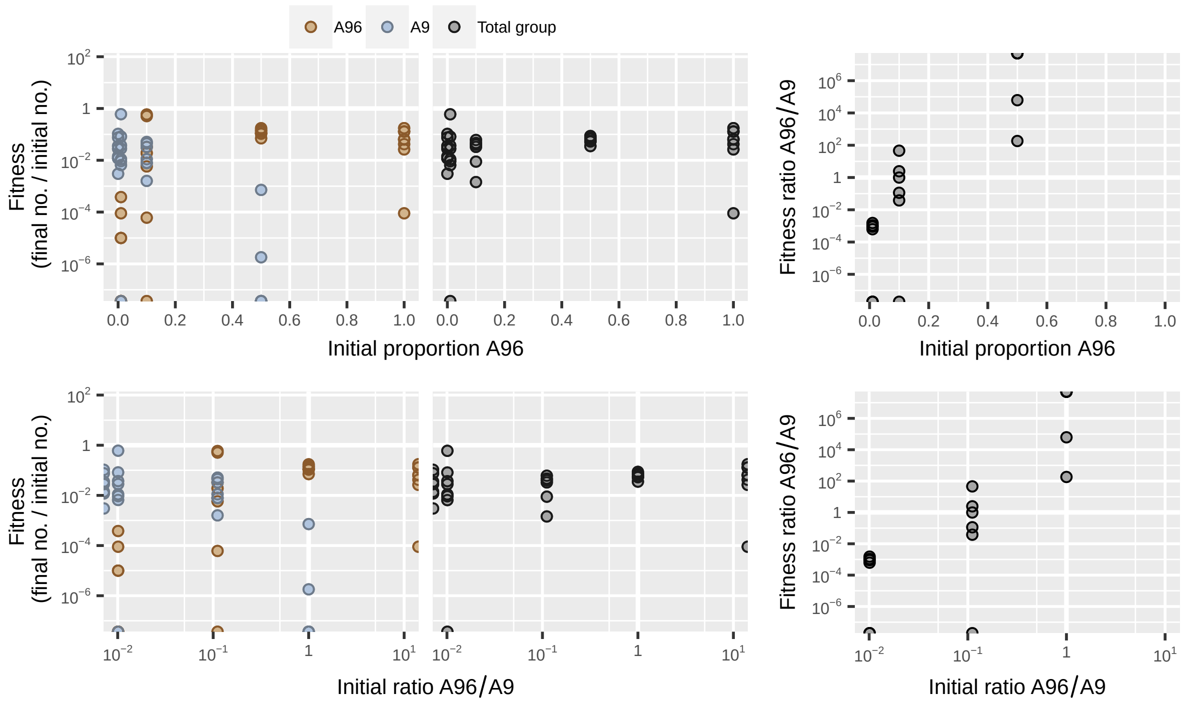

**Figure S58:** Dataset from Rendueles *et al.* [15] showing codevelopment of *Myxococcus xanthus* strains A96 and A9 in which there is contact-dependent interference competition.

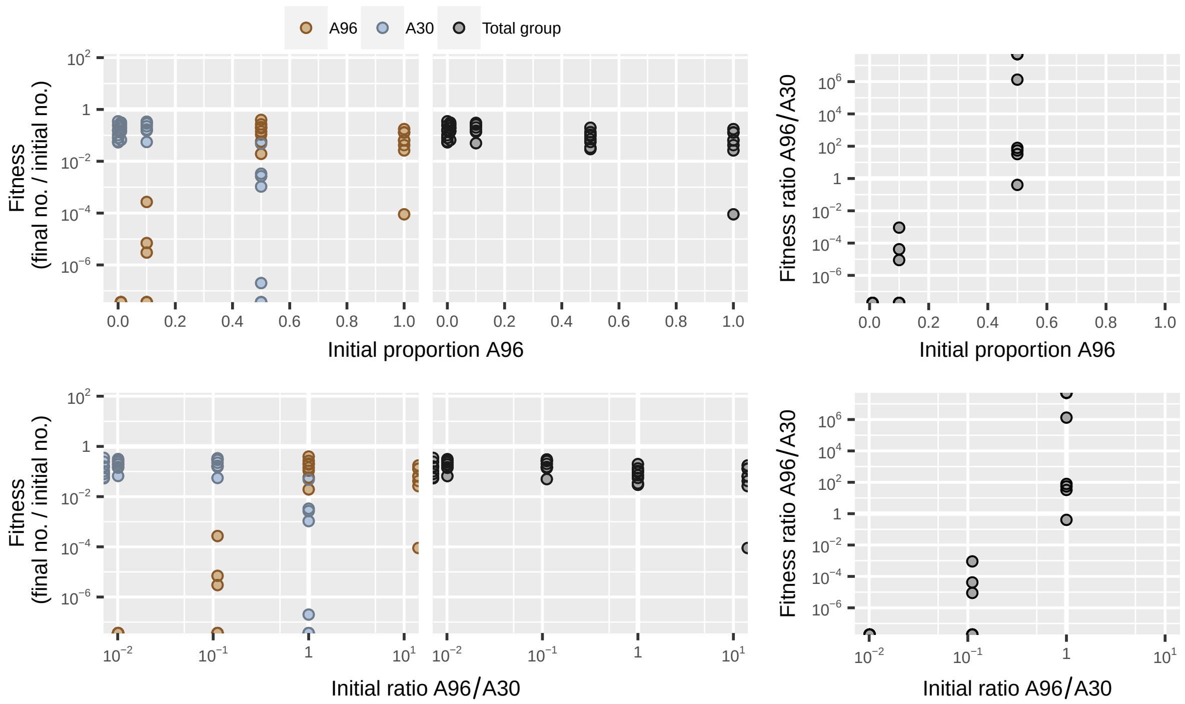

**Figure S59:** Dataset from Rendueles *et al.* [15] showing codevelopment of *Myxococcus xanthus* strains A96 and A30 in which there is contact-dependent interference competition.

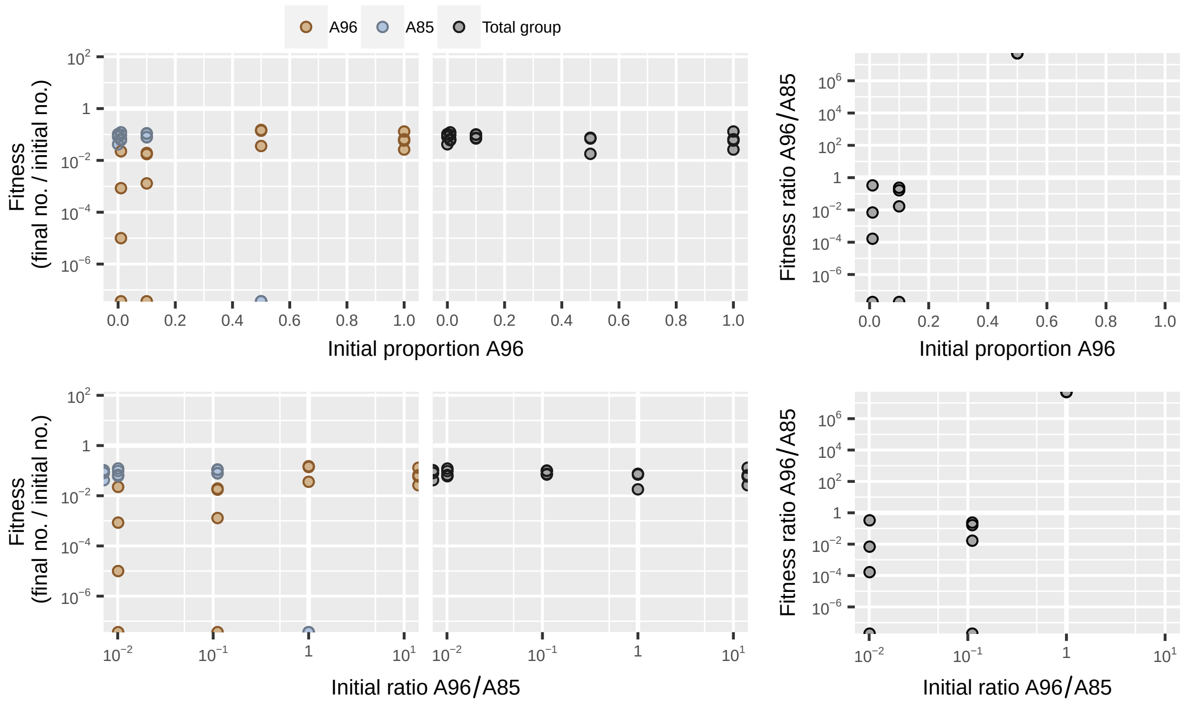

**Figure S60:** Dataset from Rendueles *et al.* [15] showing codevelopment of *Myxococcus xanthus* strains A96 and A85 in which there is contact-dependent interference competition.

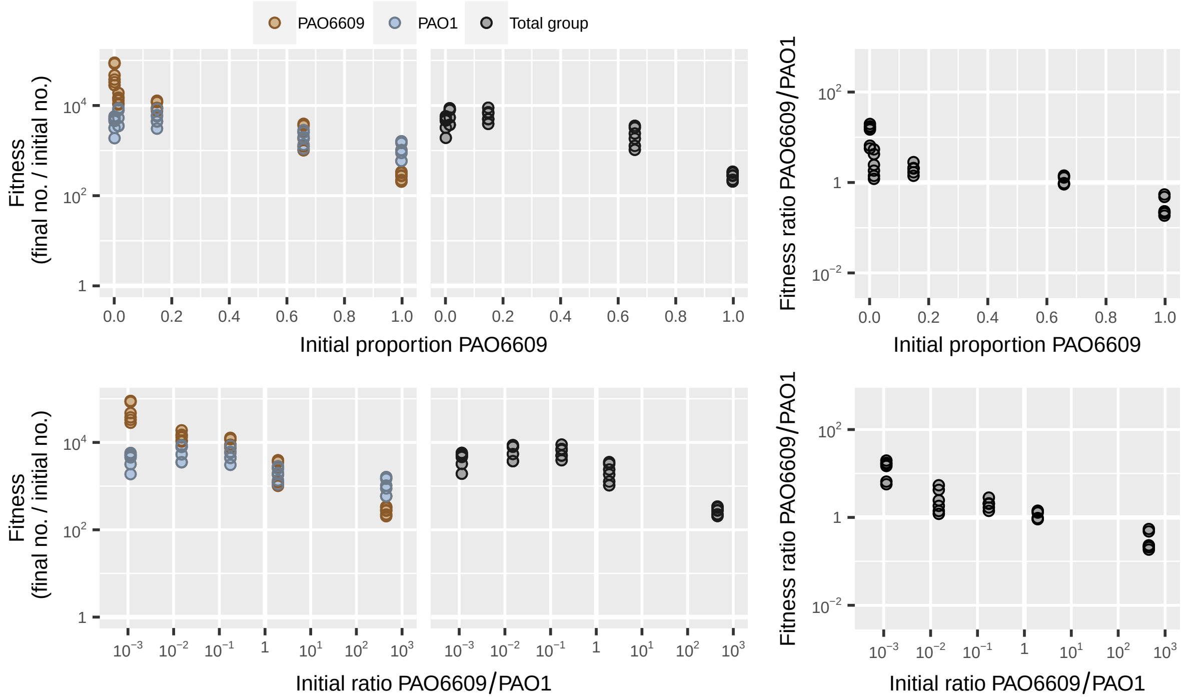

**Figure S61:** Dataset from Ross-Gillespie *et al.* [16] showing mix experiment with wild-type *Pseudomonas aeruginosa* strain PAO1 and pyoverdin nonproducer PAO6609 in iron-limited media.

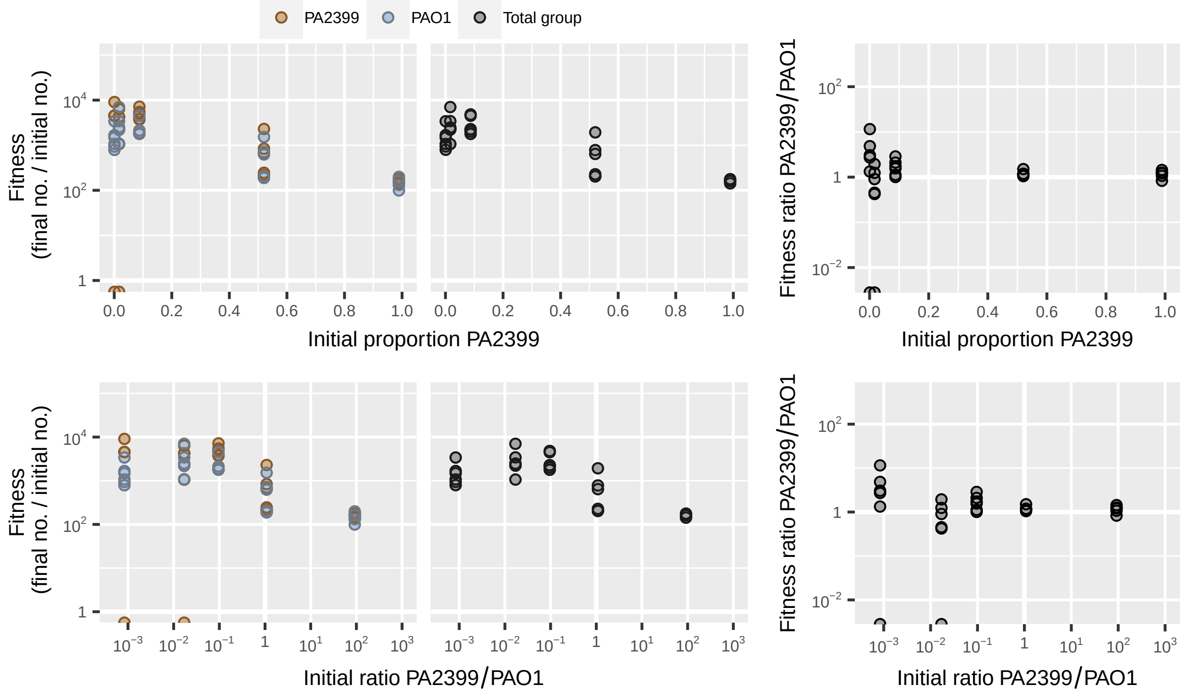

**Figure S62:** Dataset from Ross-Gillespie *et al.* [16] showing mix experiment with wild-type *Pseudomonas aeruginosa* strain PAO1 and pyoverdin nonproducer PA2399 in iron-limited media.

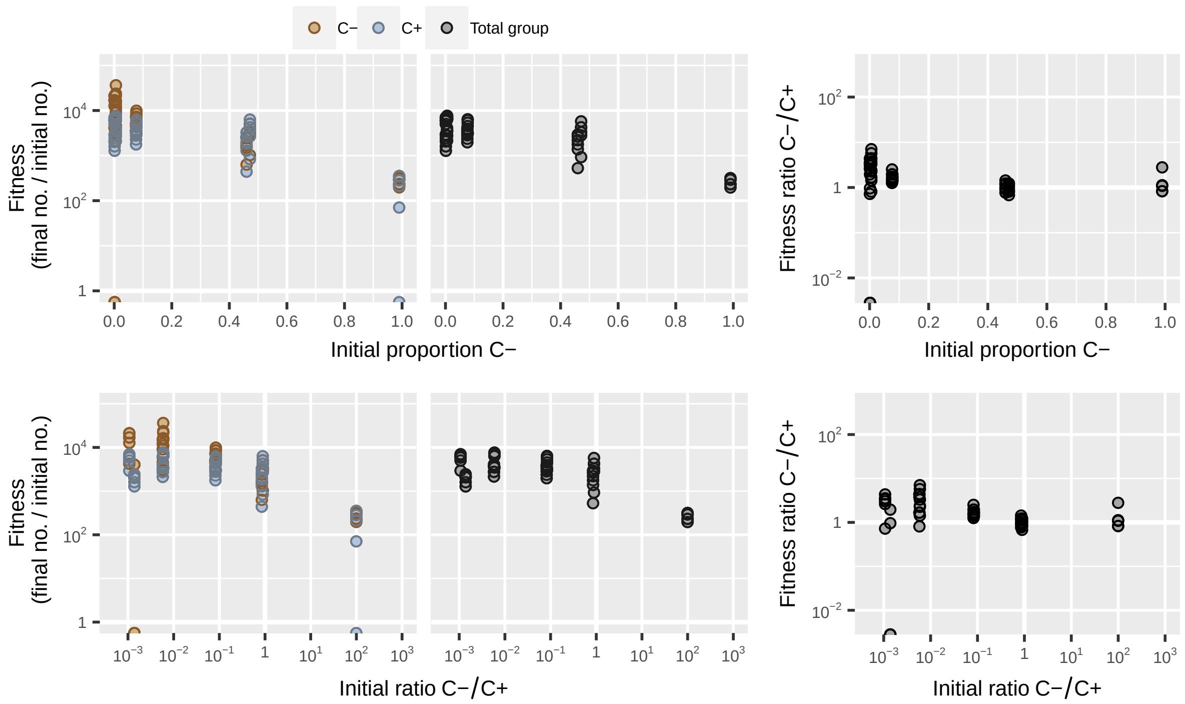

**Figure S63:** Dataset from Ross-Gillespie *et al.* [16] showing mix experimentwith wild-type *Pseudomonas aeruginosa* strain C+ and pyoverdin nonproducer C- in iron-limited media.

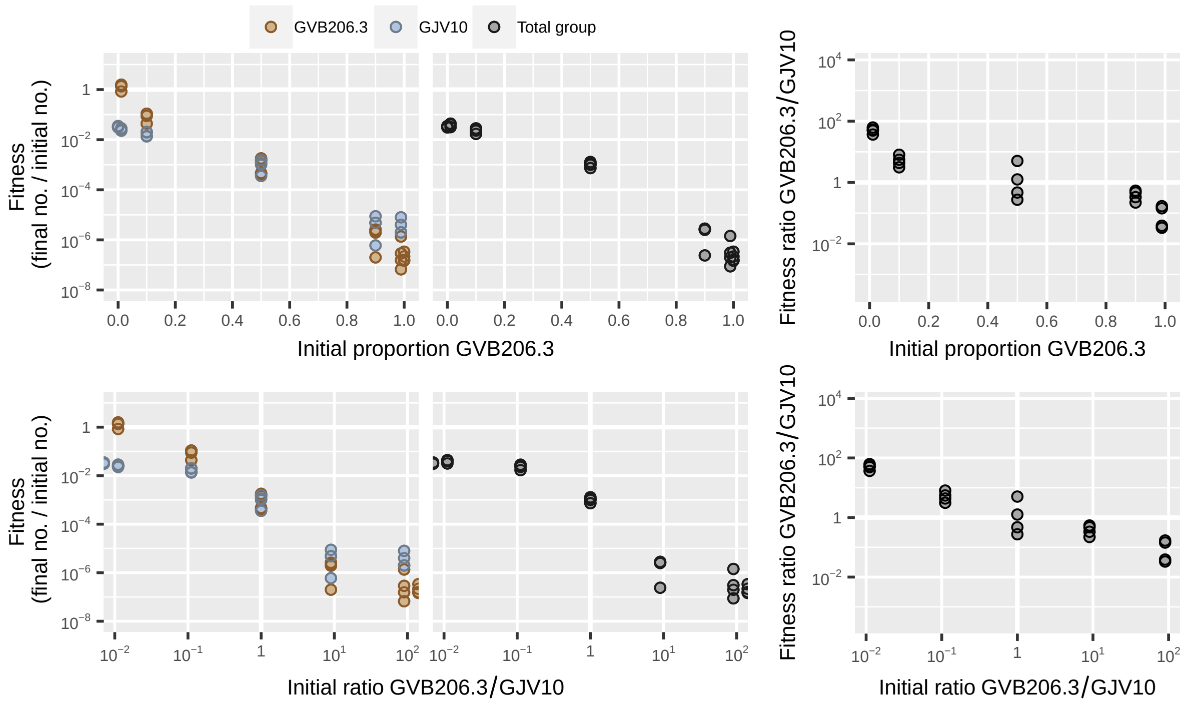

**Figure S64:** Dataset from smith *et al.* [17] showing codevelopment of wild-type *Myxococcus xanthus* GJV2 and laboratory-evolved cheater GJV206.3.

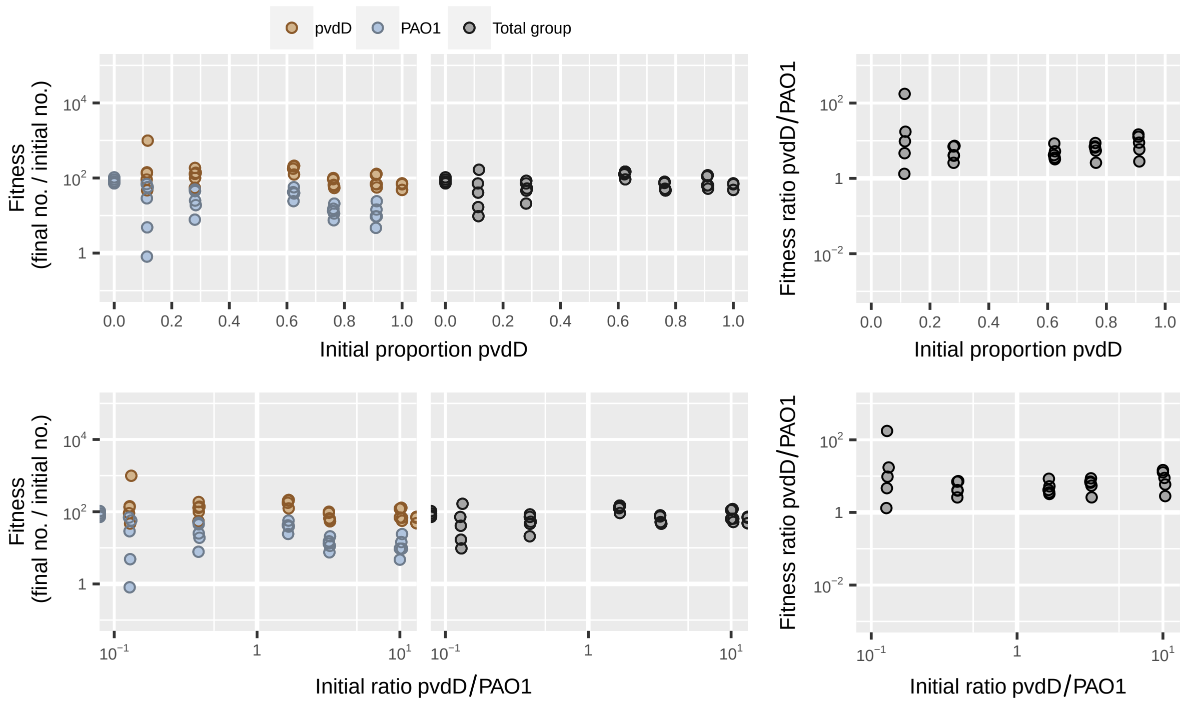

**Figure S65:** Dataset from Vasse *et al.* [18] showing mix experiment with wild-type *Pseudomonas aeruginosa* strain PAO1 and siderophore nonproducer PAO1ΔpvdD in iron-limited media without phage.

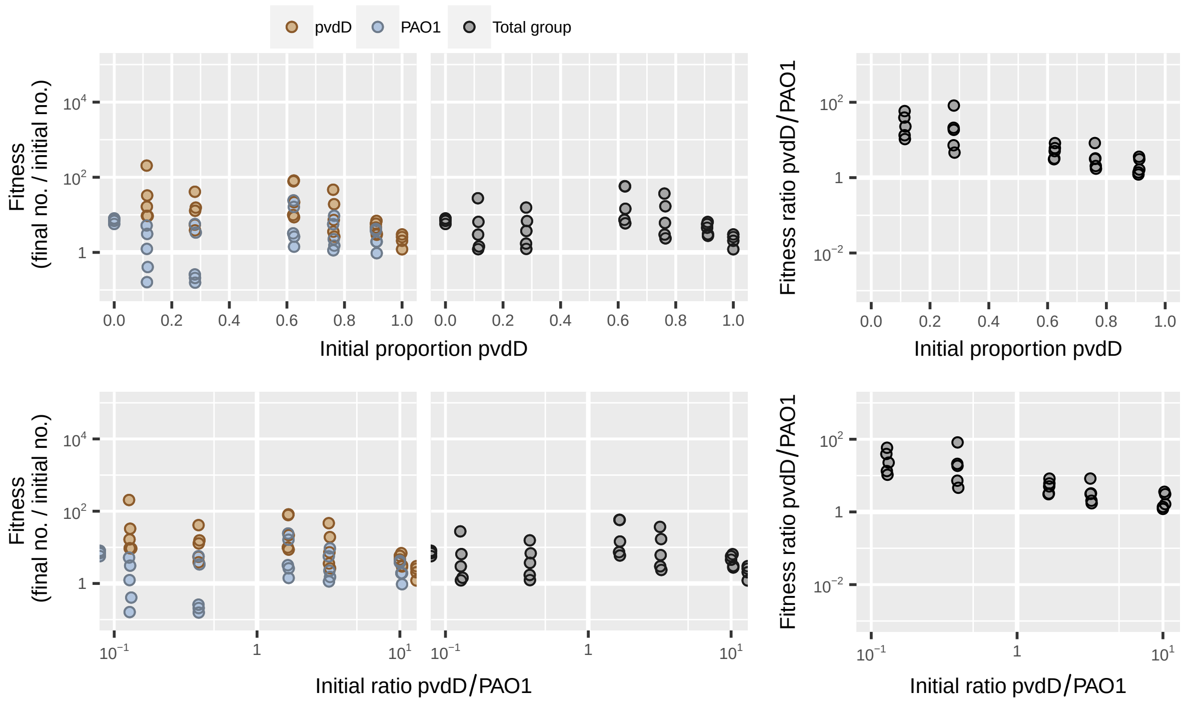

**Figure S66:** Dataset from Vasse *et al.* [18] showing mix experiment with wild-type *Pseudomonas aeruginosa* strain PAO1 and siderophore nonproducer PAO1ΔpvdD in iron-limited media with phage.

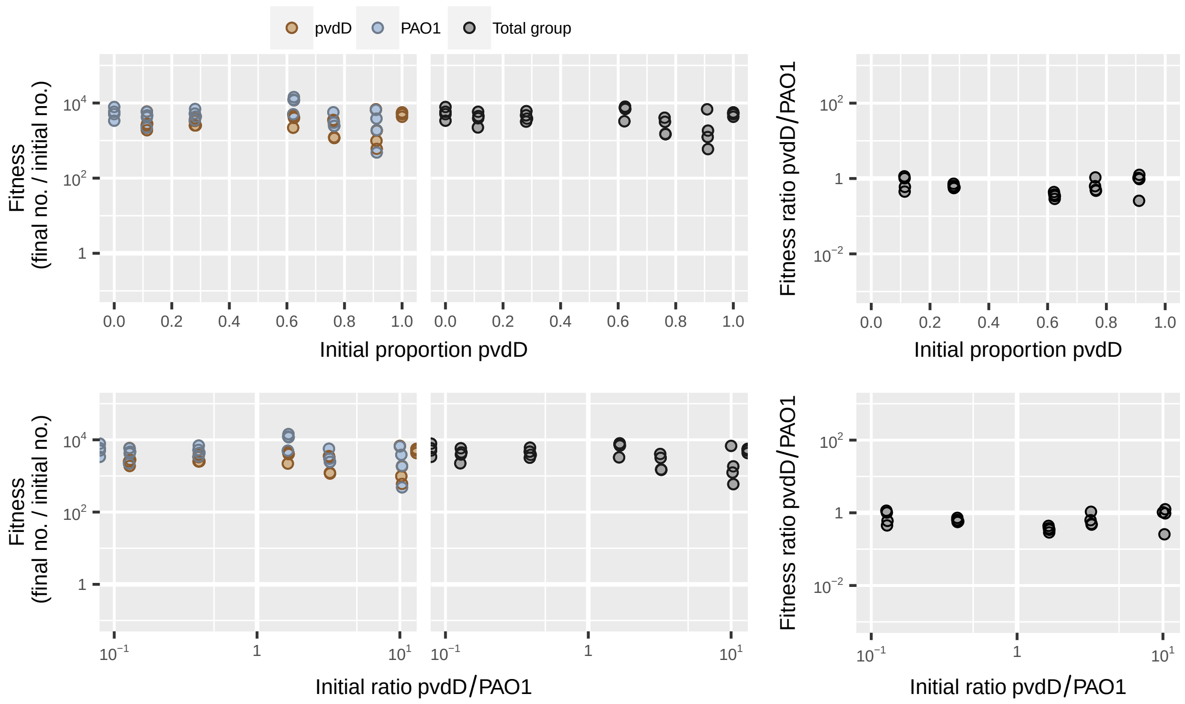

**Figure S67:** Dataset from Vasse *et al.* [18] showing mix experiment with wild-type *Pseudomonas aeruginosa* strain PAO1 and siderophore nonproducer PAO1ΔpvdD in iron-supplemented media without phage.

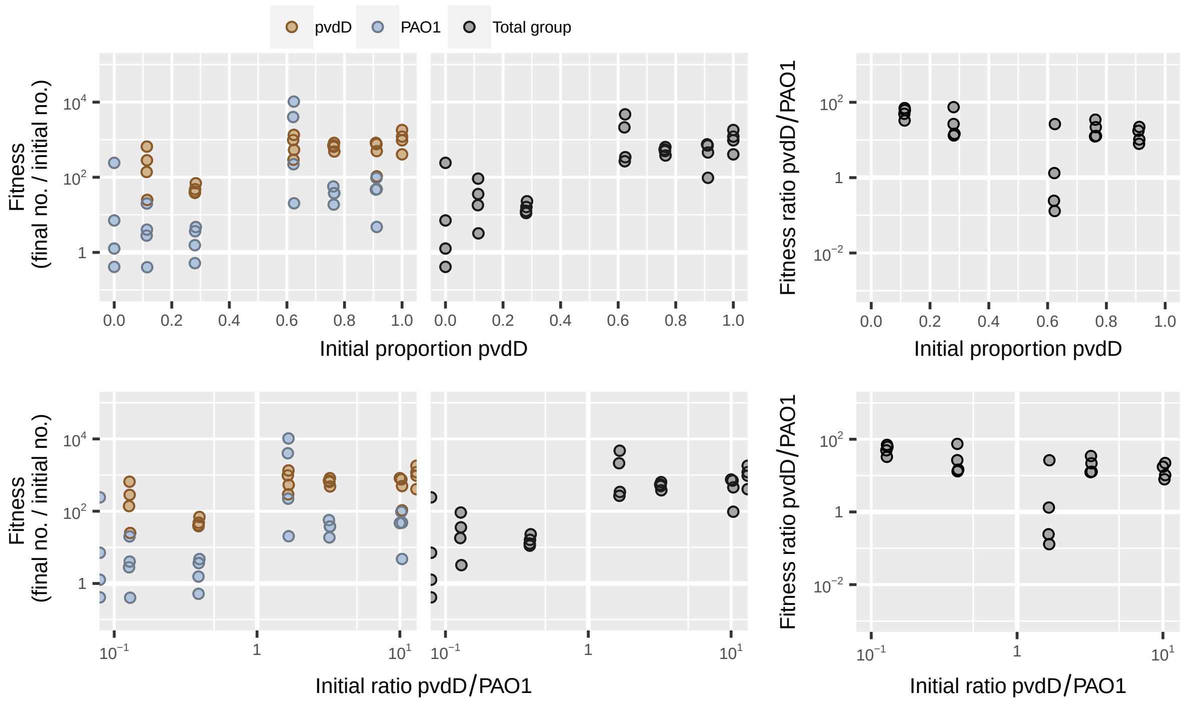

**Figure S68:** Dataset from Vasse *et al.* [18] showing mix experiment with wild-type *Pseudomonas aeruginosa* strain PAO1 and siderophore nonproducer PAO1ΔpvdD in iron-supplemented media with phage.

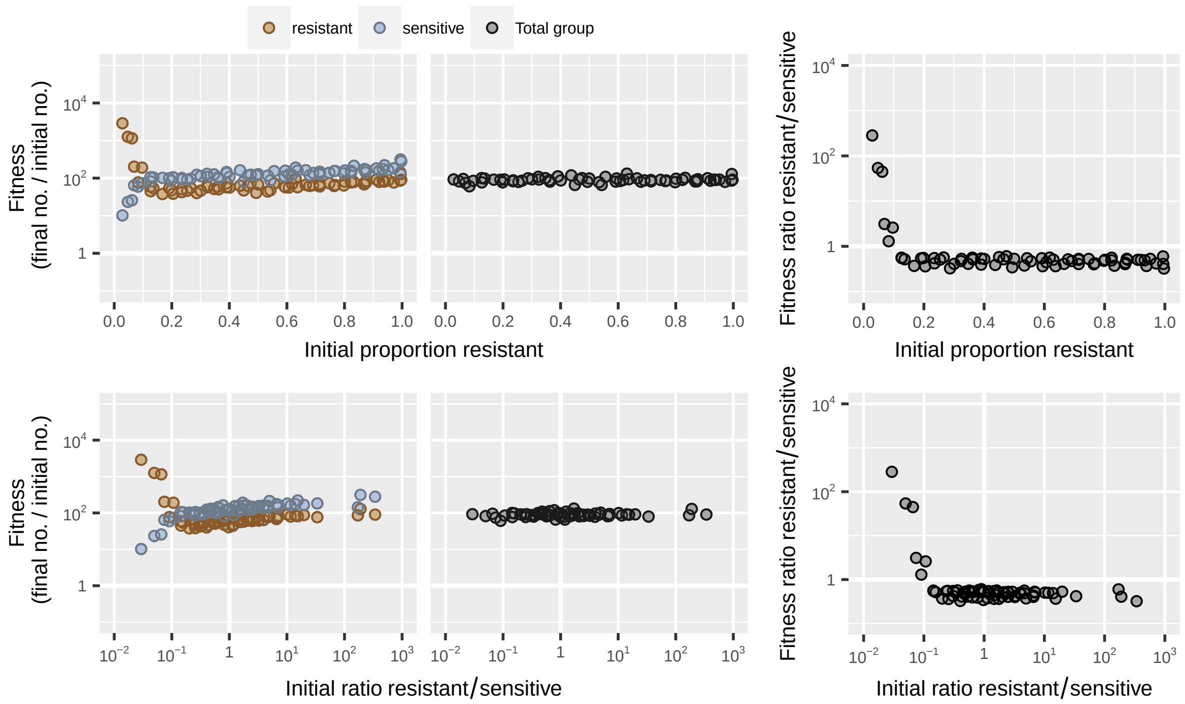

**Figure S69:** Dataset from Yurtsev *et al.* [19] showing coculture of antibiotic-resistant and sensitive *Escherichia coli* in media containing 15 $\mu$g/ml ampicillin with 100-fold dilution.

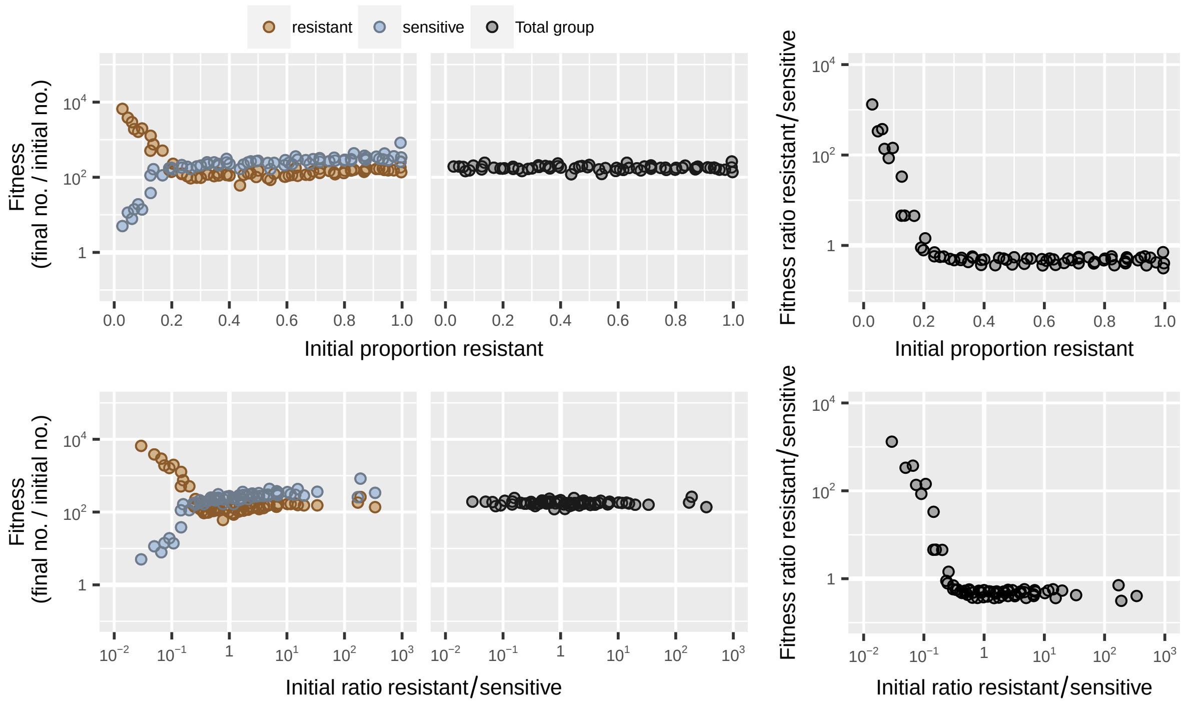

**Figure S70:** Dataset from Yurtsev *et al.* [19] showing coculture of antibiotic-resistant and sensitive *Escherichia coli* in media containing 15 $\mu$g/ml ampicillin with 200-fold dilution.

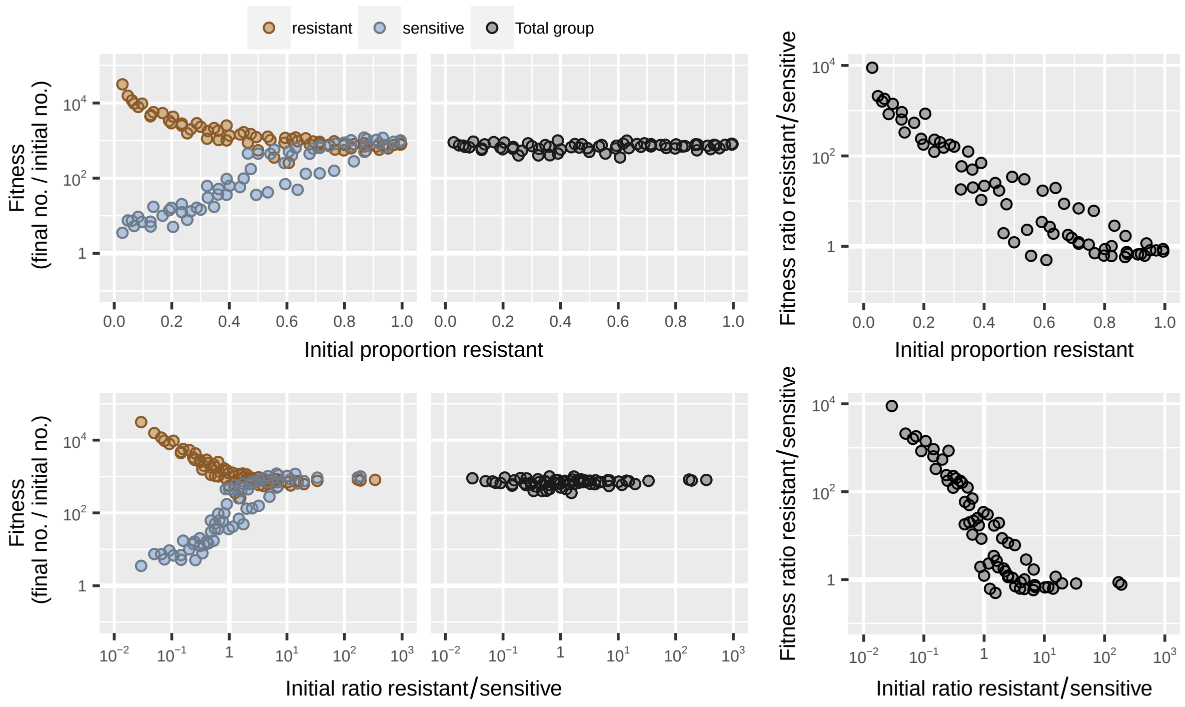

**Figure S71:** Dataset from Yurtsev *et al.* [19] showing coculture of antibiotic-resistant and sensitive *Escherichia coli* in media containing 15 μg/ml ampicillin with 800-fold dilution.

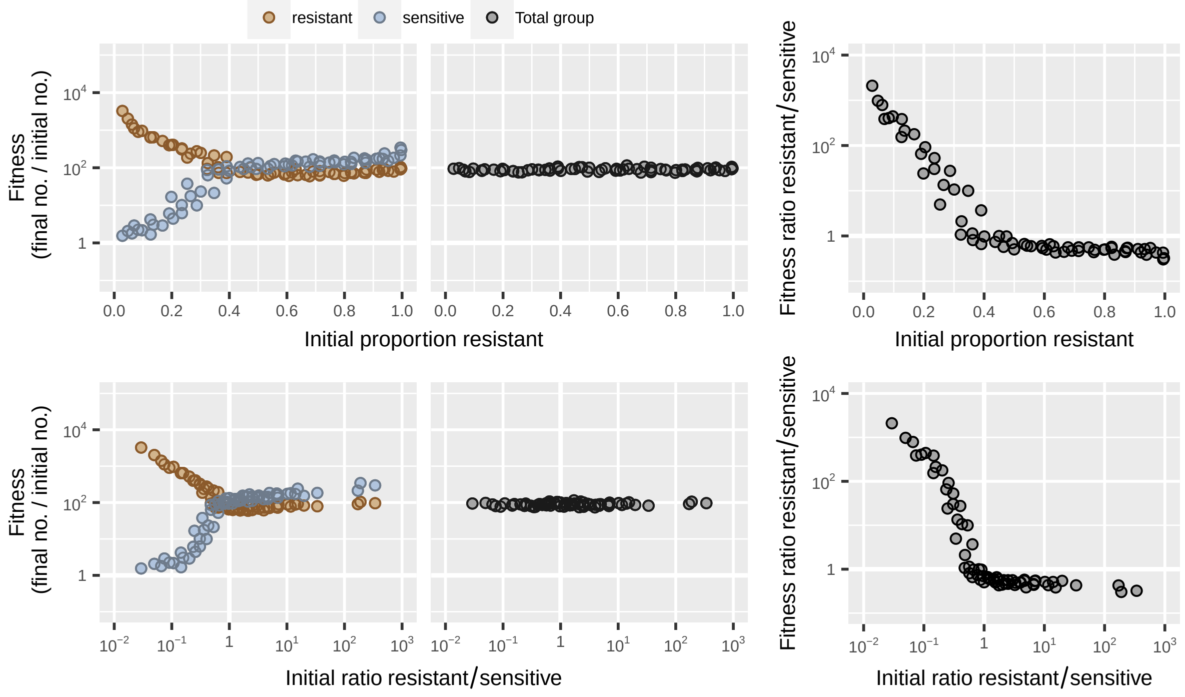

**Figure S72:** Dataset from Yurtsev *et al.* [19] showing coculture of antibiotic-resistant and sensitive *Escherichia coli* in media containing 100 μg/ml ampicillin with 100-fold dilution.

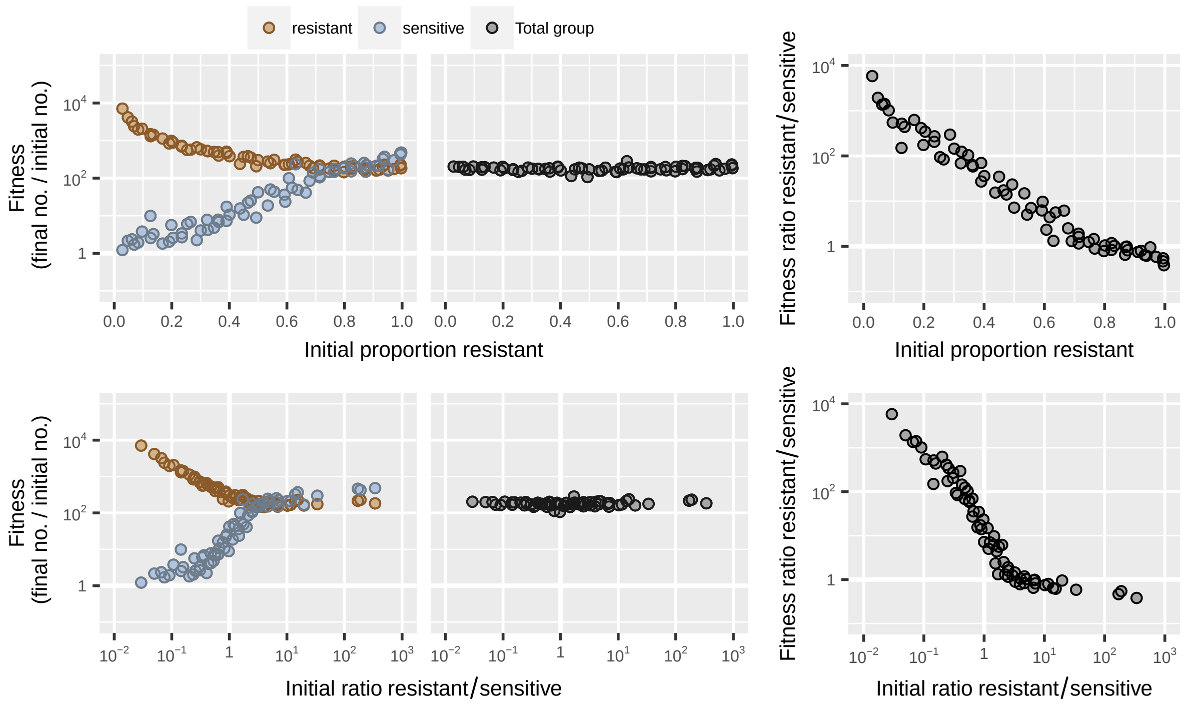

**Figure S73:** Dataset from Yurtsev *et al.* [19] showing coculture of antibiotic-resistant and sensitive *Escherichia coli* in media containing 100 $\mu$g/ml ampicillin with 200-fold dilution.

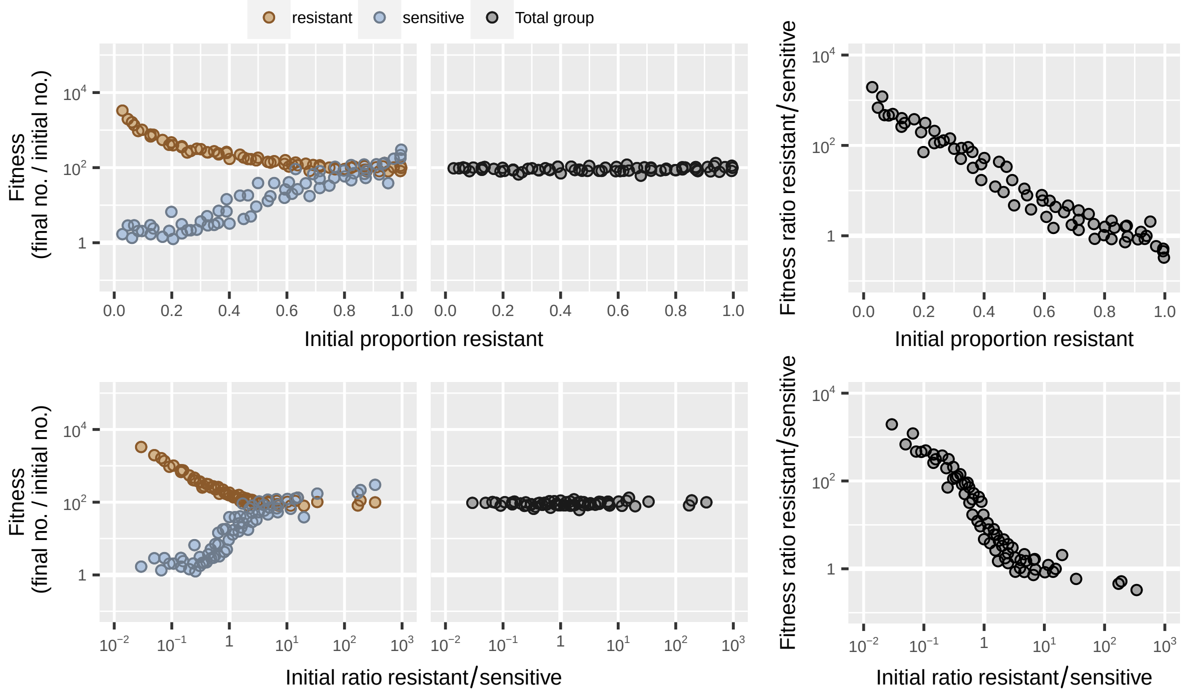

**Figure S74:** Dataset from Yurtsev *et al.* [19] showing coculture of antibiotic-resistant and sensitive *Escherichia coli* in media containing 200 $\mu$g/ml ampicillin with 100-fold dilution.

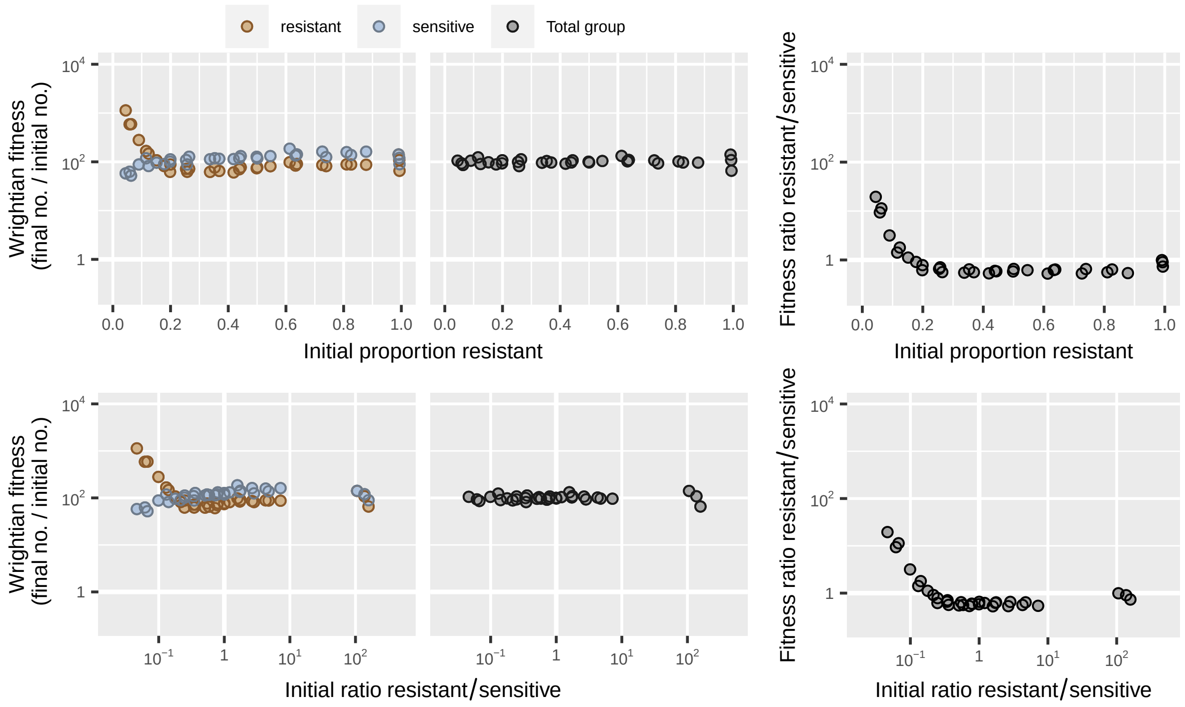

**Figure S75:** Dataset from Yurtsev *et al.* [19] showing coculture of antibiotic-resistant and sensitive *Escherichia coli* in media containing 2 $\mu$g/ml ampicillin and 41 ng/ml tazobactam inhibitor.

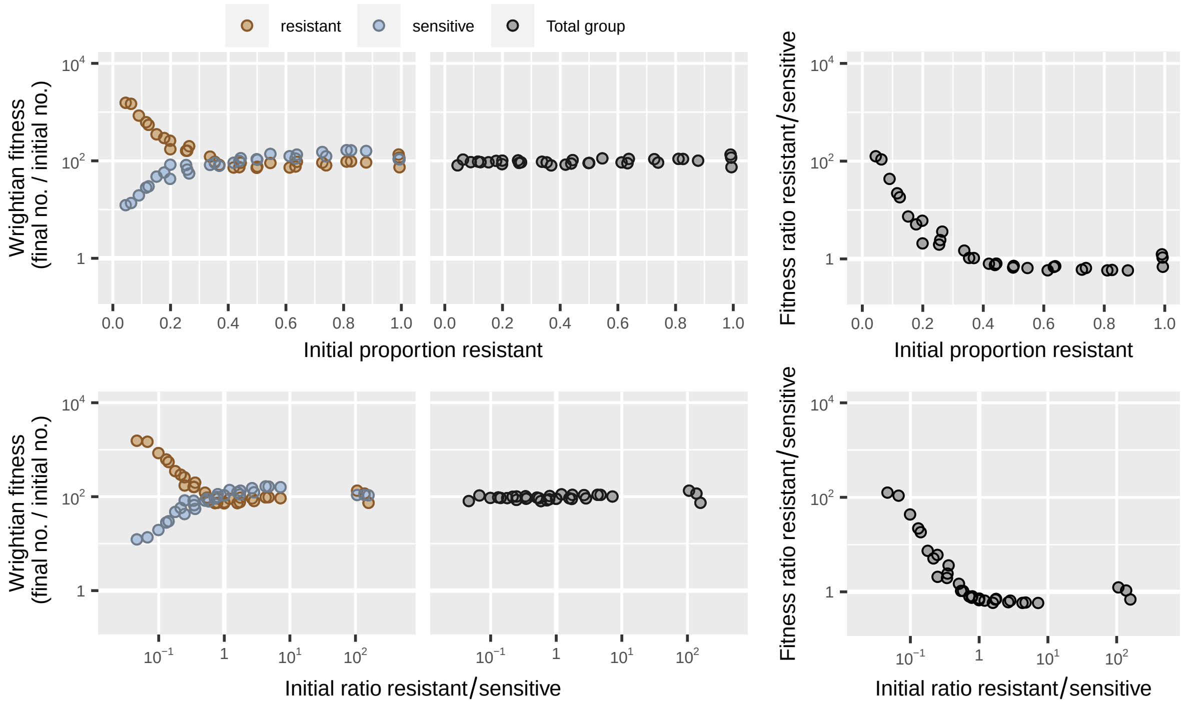

**Figure S76:** Dataset from Yurtsev *et al.* [19] showing coculture of antibiotic-resistant and sensitive *Escherichia coli* in media containing 2 $\mu$g/ml ampicillin and 118 ng/ml tazobactam inhibitor.

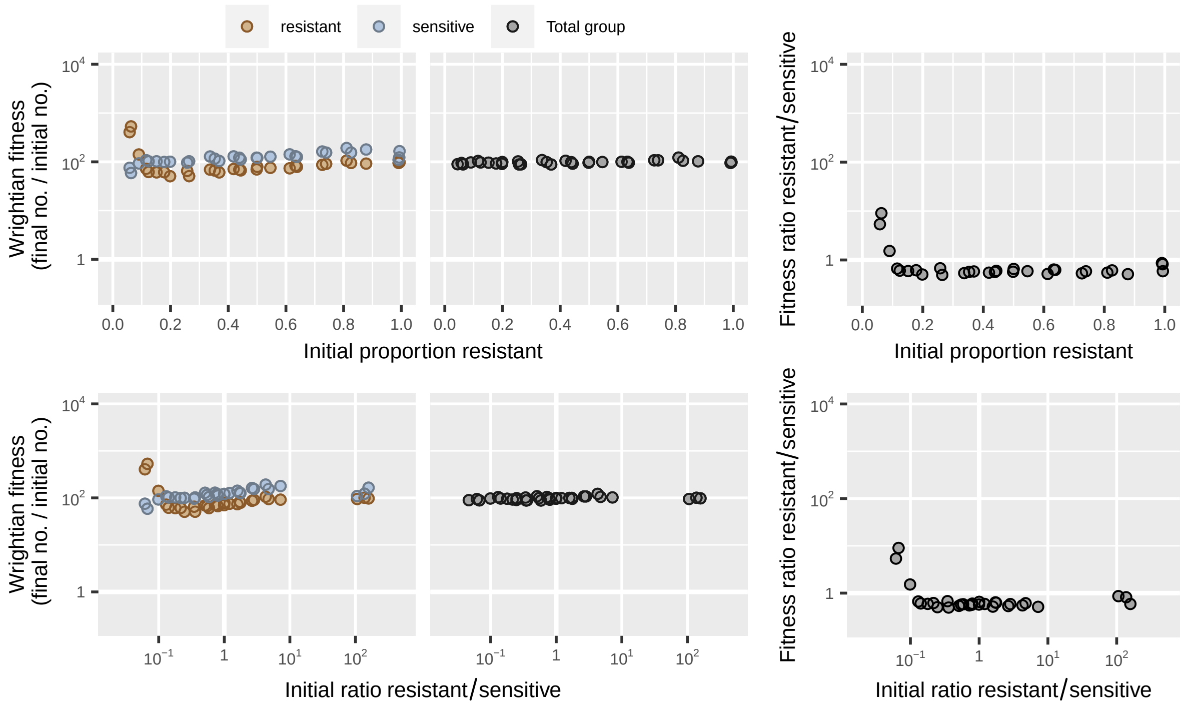

**Figure S77:** Dataset from Yurtsev *et al.* [19] showing coculture of antibiotic-resistant and sensitive *Escherichia coli* in media containing 5 μg/ml ampicillin and 8 ng/ml tazobactam inhibitor.

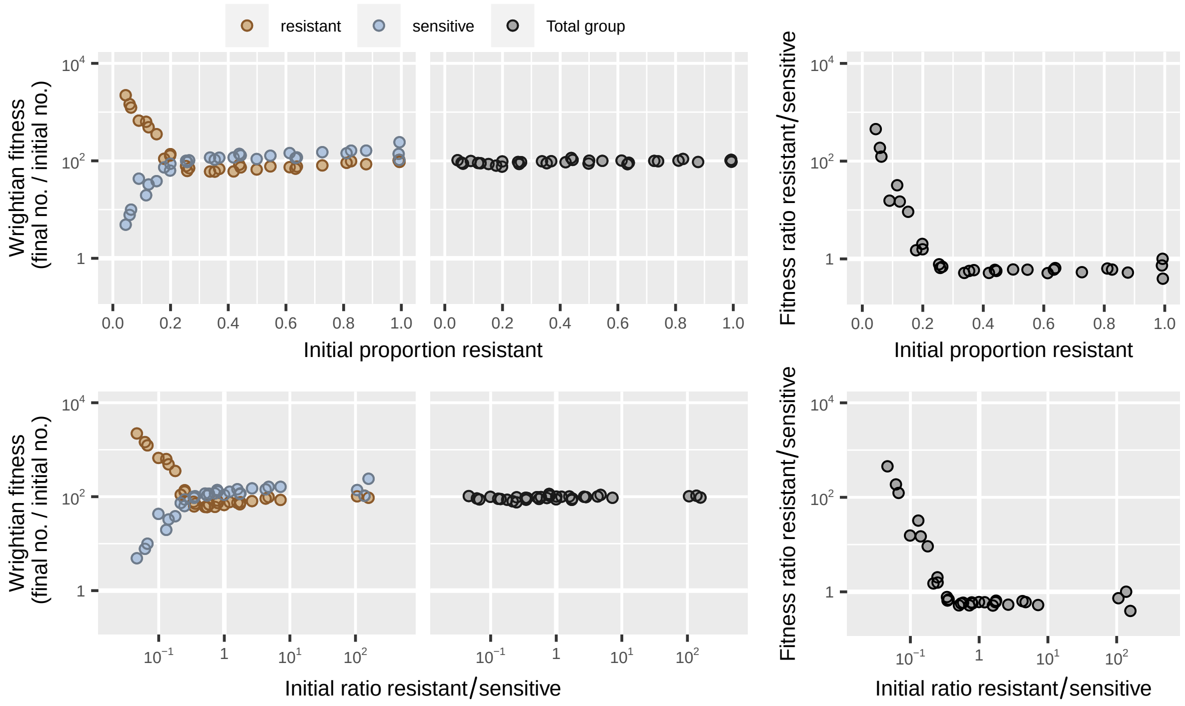

**Figure S78:** Dataset from Yurtsev *et al.* [19] showing coculture of antibiotic-resistant and sensitive *Escherichia coli* in media containing 20 μg/ml ampicillin and 8 ng/ml tazobactam inhibitor.

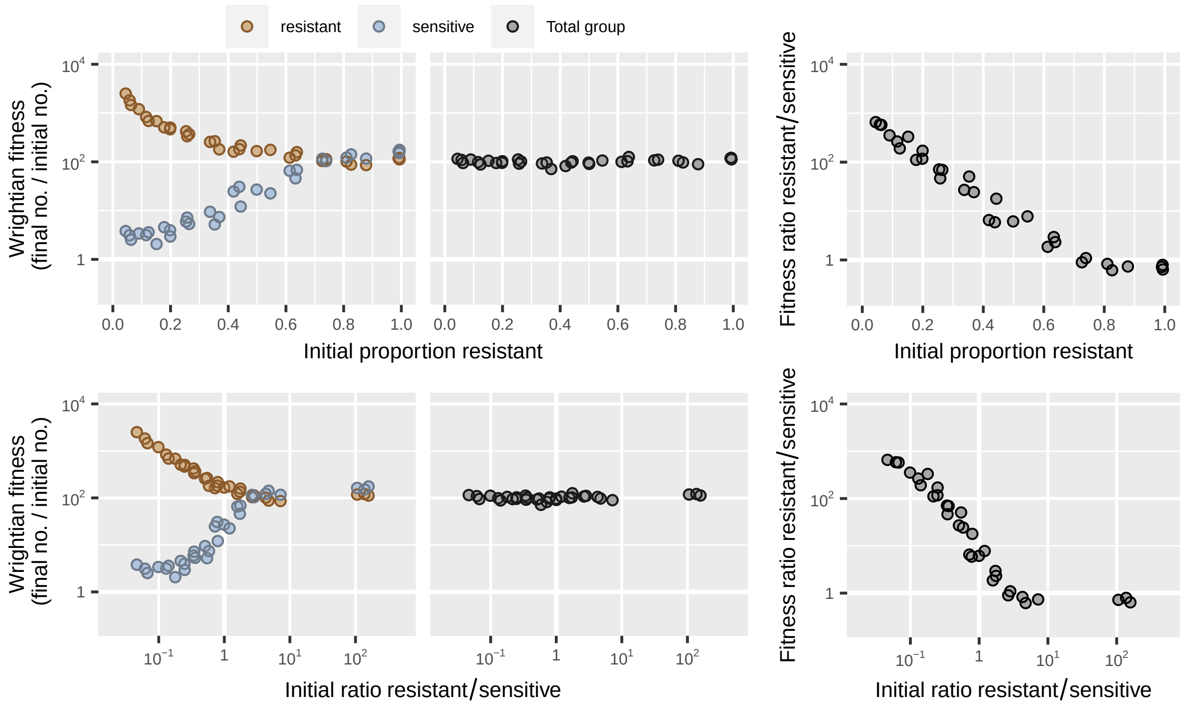

**Figure S79:** Dataset from Yurtsev *et al.* [19] showing coculture of antibiotic-resistant and sensitive *Escherichia coli* in media containing 20 $\mu$g/ml ampicillin and 41 ng/ml tazobactam inhibitor.

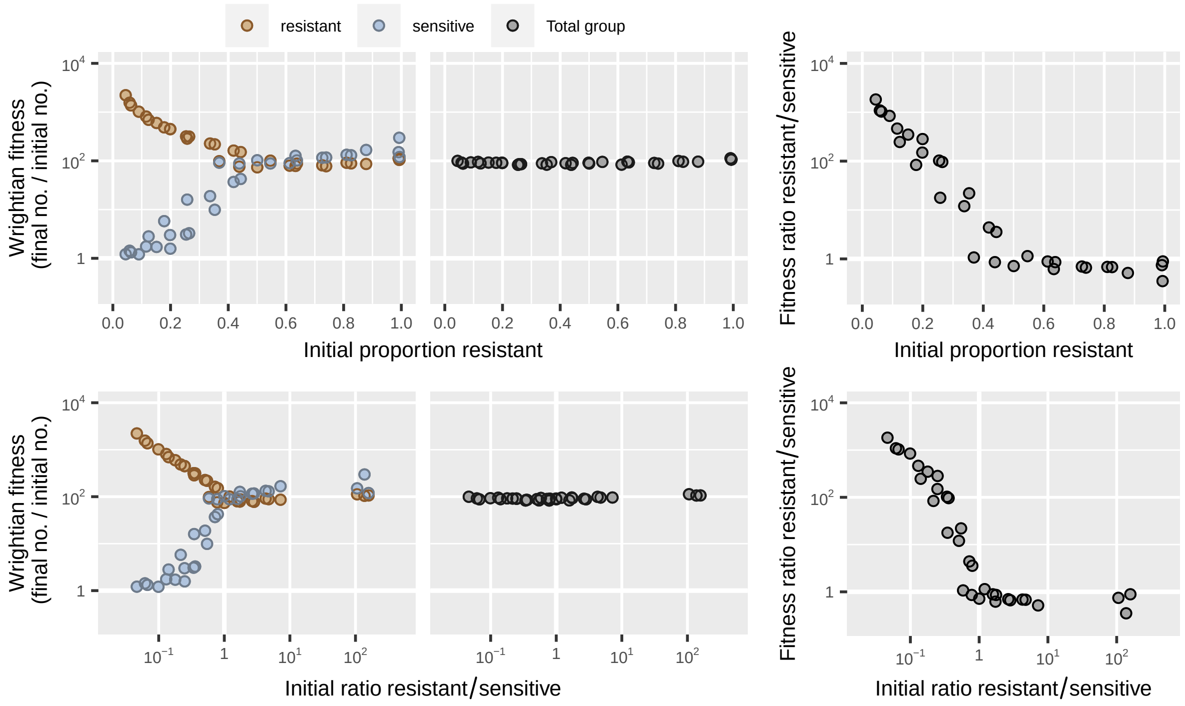

**Figure S80:** Dataset from Yurtsev *et al.* [19] showing coculture of antibiotic-resistant and sensitive *Escherichia coli* in media containing 100 $\mu$g/ml ampicillin and 8 ng/ml tazobactam inhibitor.

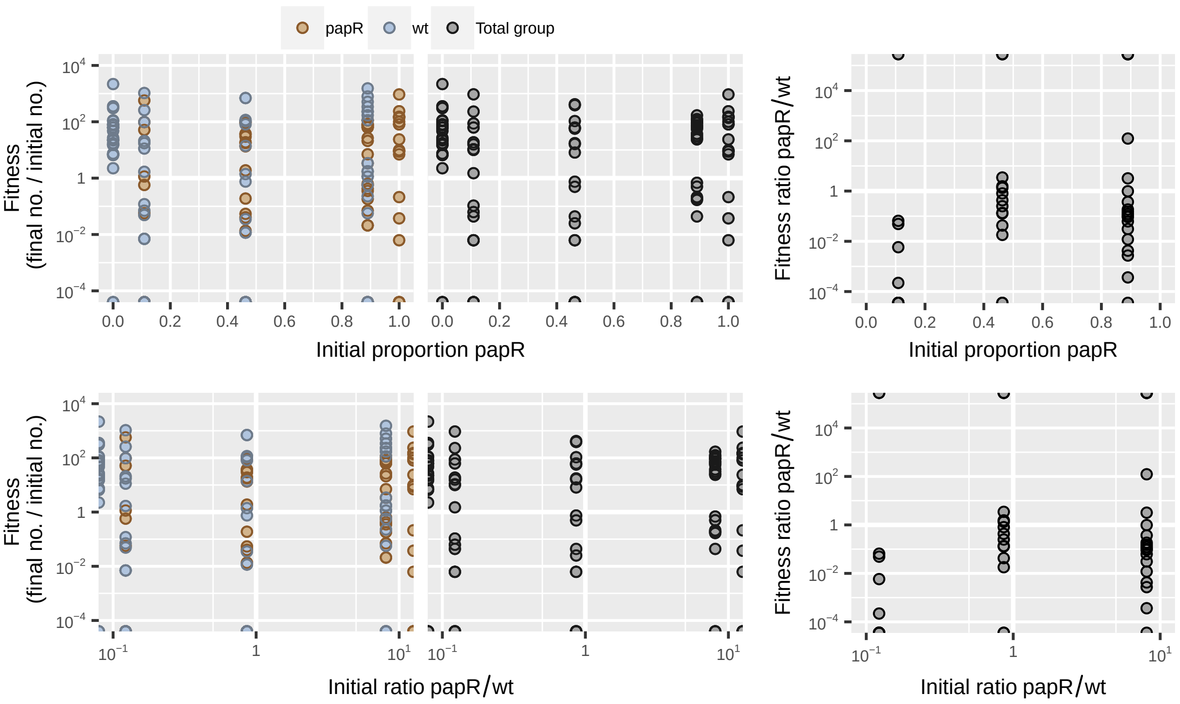

**Figure S81:** Dataset from Zhou *et al.* [20] showing coinfection of caterpillars with wild-type *Bacillus thuringensis* and a $\Delta papR$ signal-null quorum sensing mutant at $8 \times 10^3$ initial spore dose.

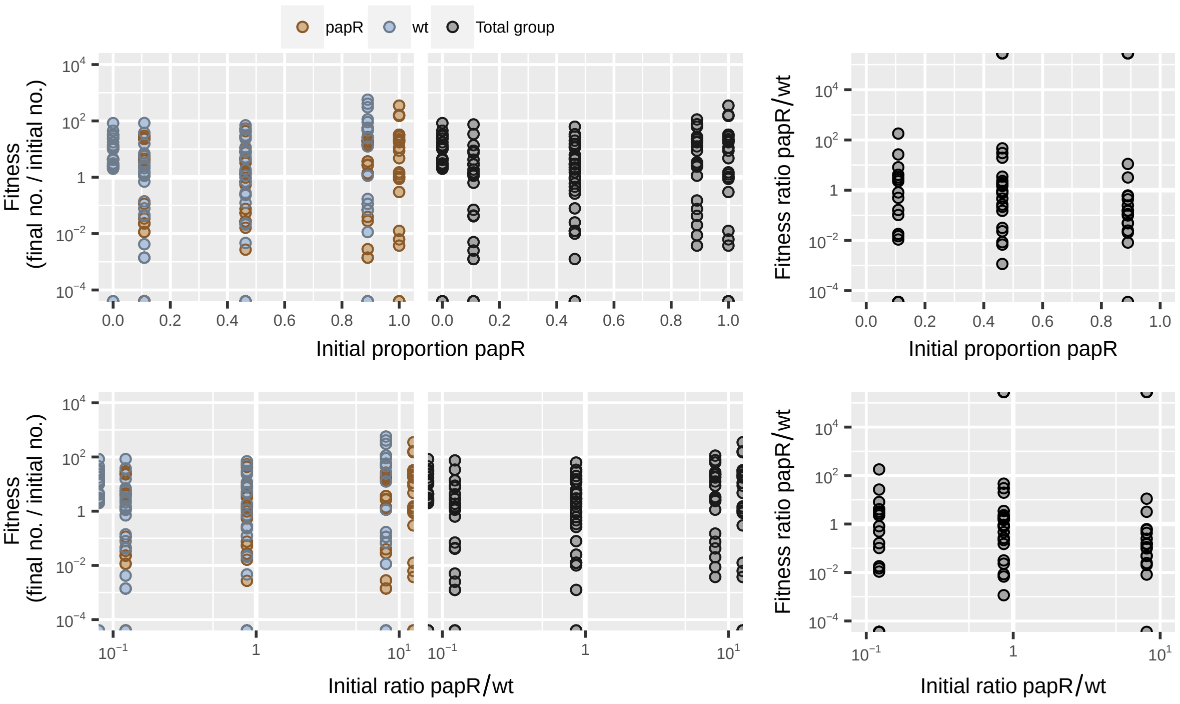

**Figure S82:** Dataset from Zhou *et al.* [20] showing coinfection of caterpillars with wild-type *Bacillus thuringensis* and a $\Delta papR$ signal-null quorum sensing mutant at $4 \times 10^4$ initial spore dose.

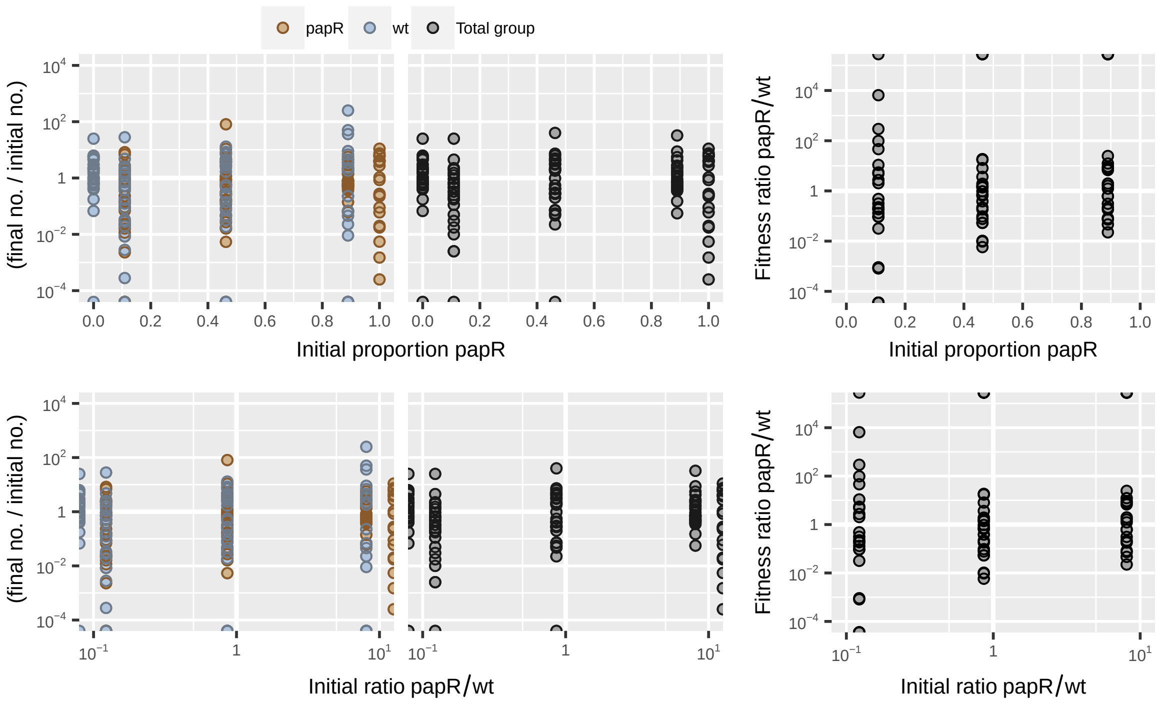

**Figure S83:** Dataset from Zhou *et al.* [20] showing coinfection of caterpillars with wild-type *Bacillus thuringensis* and a $\Delta papR$ signal-null quorum sensing mutant at $2 \times 10^5$ initial spore dose.

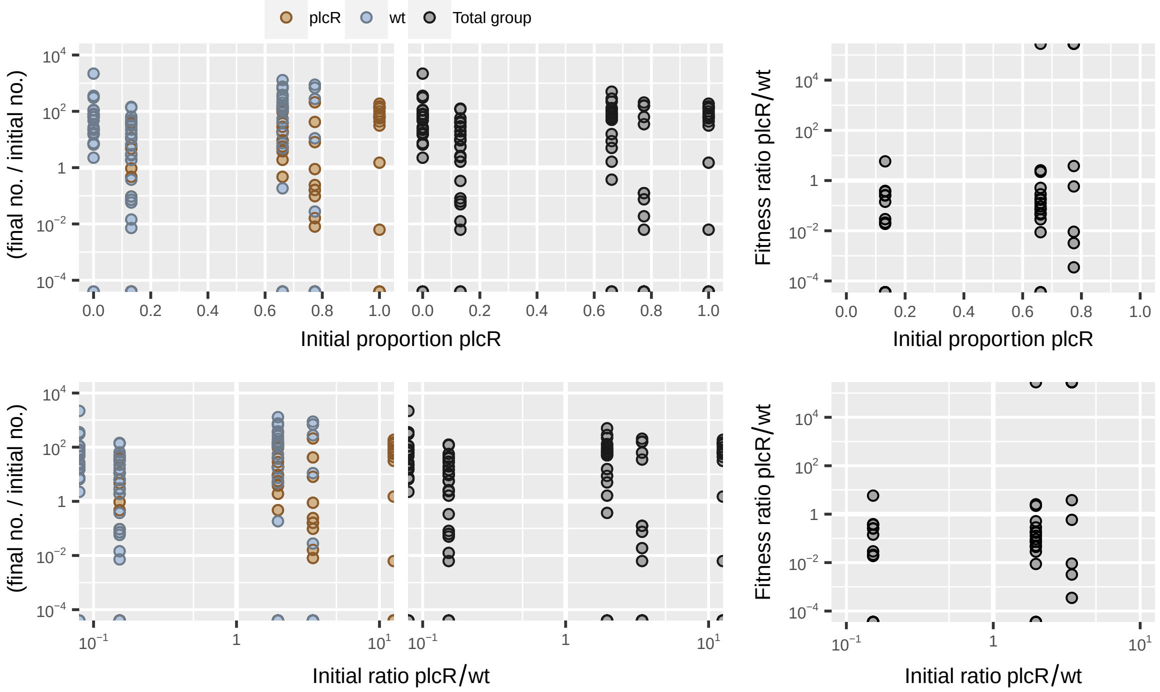

**Figure S84:** Dataset from Zhou *et al.* [20] showing coinfection of caterpillars with wild-type *Bacillus thuringensis* and a $\Delta plcR$ signal-blind quorum sensing mutant at $8 \times 10^3$ initial spore dose.

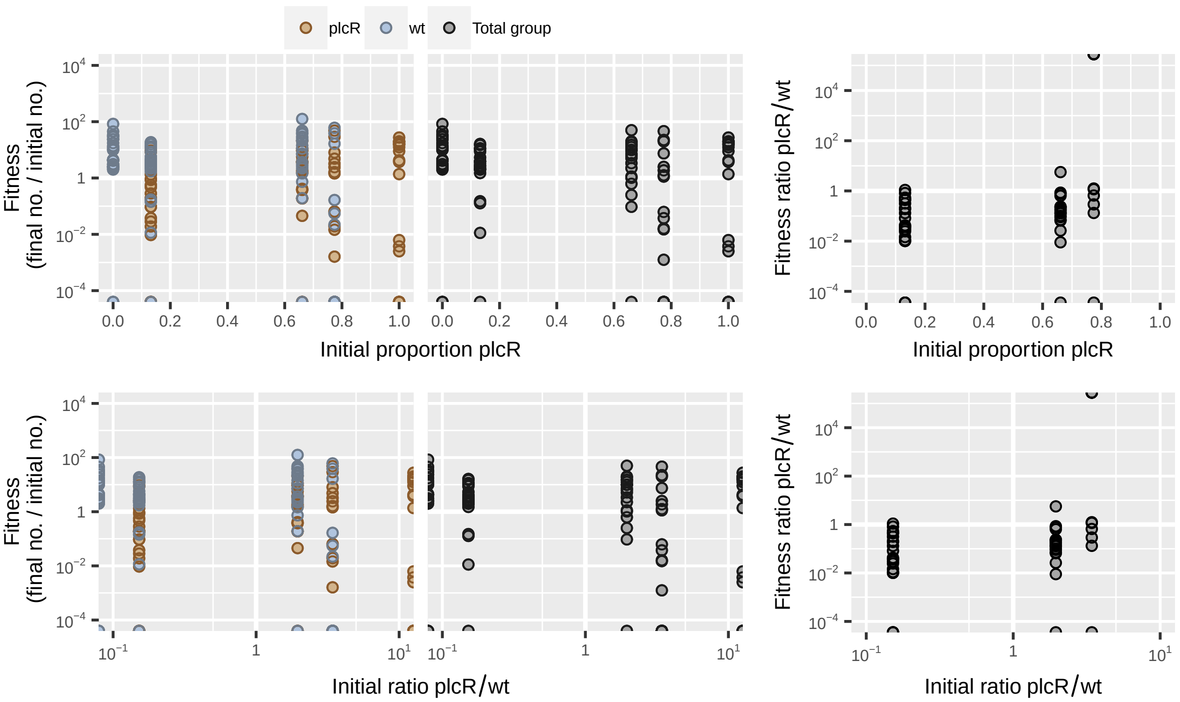

**Figure S85:** Dataset from Zhou *et al.* [20] showing coinfection of caterpillars with wild-type *Bacillus thuringensis* and a $\Delta plcR$ signal-blind quorum sensing mutant at $4 \times 10^4$ initial spore dose.

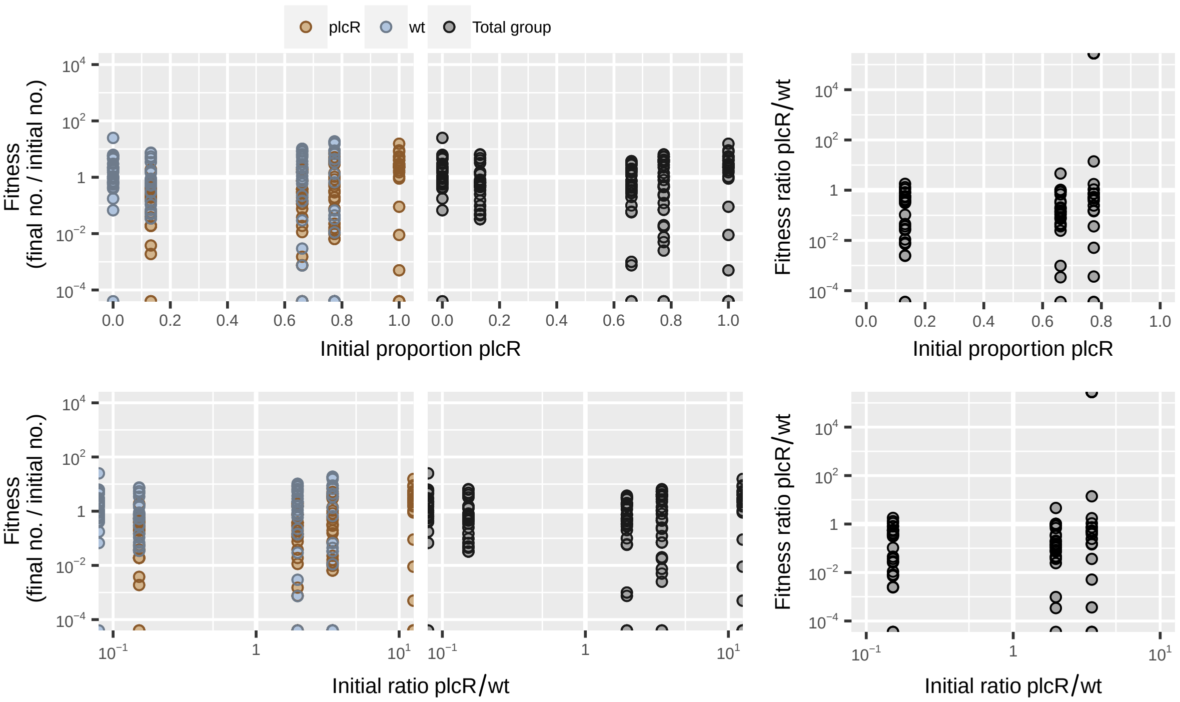

**Figure S86:** Dataset from Zhou *et al.* [20] showing coinfection of caterpillars with wild-type *Bacillus thuringensis* and a $\Delta plcR$ signal-blind quorum sensing mutant at $2 \times 10^5$ initial spore dose.

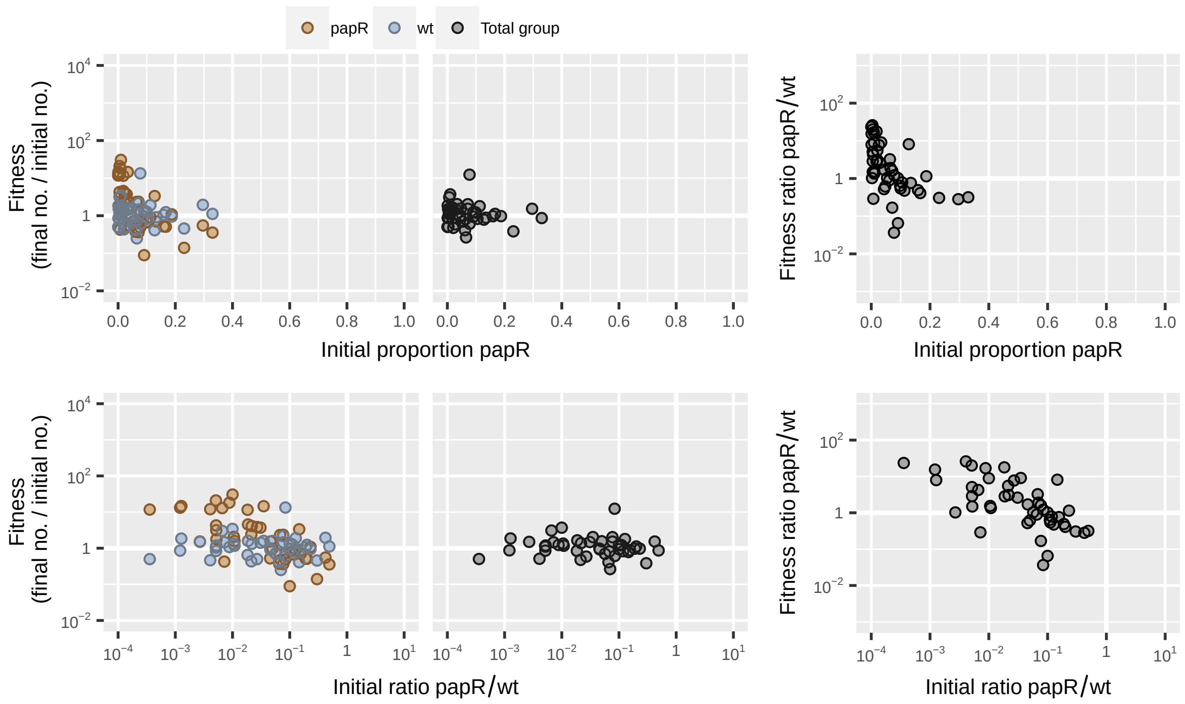

**Figure S87:** Dataset from Zhou *et al.* [20] showing fitness of wild-type *Bacillus thuringensis* and a Δ*papR* signal-null quorum sensing mutant in homogenized caterpillar cadavers.

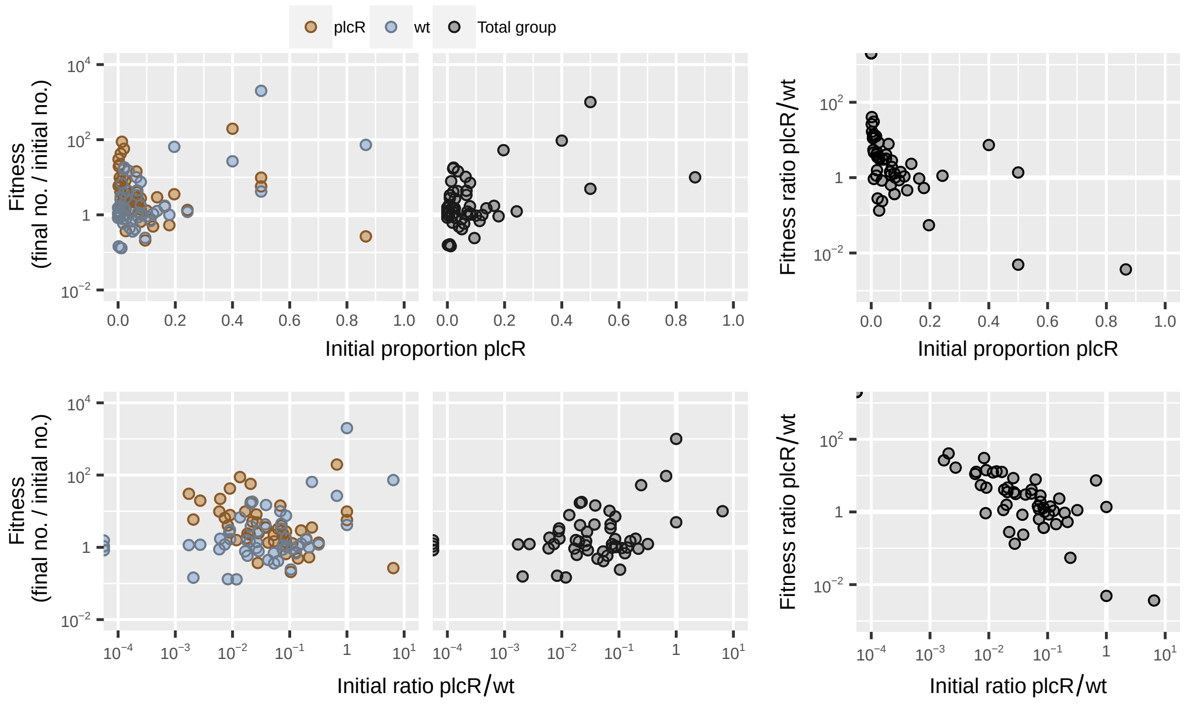

**Figure S88:** Dataset from Zhou *et al.* [20] showing fitness of wild-type *Bacillus thuringensis* and a Δ*plcR* signal-blind quorum sensing mutant in homogenized caterpillar cadavers.
